# Multispot single-molecule FRET: high-throughput analysis of freely diffusing molecules

**DOI:** 10.1101/085027

**Authors:** A. Ingargiola, E. Lerner, S. Chung, F. Panzeri, A. Gulinatti, I. Rech, M. Ghioni, S. Weiss, X. Michalet

## Abstract

We describe an 8-spot confocal setup for high-throughput smFRET assays and illustrate its performance with two characteristic experiments. First, measurements on a series of freely diffusing doubly-labeled dsDNA samples allow us to demonstrate that data acquired in multiple spots in parallel can be properly corrected and result in measured sample characteristics identical to those obtained with a standard single-spot setup. We then take advantage of the higher throughput provided by parallel acquisition to address an outstanding question about the kinetics of the initial steps of bacterial RNA transcription. Our real-time kinetic analysis of promoter escape by bacterial RNA polymerase confirms results obtained by a more indirect route, shedding additional light on the initial steps of transcription.

Finally, we discuss the advantages of our multispot setup, while pointing potential limitations of the current single laser excitation design, as well as analysis challenges and their solutions.

## 1 Introduction

### 1.1 Background

Freely-diffusing single-molecule FRET (smFRET) studies have yielded a wealth of new scientific results since their proof-of-principle demonstration almost two decades ago [1, 2]. While continuous improvements in fluorophores, labeling chemistry, excitation optics, theoretical analysis and combination with other techniques have increased its performance, there are still obstacles to the wide adoption of this powerful technology, chief among which is the issue of throughput.

Indeed, in order to ensure the separate detection of fast diffusing individual molecules, low concentrations, small excitation volumes and high excitation intensities are necessary. This is generally obtained with a so-called confocal geometry, where a laser is focused into a single diffraction-limited volume of the sample and a single-pixel detector collects light only from this volume. The low concentration needed to minimize the occurrence of simultaneous crossing of two or more molecules (<100 pM), lead to infrequent single-molecule transits so that several minutes, and in some extreme cases hours, are needed to accumulate a statistically significant number of transit events. These characteristics have thus far confined the technique to equilibrium measurements or very slow kinetics studies, with time scales on the order of several minutes [3].

Parallelizing measurements provides the necessary increase in throughput required for freely-diffusing fluorescence experiments. Rather than observing individual molecules sequentially in a single spot, different molecules are observed simultaneously at several independent locations within the sample. Camera-based approaches to smFRET [4] have a temporal resolution limited to a few ms, confining this approach to freely-diffusing molecules enclosed in small static volumes such as zero-mode waveguides [5], microscopically patterned PDMS wells [6], or tethered vesicles [7, 8]. Another, more complex approach for increased throughput involves extending the single-spot confocal geometry to a multispot confocal geometry. However, this requires parallel photon-counting capabilities, which have only become available recently with the development of single-photon avalanche diode (SPAD) arrays.

### 1.2 SPAD Arrays

Recently, we described such a confocal multispot setup using a linear array of 8 SPADs for fluorescence correlation spectroscopy (FCS) analysis of single fluorescent dyes [9]. Our group and others have since demonstrated similar achievements with much larger SPAD arrays based on CMOS technology [10, 11]. These studies were however limited to working with labeled molecules at concentrations larger than the single-molecule regime (> 1 nM), mostly because of the limited detection efficiency and large dark count rates of some of these CMOS SPAD arrays.

The SPAD array used in our original FCS study was fabricated using a custom silicon technology developed by the Politecnico di Milano (POLIMI) group [12]. This dedicated technology results in better detection efficiency than CMOS SPAD arrays in the visible range and allowed us to demonstrate single-molecule sensitivity [9]. In follow-up experiments, we further demonstrated two-color smFRET measurements. We first used a single 8-SPAD array divided into two spectral detection channels each with 4 SPADs [13], and later extended this work to two arrays of 8 SPADs [14]. More recently developed SPAD arrays comprising even more SPADs open up the perspective of significantly higher throughput measurements [15, 16].

### 1.3 Multispot smFRET

Although the principle of multispot smFRET as described above is conceptually simple, its proper execution requires a number of careful checks. In particular, even though a single optical setup is used, each excitation spot and its corresponding detection volume, as well as associated SPADs, have unique characteristics, making this single experiment in effect a series of *N* parallel but slightly different experiments. The only true common denominator to all these parallel experiments is the sample itself. Any other parameter such as laser excitation power, excitation and detection point-spread functions (PSFs), photon detection efficiency (PDE), etc., cannot be assumed to be exactly identical from one spot to the next. Therefore, in order to be able to compare or pool data originating from different spots, a number of calibration and correction factors need to be determined and a robust data analysis approach needs to be designed and validated.

In other words, using a multispot setup to perform smFRET experiments forced us to look into a rigorous way to compare data about a given single-molecule system, acquired in potentially very different conditions. This effort should therefore hopefully be of general interest to the single-molecule fluorescence community, not only when dealing with multispot arrangements, but also when trying to compare results obtained from single-spot experiments performed on different setups.

This paper is organized as follows. We first briefly introduce the two experimental setups used in this work, a “standard” single-spot two-alternating lasers (µs-ALEX) setup used as reference, and a single-laser, two-color 8-spot setup whose detector characteristics are discussed (Section 2). Section 3 provides an overview of the smFRET analysis workflow used in this study, most of the details being presented in different appendices in the Supporting Information. Section 4 compares the smFRET results obtained for a series of dsDNA samples with both single-spot µs-ALEX setup (our gold standard) and multispot setup, thus validating our method. To showcase the advantage of the higher throughput of the multispot setup, Section 5 studies the escape kinetics of *Escherichia coli* (*E*. coli) RNA polymerase from a gene promoter region during DNA transcription initiation. We conclude with a brief overview of the main developments presented in this work, as well as an outlook on future improvements (Section 6). Samples and setup descriptions, extended discussion and analysis methods are available in Supporting Information.

### 1.4 Data and Software Availability

All datasets discussed in this paper (for both single and multispot experiments) have been uploaded on Figshare and are thus citable (references can be found in SI-Appendix 1). Datasets are stored in the Photon-HDF5 file format, an open multi-platform file format for timestamp-based single-molecule fluorescence experiments [17].

Data analysis was performed with our open source FRETBursts Python software [18] and with our ALiX LabVIEW software available as a standalone executable, with some fits performed with Origin 9.1 (OriginLab). To ensure computational reproducibility, we did our best efforts to document and report all analysis details. Analysis performed with FRETBursts can be replicated by running the main Jupyter notebooks associated with this paper, linked to in SI-Appendix 2. Analysis performed with ALiX can be reproduced using scripts linked to in SI-Appendix 2.

## 2 Setup Description

Two setups were used in this work. A “classic” single-spot setup (described in ref. [19] and in SI-Appendix 3) designed around an inverted microscope was used for single-spot µs-ALEX measurements (Fig. SI-1). The setup used for multispot smFRET measurements was designed around an identical microscope but used different lasers, optics and detectors, (described in ref. [14] and SI-Appendix 3, Fig. SI-3). A brief description of the multispot detector characteristics is provided below.

The two SPAD arrays used in this study were similar to the one described in ref. [9]. Four main characteristics are of interest for this work: detection efficiency, dark count rate, afterpulsing probability and crosstalk probability.

### 2.1 Detection Efficiency

The SPAD arrays fabricated using the standard custom technology developed by POLIMI are characterized by a similar PDE as that of the single SPAD module commercialized by Micro Photon Devices (Fig. SI-2) [12, 20, 21] and used for donor (D) detection in the single-spot µs-ALEX setup. In particular, their photon detection efficiency in the acceptor (A) detection range (red area in Fig. SI-2) is about half of that of the red-enhanced SPAD used for acceptor detection in the single-spot µs-ALEX setup. These characteristics result in a noticeable difference in correction factors between the two types of measurements, as discussed later in the text.

### 2.2 Dark Count Rates

Dark counts are random and due to thermally excited charge carriers in the detection area, leading to an avalanche. Dark count rates (DCRs) measured for the two 8-SPAD arrays are reported in Table 1. The range of values is widespread, with more than two orders of magnitude separating the largest and smallest dark count rates. In particular, half the SPADs used for the acceptor channel have a DCR comparable to or larger than the typical sample background rate (from 1 to 3 kHz depending on sample and spot). The influence of these large DCR values for analysis will be addressed later in the text.

**Table 1:** Dark count rates. Dark count rates (DCR, in Hz) for individual SPADs of the two SPAD arrays used in this work. D-channel (resp. A-channel) indicates the 8-SPAD array used to collect the donor (resp. acceptor) signal from each spot. The smallest (24 Hz) and largest (7 kHz) DCR are highlighted. The top row indicates the pixel number in each array.

|  | 1 | 2 | 3 | 4 | 5 | 6 | 7 | 8 |
| --- | --- | --- | --- | --- | --- | --- | --- | --- |
| D-channel | 36 | 117 | 3,775 | 567 | 29 | 443 | 27 | 24 |
| A-channel | 3,679 | 7,207 | 78 | 901 | 1,717 | 4,321 | 49 | 53 |

### 2.3 Afterpulsing Probability

Afterpulses are the result of trapped carriers generated during an avalanche and released shortly after it. The integrated afterpulsing probability of each SPAD was measured under moderate constant illumination according to the simple relation [22]:

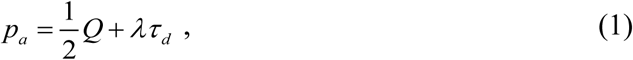

where *Q* = *Var(S)/<S>* - 1 is the Mandel parameter computed for the measured detector signal *S*, *λ* is the incident count rate and *τ*_*d*_ = 120 ns is the detector deadtime. *Q* depends in general on the time bin *T* used to record signal *S*, but for a moderate constant illumination(*λ τ*_*d*_ << 1), this dependence is negligible and the afterpulsing probability is simply half the Mandel parameter. The resulting afterpulsing probabilities (in percent) are reported in Table 2.

**Table 2:** Afterpulsing probabilities. Afterpulsing probabilities of individual SPADs (in percent) in the two SPAD arrays. The top row indicates the pixel number in each array.

|  | 1 | 2 | 3 | 4 | 5 | 6 | 7 | 8 |
| --- | --- | --- | --- | --- | --- | --- | --- | --- |
| D-Channel | 1.4 | 1.3 | 1.8 | 1.2 | 1.4 | 2.0 | 1.7 | 1.0 |
| A-Channel | 4.7 | 5.6 | 9.8 | 3.5 | 4.8 | 3.9 | 4.4 | 4.7 |

They are larger than for single-pixel SPADs, which have typically *p_a_* < 0.1 %, but, when properly accounted for by the different correction factors discussed later in the text, do not affect the results.

### 2.4 Crosstalk

In SPAD arrays, an additional source of noise originates from crosstalk between pixels. Crosstalk can be due to electrical interference between pixels (electrical crosstalk), secondary photons emitted by SPADs during the avalanche process [23] or emission point-spread-function (PSF) spillover into more than one SPAD. With a proper electrical design, the first can be made negligible, while the latter was excluded by a separation between SPADs one order of magnitude larger than the PSF extension. Optical crosstalk, on the other hand, can be minimized but not completely eliminated. We measured the optical crosstalk in the two SPAD arrays used in the multispot setup and found it to be smaller than 0.5% for all pixel pairs (see SI-Appendix 4), resulting in negligible effects on burst analysis.

## 3 smFRET Data Analysis

smFRET data analysis of freely-diffusing single-molecule involves a number of steps which have been described with some details in the literature [18, 24–28]. However, our experience is that data analysis reproducibility is oftentimes hampered not only by the lack of raw data files, but also by the absence of values for specific parameters and algorithms implementation details. Here, we strived to provide as much information as possible in the main text and Supporting Information, complemented by algorithmic implementation when feasible. We found that analysis of multispot data can be made relatively straightforward, but requires some care and cannot be oversimplified.

For the interest of legibility, the main text only introduces concepts and notations, details being provided in the Supporting Information.

### 3.1 Photon Streams Definition

In μs-ALEX measurements, 4 streams of photon timestamps (or photon streams) can be distinguished, functions of the excitation period during which the timestamps are recorded and their detection channel [29, 30]. We will denote them as D_ex_D_em_, D_ex_A_em_, A_ex_A_em_ and A_ex_D_em_. These streams identify photons detected during the D- or A-excitation period (D_ex_ or A_ex_) and by the D- or A-detection channels (D_em_ or A_em_).

By contrast, for single-laser smFRET measurements, such as done with the multispot setup, there is only one excitation period (the donor period) and therefore only two photon streams: D_ex_D_em_ and D_ex_A_em_. We will denote them as D_em_ and A_em_ for brevity.

Details on how photon streams are defined based on timestamp information can be found in SI-Appendix 5.

### 3.2 Background Rate Estimation

Background rates, noted *b_stream_*, where the photon stream are defined in the previous section (e.g. *b_DexDem_*, *b_DexAem_*, etc.), were measured for each photon stream. As discussed in SI-Appendix 6, different physical mechanisms contributing uncorrelated counts give rise to an exponential tail in the distribution of inter-photon delays [31]. At short delays, the inter-photon delay distribution departs from a pure exponential due to the contribution of single-molecule bursts [31]. Defining a cut-off time *τ*_*min*_ as the delay above which the distribution is considered exponential, the count rate corresponding to the exponential tail can be simply calculated by the maximum likelihood estimator (MLE):

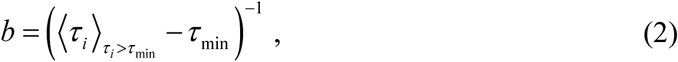

where the average is taken over all delays *τ_i_* > *τ_min_* [18]. Choosing τ_*min*_ involves some tradeoff, but can be automated (SI-Appendix 6). In addition, since background rates can occasionally vary during the measurement, we estimated background rates over consecutive time windows of, typically, 30 s.

For the multispot measurements, background rates for each of the two available streams (D_em_ and A_em_) were estimated separately for each spot.

### 3.3 Burst Search

The burst search algorithm used in this work is based on the sliding window algorithm introduced by the Seidel group [24, 32] (see also [18]). Burst search can be specified by three parameters: type of photon stream, number *m* of consecutive photons used to compute the local count rate (typically, *m* = 10), and count rate threshold *r_min_*. This threshold can be defined in different ways. When defined as a multiple of the background rate (*r*_*min*_= *Fb_stream_*), it will follow background variations during the measurement and ensure that a minimal burst signal-to-background ratio (SBR) is achieved (minimum SBR burst search [33, 34]). Alternatively, the threshold can be set to a fixed value (minimum count rate burst search). The choice between these two approaches is dictated by the specific aim of the analysis, as will be discussed later.

A detailed description of various burst searches and their influence on different observables can be found in SI-Appendix 7.

### 3.4 Burst Selection Criterion

The outcome of a burst search as just described is in general a large set of bursts, the majority of which are comprised of very few photons (Fig. SI-14A&B). Because *m* photons are used for the search, the minimum burst size before background correction is *m*, which in general is a small number (*e.g. m* = 10). These small bursts contribute very little useful information. Moreover, they tend to increase the variance of observables such as the proximity ratio (*PR*) distribution, as discussed below and in SI-Appendix 8. This makes the separation of populations characterized by similar proximity ratios, such as a low FRET population and D-only population, more difficult in the absence of laser alternation. For these reasons, rejecting bursts whose size is smaller than a preset minimum value *Smin* is a natural step in all analyses, although other criteria can be used and combined for specific purposes (e.g. minimum burst duration, minimum count rate, etc.). Further discussion of this burst selection step and the influence of parameter *S_min_* on analysis results can be found in SI-Appendix 8.

### 3.5 Proximity Ratio and FRET Efficiency Analysis

smFRET studies only aiming to distinguish between different populations of molecules on the basis of their FRET efficiencies *E* do not necessarily require the precise calculation of *E*. The proximity ratio *PR* can be used instead, which involves only background corrected quantities:

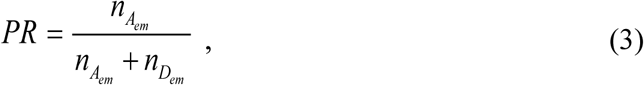

where *n*_*D*_*em*__ and *n*_*A*_*em*__ are the background-corrected donor and acceptor signals (upon donor excitation) respectively. These quantities depend on the different correction factors discussed next, and therefore, any two measurements of the same sample performed on different setups may results in different values of *PR*. For an exact comparison between these measurements, corrected quantities (*i.e.* the FRET efficiency *E*) need to be computed. The relation between proximity ratio, FRET efficiency and those correction factors has been described previously [30] and is reminded in SI-Appendix 9.

Similarly, in a multispot setup, where correction factors may be spot-dependent, computation of correction factors first requires calculation of proximity ratio values, and comparison of results from different spots requires calculation of corrected FRET efficiency values.

Statistical analysis of the proximity ratio and FRET efficiency was performed by 3 different methods: Gaussian fit of their histogram, kernel density estimation (KDE) and shot noise analysis (SNA), as discussed in SI-Appendix 10 & 11.

### 3.6 Correction Factors

Several correction factors are necessary to convert raw photon count quantities into physical observables such as FRET efficiency. These parameters are:

- the *donor leakage* factor *l*, quantifying the donor signal detected in the acceptor channel as a fraction of the donor channel signal,
- the *direct acceptor excitation* factor *d*, quantifying the amount of acceptor signal due to direct excitation by the donor excitation laser,
- the fluorescence quantum yield and detection efficiency correction factor *γ*, accounting for the different brightness of the donor and acceptor dye due to their different fluorescence quantum yield and different detection efficiency in their respective channels.

SI-Appendix 9 provides details on how these correction factors were estimated in the single-spot μs-ALEX and multispot experiments respectively. Here, we briefly summarize the differences between the two types of experiments.

In single-spot μs-ALEX experiments, it is possible to estimate the donor leakage and direct acceptor excitation factors in each sample, as long as there are enough donor-only (DO) and acceptor-only (AO) bursts, respectively. Indeed, each population (DO, AO and donor and acceptor-labeled molecules, DA) can be readily isolated in the so-called two-dimensional ALEX histogram representing the stoichiometry ratio (*SR*) of each burst as a function of its proximity ratio (*PR*) (SI-Appendix 9). Factor *l* can be computed from the DO *PR* histogram, while factor *d* can be computed from the AO *SR* histogram. These coefficients should not depend on the sample (provided the dyes are in similar chemical environments) or the specific measurement (as long as nothing has changed in the setup), but it is always useful to be able to confirm it. Average factors were used in the final analysis. In general, it is recommended to use dedicated DO and AO samples to measure these correction factors.

The single-spot μs-ALEX *γ* factor was estimated according to Lee *et al.* [30] with the series of 5 dsDNA samples as described in SI-Appendix 9.

In multispot experiments, the absence of acceptor laser excitation prevents classification of single-molecule bursts based on the stoichiometry ratio, leaving the proximity ratio as the only observable to separate between DO and DA bursts (AO bursts are in general not detected). While *PR* is sufficient to distinguish between high FRET and DO bursts, the overlap between *PR* distribution of both types of molecules increases as the FRET value in the sample decreases, which makes it difficult if not impossible to separate DO and DA populations in low FRET samples. For this reason, a DO sample was used in addition to the highest FRET samples of the dsDNA series for extracting the *l* factor.

It is possible to correct for the acceptor direct-excitation contribution (*Dir*) in multispot measurements employing a single laser excitation (SI-Appendix 9, Eqs. (SI.53)-(SI.56)), if the direct excitation coefficient *d_T_* = σ_*A*_/σ_*D*_ (ratio of acceptor and donor absorption cross-sections at the donor excitation wavelength) is known. This ratio, being a molecular characteristics, is independent from the setup or experiment used to compute it. We therefore estimated *d_T_* using a single-spot μs-ALEX measurement. Finally, the *γ* factor for the multispot system was estimated from the corrected *E* obtained from μs-ALEX measurements of one sample (12d) used as reference (see SI-Appendix 9 for details).

### 3.7 Fluorescence Correlation Spectroscopy

FCS analysis was used to characterize the excitation/detection volumes of the donor and acceptor channel for each spot of the multispot measurements. The analysis is relatively straightforward in this situation where a single sample, characterized by a unique diffusion coefficient, is observed at different locations. Any difference in diffusion time can be therefore interpreted as due to differences in excitation/detection volume. The results were compared to those obtained in the single-spot µs-ALEX measurements.

Details on the analysis, which introduces some new correction for µs alternation, and involves correcting for large afterpulsing in the multispot experiments, are presented in SI-Appendix 12.

## 4 dsDNA smFRET Measurements

To fully characterize the performance of the multispot setup, we performed smFRET measurement on a series of doubly-labeled dsDNA samples using a standard single-spot µs-ALEX setup and our 8-spot setup. All samples possess the same 40 base-pair (bp) sequence used in previous works [19, 30], one strand being labeled with the acceptor dye (ATTO647N) at its 5’-end, while the complementary strand was labeled internally with the donor dye (ATTO550). The donor position depends on the sample, and was chosen so that the D-A separation were 7, 12, 17, 22 and 27 base pairs. The sequence was designed such that the 3 base pairs surrounding each donor labeling sites were identical at each locus (see SI-Appendix 13). We will refer to these samples as the 7d, 12d, 17d, 22d, 27d and DO (D-only) samples.

### 4.1 Single-Spot Results

Single-spot µs-ALEX measurements were performed on a dedicated setup and at a different time than the multispot experiments. Moreover, although the same sample stocks were used, sample preparation was distinct, and therefore the concentration and respective fraction of singly- and doubly-labeled molecules in the two types of experiments were most certainly different.

In addition to sample concentrations, many other acquisition parameters may have differed between the two sets of experiments (e.g. excitation intensity, background levels, correction factors, etc.), all of which could potentially affect the results in both types of measurements. In order to be able to compare data acquisition parameters in single-spot experiments with those of the multispot experiments, we characterized several independent observables: mean count rates and background count rates, peak burst count rates, burst number per unit time and FCS analysis. In the interest of space and in order to focus on the multispot results, characterization of the single-spot setup can be found in SI-Appendix 14.

smFRET analysis of each sample was performed as outlined in Section 3. Donor leakage was extracted from the D-only population proximity ratio (*PR*) peak and the acceptor direct excitation from the A-only population stoichiometry ratio (*SR*) peak, as described in details in SI-Appendix 10. We then applied these two corrections to the PR and SR distribution, to obtain an estimate of the *γ* correction factor [30]. In Fig. 1 we report a comparison of corrected FRET efficiencies (*E*) for each sample, estimated using different methods (see details in SI-Appendix 10 & 11). In all cases, the FRET population was isolated by selecting bursts with *D_ex_* counts ≥ 30 and within an appropriate region of interest in the ALEX histogram.

*PR KDE* are the results of kernel density estimation (KDE) of the PR distributions, and report the position of the KDE maximum, corrected as described in SI-Appendix 9 to obtain *E KDE*. *PR Gauss* are the peak positions of a Gaussian fit to the PR histograms, while *E Gauss* values are obtained by the same correction mentioned for *PR KDE*. Finally, the last two values report results of shot-noise analysis (SNA, discussed in SI-Appendix 11). *SNA mean E* are the values of the mean of the fitted *E* probability distribution and *SNA max E*, the modal values of these distributions.

As evident in Fig. 1, all methods yielded very similar results for each sample.

**Fig. 1.**
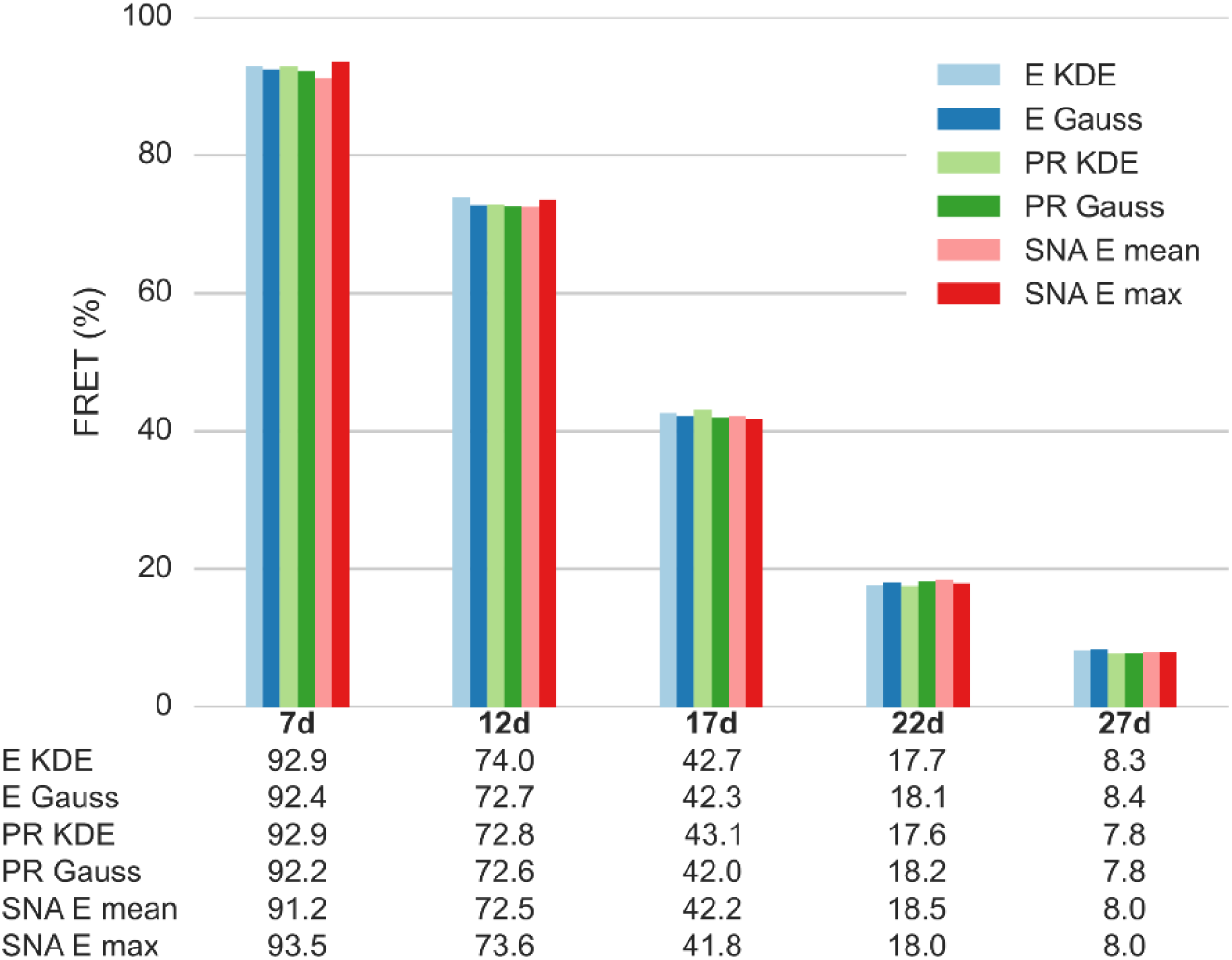
Single-spot µs-ALEX smFRET efficiency for 5 dsDNA samples estimated using different methods. Population-level FRET efficiency estimated for each of the 5 dsDNA samples: 7d, 12d, 17d, 22d and 27 (the number represents the separation in base-pair between D and A dyes). FRET is estimated using the following methods. *E KDE* – KDE maximum from the corrected E distribution. *E Gauss* – Gaussian fit of the corrected E histogram. *PR KDE* – KDE maximum of the *PR* distribution, *PR* value converted to *E* as described in SI-Appendix 9. *PR Gauss* – Gaussian fit of the *PR* histogram, *PR* value converted to *E* as described in SI-Appendix 9. *SNA mean E* – Mean of the FRET distribution returned by SNA analysis. *SNA max E* – Mode of the FRET distribution returned by SNA analysis. For full details of the analysis see section of *μs-ALEX: Corrected E figure* of the accompanying notebook (<u>link</u>). An overview of the computational notebooks can be found in SI-Appendix 2.

### 4.2 Multispot Results

As in the single-spot µs-ALEX experiments (SI-Appendix 14), potential differences in sample concentration, excitation intensity, or other setup characteristics were investigated thoroughly. The multispot geometry introduces additional degrees of variability, due to possible differences between excitation spots geometry and SPAD alignment.

As described in SI-Appendix 3, the excitation spots are generated by direct phase modulation of an expanded Gaussian beam. Because the Gaussian beam is centered in the middle of the linear pattern, lateral spots exhibit a lower intensity than the center ones. In addition, due to the size of the excitation pattern (> 30 µm in the sample plane, see SI-Appendix 3), some geometric aberrations are expected towards the edge of the pattern. While these different imperfections can be characterized by direct imaging [9, 35], this is a time-consuming process, which does not make it practical for routine characterization. Moreover, information obtained by this kind of measurement cannot be used straightforwardly for data analysis corrections. Instead, we examined different measures of the differences between spots extracted from the data itself, using several burst statistics as discussed next (details in SI-Appendix 15).

Finally, smFRET analysis involves correction factors (Section 3.6) which could potentially depend on the spot under consideration. We therefore characterized their dispersion, as described in the following. However, we obtained highly uniform FRET histograms across the 8 spots (with peaks positions varying by at most 2%) without the need to apply channel-dependent corrections for γ-factor, donor-leakage and acceptor direct excitation.

#### 4.2.1 Signal versus spot

The count rate collected from a spot depends on its excitation PSF, the number and type of molecules diffusing through it, as well as the combined detection efficiency PSF of the optics and detector. In freely-diffusing single-molecule experiments, single-molecule bursts are superimposed to a stationary (or slowly varying) background coming from detector dark counts and sample background. In single-spot µs-ALEX measurements, the contribution of SPAD dark count rate (DCR) was negligible (<250 Hz). For the multispot measurements, however, some of the detectors had DCR which dominated other sources of background (see Table 1). Subtracting the DCR contribution to the total count, provides a signal proportional to the product of excitation and detection efficiency which can be used to compare different spots.

Fig. 2 shows the DCR-corrected total signal discussed before as a function of detector for the 6 dsDNA samples. In each measurement, the signal is normalized to 1 for the detector with the highest signal. A clear signal reduction in the lateral spots compared to the central ones is visible, which reflects in part the expanded Gaussian beam profile used to generate the pattern.

**Fig. 2.**
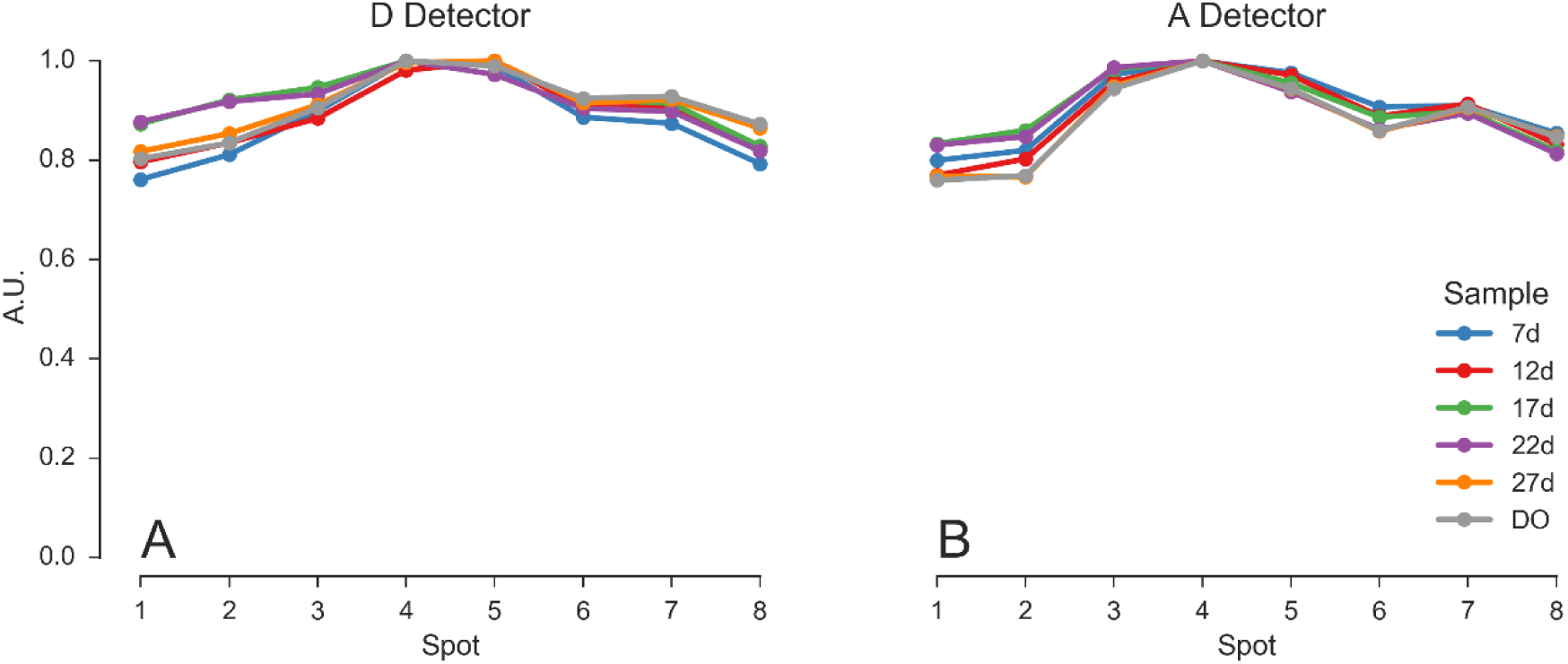
DCR-corrected total count rate as a function of the pixel. Characterization of the excitation × detection profile as a function of spot. Values computed from DCR-corrected count rate in each pixel for different samples. (A) Signal detected by the SPAD array in the D emission channel. (B) Signal detected by the SPAD array in the A emission channel. The samples are the 5 doubly-labeled dsDNA samples (7d - 27d) plus the D-only dsDNA sample (DO). For each sample, the signal is normalized to 1 for the detector with the highest signal. For computational details see SI-Appendix 2 and section *Signal vs spot* (link) of the accompanying Jupyter notebook.

This variation of the signal across spots may be due to variation in the excitation intensity and detection efficiency, indicating that either one or both change as we move from the center of the spot pattern toward its edges. FCS analysis provides complementary information helping to remove the ambiguity, as discussed in the next section.

#### 4.2.2 FCS Analysis

Fluorescence correlation spectroscopy (FCS) analysis of each spot signal can in principle provide information on sample concentration and diffusion constant, if the excitation/emission PSFs are known. In multispot experiments, the diffusing species is identical for each spot, therefore, FCS analysis can be used to detect and quantify differences in excitation/emission PSFs among spots (SI-Appendix 12).

Fig. 3 represents the diffusion times obtained from the fit of the donor (resp. acceptor) autocorrelation functions (ACFs) and of the donor-acceptor cross-correlation functions (CCFs) for the different samples (curves shown in Fig. SI-20), whose average and standard deviations are reported in Table 3. Because of very low to non-existent acceptor signal, no fits of the acceptor ACFs could be performed for samples DO, 27d and 22d.

**Fig. 3.**
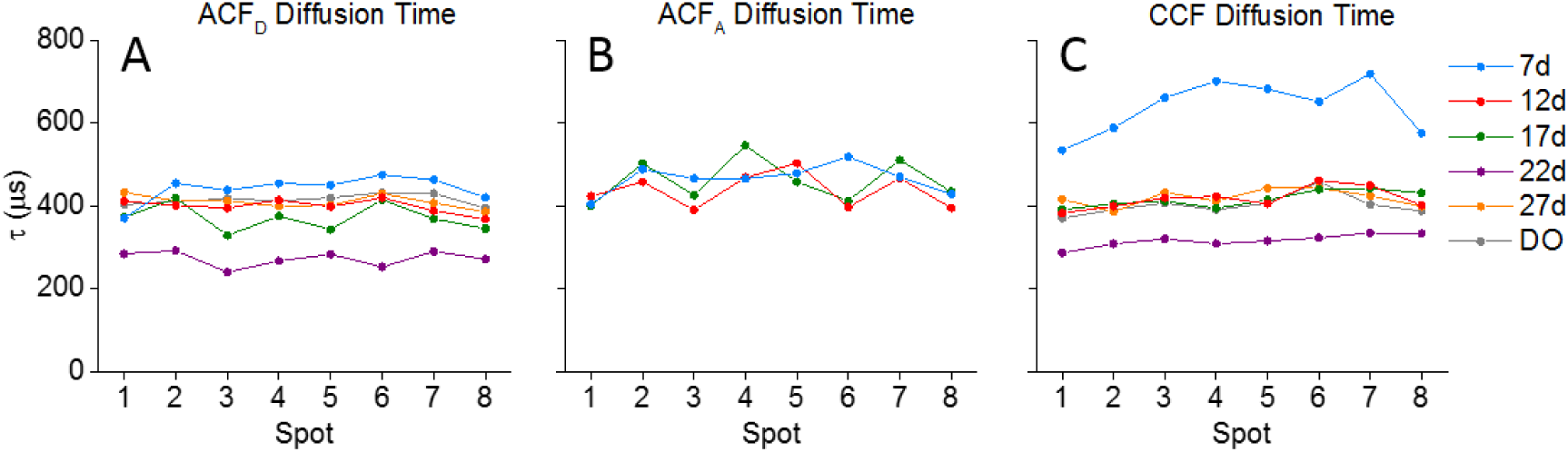
Multispot FCS analysis. For each sample, 8 donor autocorrelation functions (ACFs) and for samples with sufficient acceptor signal (7d, 12d and 17d), 8 acceptor ACFs, were fitted with a 2D diffusion model with multi-exponential afterpulsing components. Additionally, for all samples, the cross-correlation function (CCF) of the 8 donor and acceptor signals were computed. Each curve was fitted with the respective model described in the text. (A) Diffusion times for the donor ACFs. (B) Diffusion times for the acceptor ACFs. (C) Diffusion times for the donor-acceptor CCFs. A significantly shorter diffusion time is observed for the 22d sample in both the ACFs and CCFs. Sample 7d is characterized by a significantly larger diffusion time in the CCF only.

For each sample, apparent diffusion times from ACFs are relatively uniform among spots (~400 µs), suggesting that the observation volumes of the different spots are similar. Moreover, acceptor ACF diffusion times are only marginally larger than their donor counterpart, suggesting a minimal chromatic effect. Of particular note, however, is the systematically reduced diffusion time of ~300 µs observed across all spots for the 22d sample (Fig. 3A, purple). A possible explanation for this observation (detector misalignment) will be discussed after we examine the complementary information provided by CCF analysis.

The CCF fits provide similar diffusion times (*τGR*) for most samples. In particular, a lower diffusion time is computed for sample 22d as well. Since for donor-acceptor CCF, the diffusion time is theoretically equal to the mean of donor and acceptor diffusion times [36]:

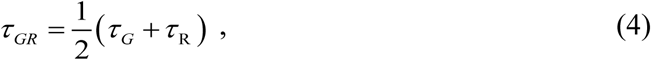

a lower value indicates that the corresponding acceptor channel ACF diffusion time *τR* (which could not be measured due to insufficient acceptor signal in sample 22d) is comparably small. Eq. (4) is indeed verified for sample 12d and 17d, for which both *τG* and *τR* are available (Table 3).

For sample 7d, however, the CCF exhibits a much larger diffusion time and larger variance than their ACF counterparts (~650 µs versus ~450 µs), suggesting that misalignment of one of the two detector arrays is responsible for this discrepancy. Using Eq. (SI.83), with parameter *τGR* fixed for all spots to the value obtained from Eq. (4), a value of the relative shift between donor and acceptor PSFs, *d*/*ωGR* = 0.56 ± 0.06 was obtained, where the uncertainty is equal to the standard deviation over the 8 spots (see SI-Appendix 12). Assuming a typical PSF waist parameter, *ωGR* ~ 100 nm, this corresponds to a *d* ~ 60 nm shift in the sample plane, or equivalently, taking into account the 60× magnification of our setup, a ~ 4 µm shift in the detector plane. This is a rather small shift, which fortunately does not significantly affect the number of photons collected during each burst.

**Table 3.**
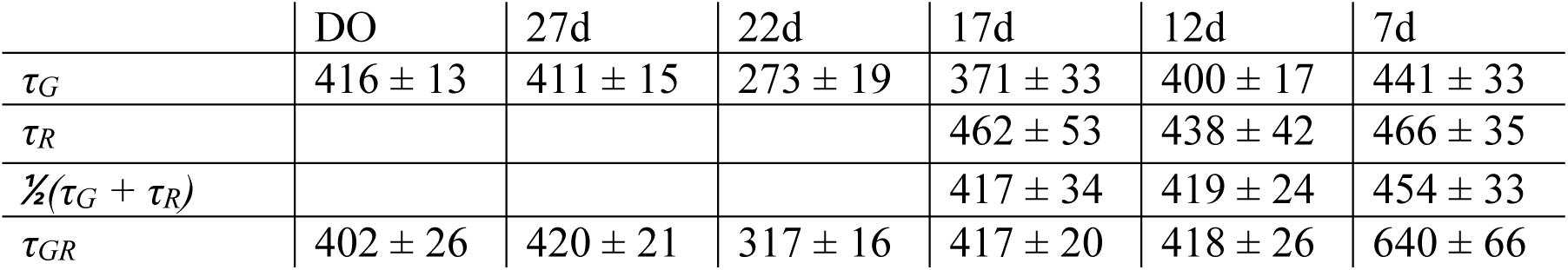
8-spot average of diffusion times (μs) fitted from ACF and 2-color CCF curves. Average diffusion times (*±* 1 standard deviation) obtained from correlation function fits (in µs). *τ_G_*: donor-ACF, *τ_R_*: acceptor ACF, *τ_GR_*: donor-acceptor CCF diffusion time obtained for a 2D diffusion model with no offset. No fit was performed for the acceptor ACF (A-ACF) of sample DO, 27d and 22d due to lack of sufficient acceptor signal.

|  | DO | 27d | 22d | 17d | 12d | 7d |
| --- | --- | --- | --- | --- | --- | --- |
| $\tau_G$ | $416 \pm 13$ | $411 \pm 15$ | $273 \pm 19$ | $371 \pm 33$ | $400 \pm 17$ | $441 \pm 33$ |
| $\tau_R$ | | | | $462 \pm 53$ | $438 \pm 42$ | $466 \pm 35$ |
| $\frac{1}{2}(\tau_G + \tau_R)$ | | | | $417 \pm 34$ | $419 \pm 24$ | $454 \pm 33$ |
| $\tau_{GR}$ | $402 \pm 26$ | $420 \pm 21$ | $317 \pm 16$ | $417 \pm 20$ | $418 \pm 26$ | $640 \pm 66$ |

#### 4.2.3 PSF Characterization by Burst Analysis

When the sample is homogeneous, as is the case across the spots in the multispot system, and burst search criteria are chosen appropriately (as discussed in SI-Appendix 7), burst statistics such as *burst size* and *burst duration* can be used to characterize the product of the excitation-detection PSF. Burst size, for instance, is directly related to the integral of the excitation-detection PSF along the molecule’s trajectory, while burst duration is related to the PSF size. Additionally, the *peak count rate* reached in each burst depends directly on the peak excitation PSF intensity. All these burst statistics are characterized by probability densities with an asymptotic exponential behavior due to diffusion and the profile of the excitation/detection volume [31]. This property allows estimating mean values for each of these burst statistics, as described in SI-Appendix 15.

Using these statistics, one needs to keep in mind that burst search parameters may influence some of them, such as burst size and duration distribution. For instance, the count rate is low at the beginning of a burst, increases and eventually decreases at the end. Therefore, considering a particular burst, increasing the photon rate threshold used for burst search, will reduce its duration and size (the burst will start later and end earlier, everything else being equal).

On the other hand, the peak count rate during a burst is typically attained well within the burst and therefore it is expected to be less, if at all dependent on the count rate threshold used for burst search.

Using a minimum SBR burst search, a larger background results in an increase in the count rate threshold used for burst search (Section 3.3). This causes a reduction in burst duration and size for spots characterized by a larger background rate, other spot characteristics being identical. This situation is relevant here, since a significant fraction of the background is due to detector dark counts, which differ significantly between SPADs within each array (Section 2.2). The peak count rate for identical species, however, should not be affected, provided all other acquisition parameters are identical (e.g. excitation intensity and alignment).

Fig. 4 shows the effect of different burst searches on DO burst statistics for each of the 6 dsDNA samples. Focusing on the DO population eliminates complexities introduced by the different measured brightnesses of the FRET population of each sample (SI-Appendix 15). The top row (Fig. 4A-C) shows results obtained with a minimum SBR criterion burst search, while the second row (Fig. 4D-F) reports results obtained with a fixed threshold burst search.

**Fig. 4.**
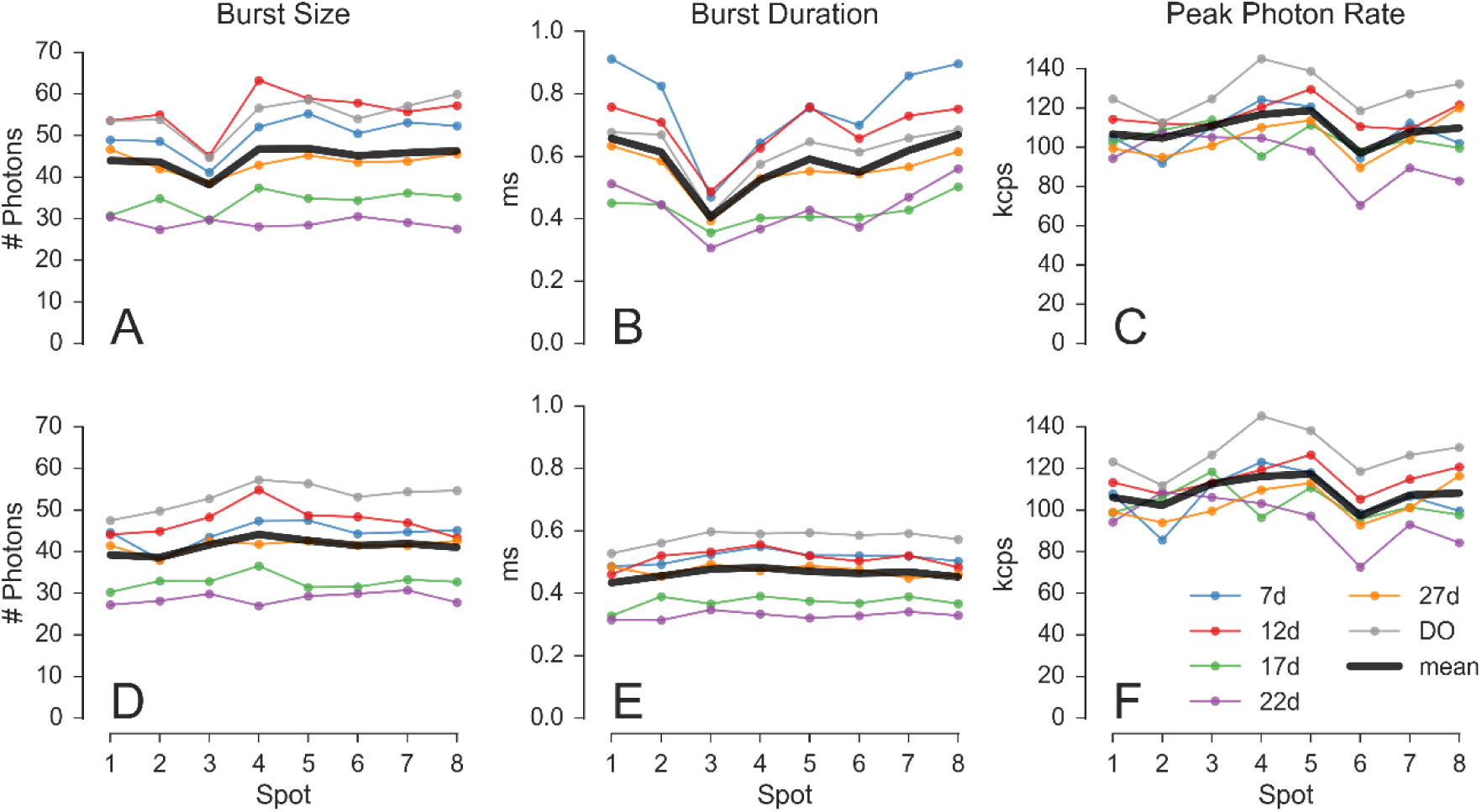
Burst statistics versus spot. Burst statistics extracted for the D-only population in the series of 6 dsDNA samples (7d, 12d, 17d, 22d, 27d and DO) using burst search on D_em_ photons. The first row (A, B, C) shows results obtained with a minimum SBR criterion burst search. The second row (D, E, F) show results obtained with a fixed threshold burst search. The three columns report different burst statistics as a function of the spot number. A, D: Mean burst size. B, E: Mean burst duration. C, F: Mean burst peak count rate. While the peak count rate is largely invariant from the type of burst search (C & F), burst size and duration are affected. In particular, the uneven DCR distribution of the donor SPAD array (maximum DCR in pixel 3), causes a dip in both mean burst size and duration (A & B). The DCR influence is eliminated when using a fixed threshold (D & E). Graphs D - F show the fairly uniform properties of the excitation-detection PSFs across the different spots (see main text). Additional computational details can be found in Section Burst statistics vs spot (link) of the accompanying Jupyter notebook.

Burst size and duration computed in the first case (panel A & B) both show a dip for spot 3, whose SPADs are characterized by the highest DCR (Table 1). By contrast, burst size and duration computed in the second case (panel D & E) show a flatter profile, independent from DCR, suggesting minimal differences between spots. Differences from one experiment to the next can be observed, which can be explained by different excitation intensities.

Fig. 4C & F show the mean peak count rate for both types of burst search. Their similarity demonstrates that, as expected, the peak count rate is largely independent from the burst search parameters (unlike burst size and duration). The mean peak count rate is therefore a reliable reporter for the excitation-emission PSF peak intensity.

Finally, looking at Fig. 4D & E, we observe that, in agreement with the results of FCS analysis, the characteristics of the different spots in a given experiment are fairly uniform. Burst analysis also reproduces the difference between the measured samples obtained from FCS. In particular, the mean burst duration (*i.e.* the PSF width, since all samples are characterized by the same diffusion coefficient) is lower for samples 22d and 17d, suggesting differences in alignment in these measurements.

These results demonstrate that burst analysis following the proper burst search (fixed count rate threshold) can provide information on PSF size and peak intensity consistent with information obtained by FCS analysis. Reciprocally, when comparing a series of samples, bursts analysis can be used to estimate changes in molecular properties such brightness and diffusion coefficient.

#### 4.2.4 Donor Leakage Factor

Because the data collected from each spot involves two separate SPADs, and the SPAD array alignment is a global process, it is possible that acceptor signal is collected with different efficiency in different spots. As a consequence, the donor leakage factor characterizing the amount of donor signal collected in the acceptor channel could vary across spots. Moreover, because occasional realignment was performed in between measurements, it is not even certain that the leakage coefficient was identical for all samples. For these reasons, we computed the donor leakage factor for each spot individually as described in SI-Appendix 9. Since this estimation requires isolating the D-only subpopulation on the basis of the PR histogram, it was only performed with the 7d, 12d, 17d and D-only (DO) samples. It turns out that the leakage coefficient is small (3-4%), therefore applying a spot-specific correction is not critical. This analysis, however, serves as further characterization of spot uniformity.

Fig. 5 compares the individual leakage coefficients for each spot and for each measurement (see also SI-Tables SI-4 to SI-6 and Fig. SI-17). In the left panel (A) the estimated leakage coefficients are grouped by sample while in the right panel (B) are grouped by spot number. The DO sample yields slightly lower values for all spots (Fig. 5A), while spot 1 appears to have a larger leakage factor for all samples (Fig. 5B). However, the absolute variation is of the order of 1% and results in negligible changes in the computed FRET efficiency *E* values. For this reason, we used a fixed value of *l* = 3.3% for all spots and samples in the remainder of the multispot analysis.

**Fig. 5.**
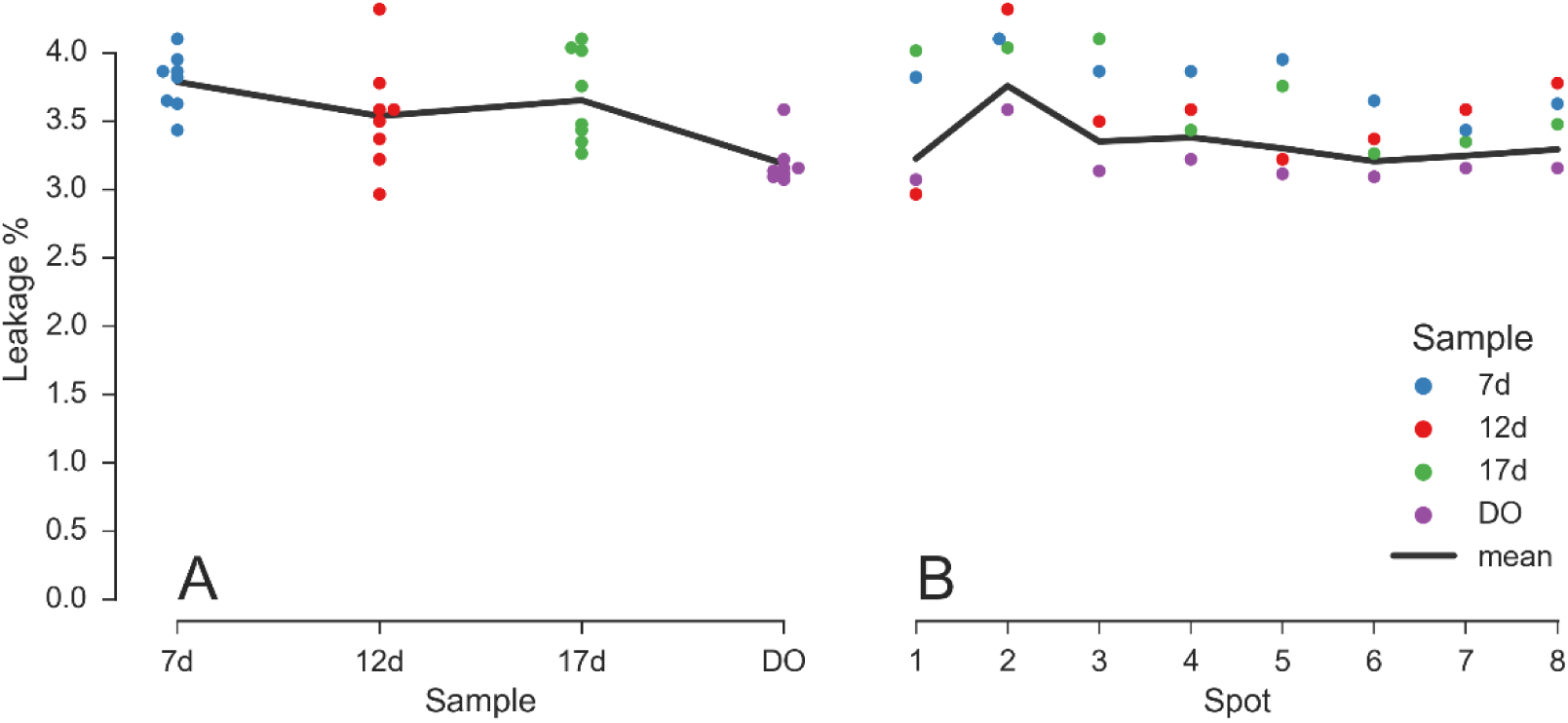
Leakage coefficient for different samples and spots. Leakage coefficient estimated for each spot for dsDNA samples 7d, 12d, 17d and DO. Each dot is the estimated leakage coefficient for a given spot and sample (the color indicates the sample). The left panel (A) shows the dependence of leakage versus sample, while the right panel (B) shows the dependence of leakage versus spot number. Black lines are weighted mean with weights proportional to the number of bursts detected in each sample (A) or spot (B). Both graphs show a good uniformity across samples and spots, with leakage factors in the 3-4% range. For computational details (including the numerical values used in this figure) see section *Leakage coefficient* (link) of the accompanying Jupyter notebook.

#### 4.2.5 Direct Acceptor Excitation Factor

Due to the absence of an acceptor-specific laser excitation signal, the contribution to the acceptor signal due to direct excitation of the acceptor by the donor-excitation laser needs to be expressed as a function of the total (*γ*-corrected) burst size as explained in SI-Appendix 9 [30, 37]. The corresponding factor, *d_T_*, depends only on the dye pair and can be therefore estimated from µs-ALEX measurements (Section 3.6 and SI-Appendix 9).

In this work, we found that the mean value of this factor was *d_T_* = 4.9%. For computational details, see Section Direct excitation: physical parameters (link) of the accompanying Jupyter notebook.

#### 4.2.6 Correction Factor *γ*

As discussed in SI-Appendix 9, the multispot correction factor *γ_m_* is computed from the measured *PR* value of a sample, so that the *γ*-corrected FRET efficiency *E* of that sample matches the corresponding FRET efficiency measured in the single-spot µs-ALEX measurement (Eq. (SI.61)). This can in principle be done individually for each spot, and separately for each sample. Considering the low dispersion of the donor leakage correction factors (Section 4.2.4) and the good uniformity across spots of the FCS analysis results (Section 4.2.2), we only report here the average *γ_m_* factor (and its standard deviation across spots) for each sample. Table 4 reports *γ_m_* estimated using different *PR* analysis methods. The results for samples 22d and 27d are also shown, even though they are strongly affected by the overlap of the FRET and D-only *PR* distribution and are therefore unsuitable for *γ* factor estmation.

**Table 4:** Multispot γ factor. Line 1-3: multispot *γ_m_* values and their obtained from Table 5 and Fig. 1, using Eq. (SI.61) with *PR* estimates obtained by Gaussian fit, KDE or SNA, respectively. Values computed for samples 22d and 27d (indicated in italics) are unreliable due to the overlap between DO and DA *PR* peaks. Last line: factor *K* (Eq. (SI.65)), proportional to the uncertainty on *γm*, computed for the Gaussian fir approach. Values for the other methods are comparable. A common set of parameters *d_s_* = 0.061, *β_s_* = 0.81, *l_m_* = 0.033 was used, average of the values obtained with the various analysis methods.

|  | 7d | 12d | 17d | 22d | 27d |
| --- | --- | --- | --- | --- | --- |
| Gaussian | $0.44 \pm 0.03$ | $0.45 \pm 0.04$ | $0.46 \pm 0.02$ | $0.37 \pm 0.09$ | $0.53 \pm 0.08$ |
| KDE | $0.41 \pm 0.03$ | $0.39 \pm 0.03$ | $0.41 \pm 0.05$ | $0.24 \pm 0.09$ | $0.54 \pm 0.09$ |
| SNA | $0.41 \pm 0.03$ | $0.44 \pm 0.02$ | $0.43 \pm 0.02$ | $0.29 \pm 0.09$ | $0.64 \pm 0.20$ |
| $K$ | 3.5 | 1.9 | 2.4 | 4.6 | 8.5 |

We observe a reasonable uniformity between the different estimates for samples 7d-17d. Error analysis reported in SI-Appendix 9 (Eq. (SI.65)) provides *K*, a scaling factor proportional to the uncertainty on *γ_m_*. The minimum *K* value is obtained for sample 12d, which was therefore used as reference in the remainder of this work (*γ_m_* = 0.45).

#### 4.2.7 FRET Results

The *PR* histograms of most spots and samples yielded two peaks, the lowest one corresponding to the donor-only (DO) population and the largest one to the doubly-labeled (DA) population. In the low FRET cases (sample 22d and 27d), only one peak could be resolved. Its location was used to estimate the *PR* value of the DA population. While this location is probably biased towards small values due to the presence of a residual DO population, the DO fraction observed in the corresponding samples studied on the single-spot µs-ALEX setup was below 10%. Gaussian fitting of the 22d and 27d data assuming an additional DO fraction (kept below 20%) did not significantly affect the peak position estimate (*ΔPR/PR* < 2%).

The results of Gaussian fit, KDE and SNA analysis of the different samples and spots are provided in SI-Table SI-13 to SI-17. Analysis of the pooled data of all spots for each sample yielded results essentially identical to the average of all individual spots. The resulting proximity ratios are reported in Table 5.

**Table 5:** Fitted PR values. Mean and standard deviation of *PR* histogram peak values obtained using Gaussian fit, KDE or SNA analysis (SNA mean value).

|  | 7d | 12d | 17d | 22d | 27d |
| --- | --- | --- | --- | --- | --- |
| Gaussian | $0.85 \pm 0.01$ | $0.57 \pm 0.02$ | $0.29 \pm 0.01$ | $0.12 \pm 0.02$ | $0.10 \pm 0.01$ |
| KDE | $0.85 \pm 0.01$ | $0.55 \pm 0.02$ | $0.27 \pm 0.02$ | $0.09 \pm 0.02$ | $0.08 \pm 0.01$ |
| SNA | $0.82 \pm 0.01$ | $0.55 \pm 0.02$ | $0.28 \pm 0.01$ | $0.10 \pm 0.02$ | $0.08 \pm 0.01$ |

*PR* values from Table 5 were corrected using this value of *γ_m_* and the values of *l* and *d’* obtained earlier, using Eq. (SI.57). They are compared to the previously obtained single-spot µs-ALEX values in the next section

### 4.3 Comparison of Multispot and Single-Spot Results

In this section, we compare the results obtained in the single-spot µs-ALEX measurements and those obtained with the multispot setup, and review the main differences between both measurements, and lessons to be drawn from their comparison.

#### 4.3.1 FRET Efficiencies

Fig. 6 represents the corrected FRET efficiencies obtained in the previous section, together with the results of the single-spot µs-ALEX analysis (Section 4.1). The prediction of a simple DNA model (discussed in SI-Appendix 16) is indicated as a guide for the eye.

**Fig. 6.**
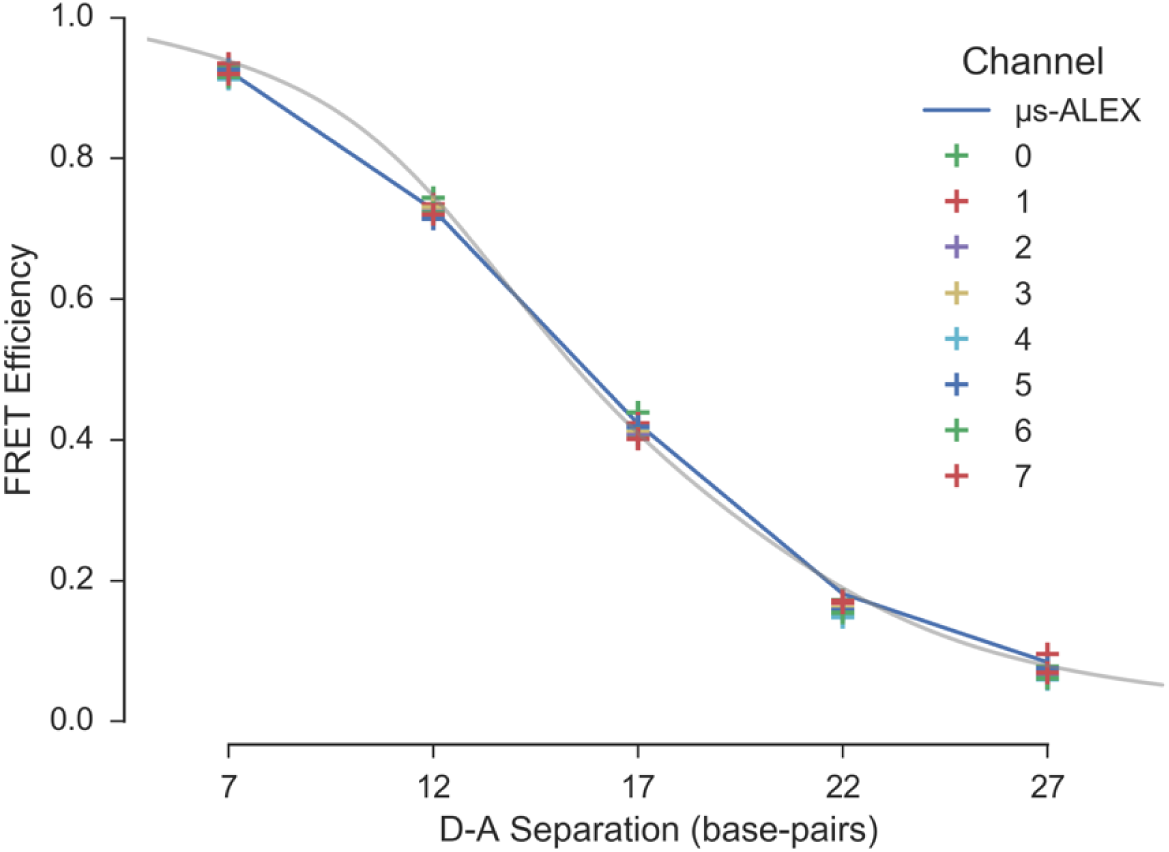
FRET efficiency versus dye separation in the multispot experiments. The *γ*-corrected FRET efficiency values obtained by Gaussian fit are represented as a function of dye separation (in base pair unit). Spots are indicated by different colors. The single-spot µs-ALEX results are represented as a connected line. The distance dependence of E predicted by a simple model of the DNA double helix, to which dyes are attached by a linker at a fixed position, is shown as a guide to the eye. The parameters used in the model are defined in SI-Appendix 16.

The agreement between the two measurements is good, considering the impossibility to separate the contributions of donor-only and low FRET populations for sample 22d and 27d (resulting in an underestimation of the low FRET values in the multispot experiment), and the possible differences between spots in the multispot measurements. It is sufficient for studies concerned with identifying different populations more than extracting precise structural information.

#### 4.3.2 Detection Efficiency Differences

The photon detection efficiency (PDE) of the SPAD arrays in the red region of the spectrum (acceptor emission) is about half lower than that of the SPADs used in the single-spot experiments (Fig. SI-2). While the *γ_m_* factor value of ~0.45 computed for the multispot setup (while *γ* ~ 1 for the single-spot setup) takes care of this difference when computing FRET efficiencies, too small a PDE could result in low detected counts and therefore, low signal-to-noise and signal-to-background ratios.

However, despite this lower PDE, the number of acceptor channel counts per burst (*n_DexAem_*) in the multispot measurements were about 30% larger than in the single-spot µs-ALEX measurements (Fig. 7A-B). A first contributing factor to this surprisingly large acceptor signal is the longer burst duration observed in the multispot measurements (Fig. 7C-D). This increased duration is confirmed by FCS measurements of diffusion times (Table 8 and SI-5), and discussed further in the next section.

Another contributing factor to the measured burst size is the excitation intensity. The peak intensity in a spot can be estimated from the measurement of the peak count rate in that spot. Fig. 7E-F show the distribution of peak acceptor photon count rates during donor excitation (D_ex_A_em_) in both types of measurement. The single-spot results were corrected for the fact that the D_ex_ period represents only about half of the total period, while the multispot results are raw values, uncorrected for the difference in PDE between the two experiments. Because the observed peak count rates are comparable, but the PDE of the detectors used in the multispot experiments is about half that of the detector used in the single-spot experiment, we conclude that the effective acceptor excitation intensity in the multispot measurements was approximately double that of the single-spot measurements.

The fact that an increased excitation intensity was used to obtain similar acceptor burst counts in the multispot measurements has several potential drawbacks. First, due to the cumulated losses in the excitation path, the required output laser power is quite considerable, calling for expensive laser sources. Secondly, since some amount of saturation due to triplet-state blinking was observed in the single-spot measurements with lower excitation intensities (Table SI-5), larger amounts of saturation were likely present in the multispot measurements. Finally, since a pulsed laser was used during multispot measurements, the probability to excite dyes into higher excited states was higher in these experiments, potentially accelerating photobleaching.

While these were not issues in these particular experiments, such a situation is detrimental and will benefit from detectors with better PDE.

**Fig. 7.**
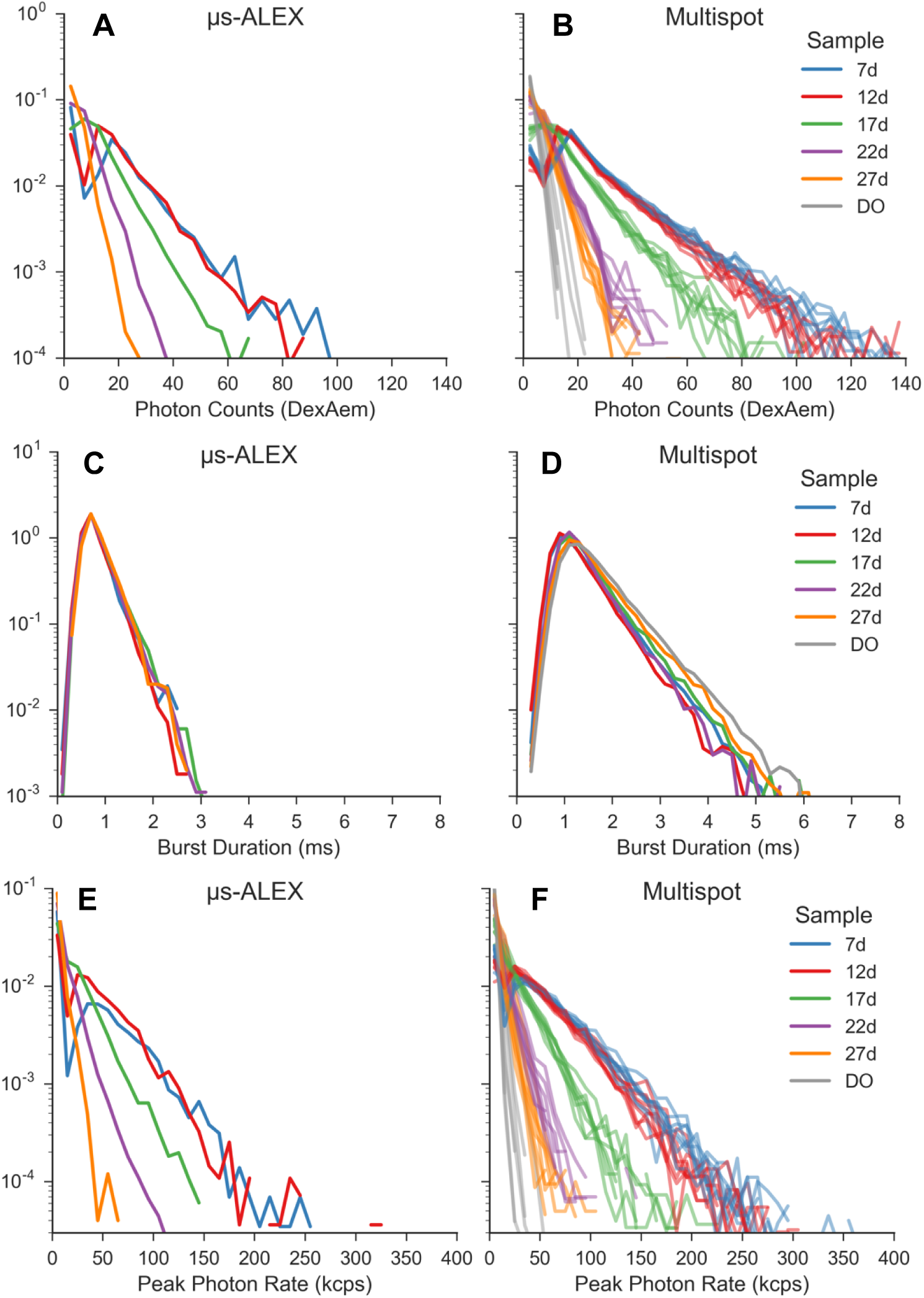
μs-ALEX vs multispot burst data distributions. Distributions of acceptor counts after donor excitation, burst duration and peak photon rate in each burst for both single-spot µs-ALEX (left column) and multispot (right column) experiments. Bursts were searched using the donor-excitation stream and a constant count rate threshold (*r_min_* = 25 kHz for both single-spot and multi-spot measurements), followed by a selection with *γ*-corrected burst size ≥ 15. All the distributions (i.e. histograms) are normalized so that their integral is equal to 1. For the multispot setup, acceptor counts and peak photon rates distributions (first and last row) are reported for each channel separately; for readability, for the multispot burst duration distribution (second row, right), we report the mean across the channels. For more details see section *Burst statistics vs usALEX* accompanying notebook (link).

#### 4.3.3 Setup Characteristics

While the multispot setup is intended to be equivalent to multiple single-spot setups working in parallel, constraints related to the generation of an excitation spot array, and to the alignment of SPAD arrays, create imperfections whose signature can be detected. In particular, as discussed above, both FCS and burst duration analysis indicate significantly longer diffusion times in the multispot than in the single-spot measurements. This increase could be due to two different causes: differences in PSF dimension or saturation effects.

A wider excitation PSF is expected (and has been observed previously [26]) due to the reduced overfilling of the objective lens back aperture by each of the beamlets constituting the excitation pattern source. Moreover, optical aberrations affecting the illumination spots furthest away from the optical axis, could result in deformation of the corresponding excitation PSF [9]. This effect does not appear to be noticeable in the ACF measurements (Fig. 3A & 3B), but could be responsible for some of the differences observed in the CCF apparent diffusion times (Fig. 3C). This was in particular noticeable for sample 7d, which exhibits an increased apparent diffusion time toward the center of the pattern, or sample 22d, which is characterized by a small steady increase from one end of the pattern (spot 1) to the other (spot 8).

Saturation effects could not be directly characterized in the multispot measurements because of complications in the afterpulsing contribution to the ACFs (SI-Appendix 12). However, as mentioned before, some saturation effect was detectable in the single-spot measurements, as demonstrated by the presence of a 10-15% fraction of triplet state blinking (Table SI-5). Since the mean excitation power used in the multispot measurements was about twice larger than in the single-spot measurements, it is plausible that an increased triplet state population was present in the multispot measurements. The use of pulsed excitation in the multispot versus µs alternated CW excitation in the single-spot measurement further complicates the comparison between both types of measurements [38]. Finally, saturation increases the apparent diffusion time obtained from FCS analysis [39], which could further contribute to the longer diffusion times measured in the multispot experiments.

In summary, while they did not affect the results after proper analysis, some of the observed differences between the single and multispot setups could in principle be reduced by using a CW laser source (to reduce saturation).

#### 4.3.4 Limitation of non-ALEX Measurements

The absence of an acceptor excitation laser in the multispot experiment limited the amount of information which could be extracted from the measurements. In particular, correction factors had to be estimated in non-ideal ways:

- The *γ* correction factor had to be estimated from comparison of the proximity ratio obtained in one measurement (12d sample) with the accurate FRET efficiency value computed from the corresponding single-spot µs-ALEX measurement.
- The direct acceptor excitation correction had to be estimated from µs-ALEX measurements and expressed as a function of the corrected burst size (Eq. (SI.53) or (SI.55)).
- For low FRET sample measurements (22d, 27d), no independent estimation of the donor-leakage coefficient *l* could be performed, since the DO population could not be distinguished from the doubly-labeled one (and was probably imperfectly corrected for direct acceptor excitation).

In addition to these issues, proximity ratio estimations were affected by the difficulty (or impossibility for some samples) to isolate singly-labeled from doubly-labeled molecular bursts:

- For all samples, bursts corresponding to molecular transit during which the acceptor blinked or bleached (resulting in a proximity ratio between the DO value and the actual FRET population value) created a “bridge” between the donor-only PR peak and the FRET PR peak, affecting the accuracy with which the peak position could be determined (in particular using a Gaussian fit method, or the shot noise analysis approach).
- For low FRET samples, the overlap between the donor-only and doubly-labeled PR peaks, prevented determining the DO fraction and thus, correcting for its influence on the computed FRET efficiency of the doubly-labeled population.

While these limitations are important, we have shown that a quantitative analysis covering the whole range of FRET efficiencies can be performed with a careful analysis. In cases where analysis is limited to samples with medium to high FRET efficiency, and when no accurate measurement of the FRET efficiencies is needed, the current multispot setup can provide high throughput without overly complex analysis, as demonstrated in the next section.

## 5 RNAP Promoter Escape Kinetics

### 5.1 Introduction

To illustrate the high-throughput capabilities of the multispot system, we studied the kinetics of DNA transcription by the RNA polymerase (RNAP) of *Escherichia coli* (*E. coli*) in a reconstituted *in vitro* assay. RNAP carries the task of DNA transcription, which can be divided into three major steps:

1. Initiation, during which RNAP identifies and binds the promoter sequence of a gene and prepares RNAP for RNA polymerization activity;
2. Elongation, during which RNAP escapes from the promoter sequence and rapidly processes downstream DNA and polymerizes RNA;
3. Termination, during which RNAP identifies a stop signal and terminates the RNA polymerization process.

The strong interaction between the promoter sequence and the *σ* promoter-specificity factor of the RNAP complex delays the transcription process and makes transcription initiation the rate limiting step of DNA transcription in many genes.

In *in vitro* transcription assays, the system starts in an initial state comprised of a stable RNAP-promoter open complex, in which a ~13 bases transcription bubble is formed by melting the promoter sequence between promoter sequence positions -11 and +2 (where +1 is the transcription start-site). In the *in vitro* assay, the open bubble is further stabilized by a dinucleotide to form an initially transcribed complex with a 2 bases nascent RNA (RP_ITC=2_, Fig. 8A). The final state (elongation and run-off, Fig. 8B) is reached after RNAP escapes from the promoter sequence and starts transcribing the sequence downstream from the promoter.

The onset of the transcription reaction is triggered by the addition of all four nucleoside triphosphates (NTPs: ATP, TTP, GTP, and CTP). Due to the timescale of the whole process (< 3 min), direct measurement of the kinetics at the single-molecule level on freely-diffusing smFRET setups is difficult due to the small number of bursts available for population analysis over this short time scale. Here, we report the first real-time measurement of such kinetics at the single-molecule level. Exploiting the higher throughput of the 8-spot smFRET system, we recorded the kinetics of promoter escape from initiation into elongation in the lacCONS promoter [40-42].

### 5.2 Principle of the Experiment

The change in DNA conformation during the reaction is monitored by labeling the two complementary DNA strands with fluorescent probes (donor: ATTO550 and acceptor: ATTO647N). The dyes are positioned within the transcriptional bubble opened in the RP_ITC=2_ state, as shown in Fig. 8. During the transition into elongation, the RNAP moves downstream with the transcription bubble, causing re-annealing of the upstream DNA strand and a corresponding melting of downstream template DNA. Re-annealing of the template DNA causes reduction in the donor-acceptor distance and increase in FRET efficiency. With this labeling configuration, we obtain a medium-FRET population (*PR* = 0.62) for RP_ITC=2_, and a high-FRET population (*PR* = 0.952) for the final state. The medium-FRET population is well separated from the donor-only peak, mitigating the limitation of the single laser excitation configuration of our 8-spot setup.

**Fig. 8.**
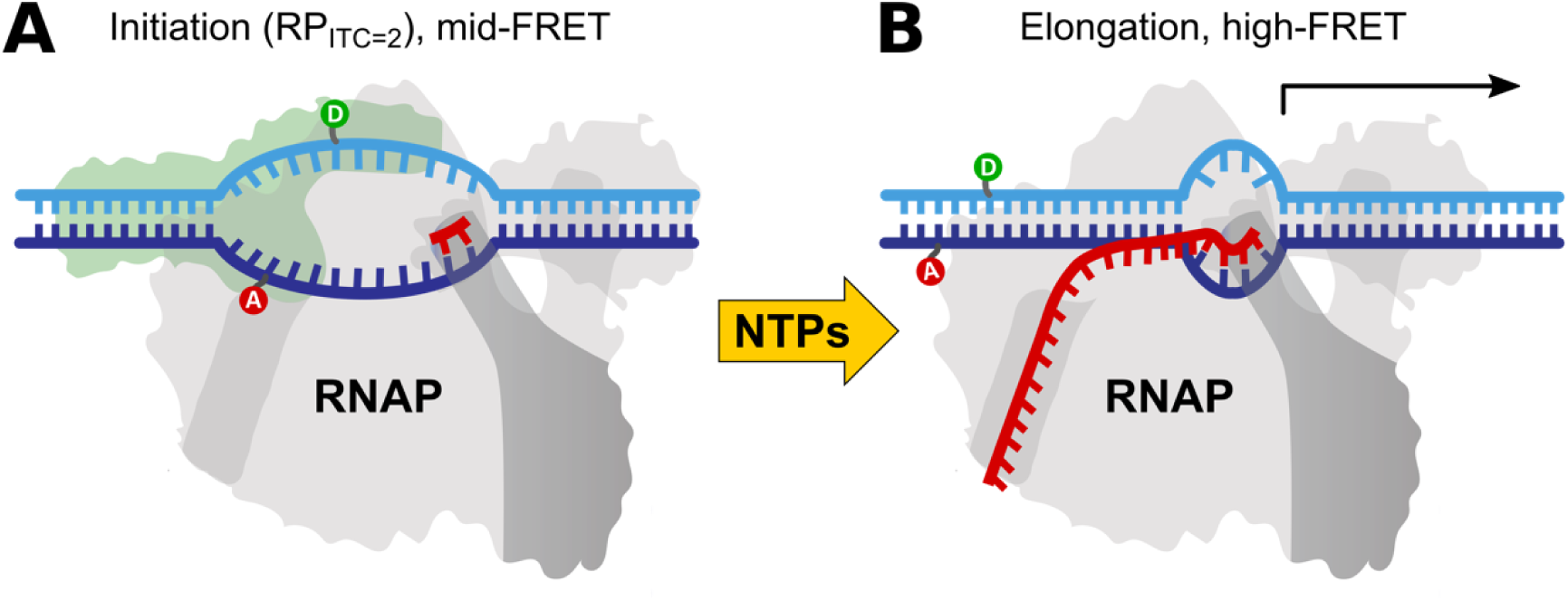
Schematic of the reaction observed in real time with the 8-spot smFRET setup. **(A)** The RNAP-promoter initially transcribed complex (RPITC) is prepared with an initiating dinucleotide (red “π” symbol) as the nascent RNA chain. Complementary DNA strands are labeled at DNA promoter bases with donor (D, green, position -5) and acceptor (A, red, position -8) dyes. After formation of a transcription initiation bubble, the dyes are separated, resulting in medium FRET. The initial state remains in stationary conditions until the addition of the four missing nucleotides (NTPs, yellow arrow), which triggers transcription initiation and elongation. (**B**) During elongation, the transcriptional bubble moves downstream (to the right), causing hybridization of the sequence of the initial transcriptional bubble and a corresponding decrease of the D-A distance (FRET increase).

The measurement is performed during a continuous acquisition comprised of 3 phases:

- Phase 1 (10-15 min): 100 μl of the sample in the RP_ITC=2_ state is placed on the setup and measured in order to acquire an accurate representation of the initial FRET efficiency histogram.
- Phase 2 (<20 s): without stopping the acquisition, 2 μl of NTPs are manually added to the sample in order to reach a final concentration of 100 μM.
- Phase 3 (30-45 min): the acquisition continues until an asymptotic steady state is reached.

### 5.3 Data Analysis

Burst search was performed on the full data set, in windows of constant durations, selecting bursts containing more than 30 counts after background correction (*n_Dex_* ≥ 30). Since the experiment’s purpose was to monitor the transition from one state (medium FRET) to a very distinct one (high FRET), there was no need for *γ-* or other corrections. *PR* distributions exhibited three distinct peaks: DO, medium-FRET (open complex in RP_ITC=2_ state) and high-FRET (hybridized DNA), which were fitted with 3 Gaussians having constant peak positions and widths, while amplitudes were allowed to vary as a function of time.

Two pre- and post-kinetics steady-state regimes (10-15 min long) in the measurement were identified and used to fit the initial and final *PR* histograms. These initial and final *PR* peak positions and widths, were used throughout the whole measurement, the only adjustable parameters being the respective fraction of each peak.

To compute a kinetic curve, we selected bursts within sliding integration windows (duration: 5 or 30 s, step: 1 s). For each window (*i.e.* time point in the kinetics), the amplitudes of the 3-Gaussian model were fitted (Fig. 9A). The ratio of fitted component fractions (high-FRET / (mid-FRET + high-FRET)) was then represented as a function of time (Fig. 9B). A first order kinetic time constant *τ* was fitted using an exponential model incorporating the effect of the integration window. Monte Carlo simulation of simulated kinetic curves with additive Gaussian noise (with variance equal to that observed in the experimental trajectories) were used to estimate the uncertainty on the kinetic time constant for different values of *τ*. Simulations show that time constants as small as 10 s can be reliably measured using this approach (*σ_τ_* = 2.3 s).

**Fig. 9.**
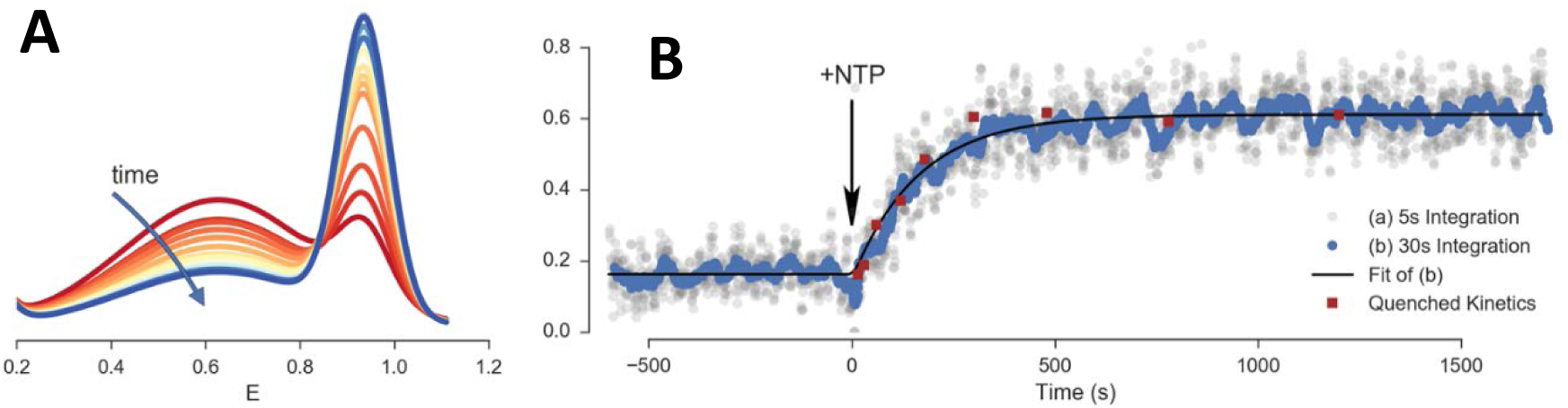
Real time transcriptional bubble closure kinetics results. (A) Evolution of FRET efficiency distribution as function of time (one curve per 30 s). The curves represent Gaussian fit of the FRET histograms. (B) Fraction of high FRET population obtained in the real time kinetics measurement (grey and blue dots) and fraction of probe hybridization to a run-off transcript from quenched kinetics assays (red squares). Dots are computed as a function of time using either a 5 s (grey) or 30 s (blue) moving integration window. The solid black curve is a single-exponential model fitted to the 30 s moving integration window. Quenched kinetics data (from ref. [43]) are normalized to fit initial and final values of real time kinetics trajectory. For more details on the analysis see accompanying *Realtime Kinetics Analysis* notebook (link).

### 5.4 Results

Fig. 9 shows the measured evolution of the high FRET fraction in a single continuous acquisition. Time zero represents the time of NTP injection (Section 5.2) and marks the start of the reaction (RP_ITC=2_ to elongation, Fig. 8). After injection and mixing, the reaction starts with a short delay (< 20 s), due to the time needed for NTPs to reach the excitation volume by diffusion and convection. Data processed using sliding windows of 5 or 30 s illustrate the necessary trade-off between accuracy and time resolution. In this particular case, fit of the 30 s resolution curve was sufficient and led to first order kinetic time constant *τ* = 172 ± 17 s (mean and standard deviation of 3 measurements).

Fig. 9B also shows data points from a series of quenched kinetics experiments red square), obtained with the same system but a completely different experimental strategy (see ref. [43] for details). In this approach, a doubly-labeled complementary ssDNA probe hybridizes with the RNA transcript once the transcription reaction is stopped after a specified reaction time *t*. Hybridization of the probe results in a shift from high to low FRET. As in the experiment reported here, the reaction is started from a RP_ITC=2_ state by addition of NTPs, but it is then stopped (quenched) after a fixed time *t*, by addition of 0.5 M Guanidine Hydrochloride (GndHCl). The sample is then incubated for 30 min for hybridization with the ssDNA probe to occur, and finally measured on a single-spot μs ALEX setup for 10-20 min. The procedure is repeated for different time points (reaction times), yielding the data points (red squares) reported in Fig. 9B.

The remarkable agreement between the two experimental approaches used to probe transcription: (i) quenched kinetics experiments probing the transcript production and (ii) real-time kinetics experiments, probing the change in conformation upon promoter escape, demonstrates the advantages of a multispot approach for tracking kinetics in real-time. From a biochemical point of view, the agreement between the two results confirms that elongation, the step required for producing an RNA transcript after the initiation bubble has moved downstream from the promoter sequence, is faster than our time resolution [3, 44-46]. This means that the run-off transcript production kinetics experiment discussed here mostly probes the initiation part of the reaction, confirming it as the rate limiting step in DNA transcription [47].

## 6 Conclusion and Perspectives

We have presented a detailed description of an 8-spot confocal setup and illustrated its use for smFRET studies with the following examples. First, measurements of a series of freely diffusing doubly-labeled dsDNA samples demonstrated that data simultaneously acquired in different spots could be properly corrected and analyzed in parallel, resulting in measured sample characteristic identical to those obtained with a standard single-spot µs-ALEX setup. We discussed the advantages of a multispot setup, while pointing potential limitations of the current single laser excitation design, as well as analysis challenges and their solutions. Second, we leveraged the increased throughput provided by parallel acquisition to address an outstanding question in the field of bacterial RNA transcription. We showed that real-time kinetic analysis of promoter escape by bacterial RNA polymerase confirmed results obtained by a more indirect route, shedding additional light on the initial steps of transcription.

The concepts discussed here are relevant to future generations of multispot setups, including those using multicolor µs-ALEX or ns-ALEX/PIE excitation schemes [48, 49]. Freely diffusing smFRET and other single-molecule fluorescence applications will directly benefit from one to two orders of magnitude throughput improvements afforded by the development of custom-technology SPAD arrays with enhanced sensitivity in the red region of the spectrum [50, 51], or larger number of detectors and different geometries [16].

Progresses making this technology more accessible can be anticipated in different areas. Improvements to the LCOS-SLM-based excitation scheme presented here can lead to a much better utilization of the laser output power. While this approach provides flexibility in designing the illumination pattern geometry and helps with alignment, simpler and cheaper alternatives can be envisioned.

Efficient, parallel and user-friendly analysis tools will be needed to handle the large amount of data generated by these types of measurements. An open source and rigorous approach to algorithm development will be indispensable to allow cross-validation of massive data sets acquired on different setups across research groups [18].

Finally, coupling this type of setup with other technologies such as microfluidics [52, 53] is poised to transform single-molecule analysis from a niche technology limited to research laboratories to a mainstream and powerful tool with applications in diagnostics and screening [40].

## Acknowledgements

The authors thank Maya Lerner for preparation of RNAP and transcription illustrations. This work was supported by NIH grants R01 GM095904 & R01 GM069709 and NSF grant MCB 1244175.

Conflict of interest statements: S. Weiss discloses equity in Nesher Technologies and intellectual property used in the research reported here. The work at UCLA was conducted in Dr. Weiss's Laboratory. M. Ghioni discloses equity in Micro Photon Devices S.r.l. (MPD). No resources or personnel from MPD were involved in this work.

**Convention used in this document**: underlined text is associated with a clickable web link. To access the linked document, follow the instructions for the PDF reader you are using (generally, Ctrl+click on the link will open the default browser at the specified address). Contact the authors if you encounter any problems with the links or documents.

## Appendix 1 Data Files used in this work

Original data files as well as accessory files useful during data analysis have been deposited on Figshare and given a digital object identified (DOI), which can therefore be used for retrieval and citation. Direct links are provided below.

### 1.1 Photon-HDF5 Files

Photon data files, saved in the Photon-HDF5 format (.hdf5 extension) [1], can be downloaded using the following DOI/links:

- Single-spot µs-ALEX Data Files [2]: https://doi.org/10.6084/m9.figshare.1098961
- Multispot Data Files [3]: https://doi.org/10.6084/m9.figshare.1098962
- RNAP Promoter Escape Kinetics Data Files [4]: https://dx.doi.org/10.6084/m9.figshare.3810930

These files can be analysed using FRETBursts, ALiX or any other single-molecule data analysis software supporting the Photon-HDF5 format described in ref. [1].

### 1.2 Other Files

Files used to correct the single-spot µs-ALEX autocorrelation function for afterpulsing [5] can be downloaded using the following DOI/link: https://doi.org/10.6084/m9.figshare.3817062

## Appendix 2 Software

### 2.1 Jupyter notebooks

**Installing FRETBursts.** In order to execute the notebooks, you need to install FRETBursts first. If you have already installed python through conda just type: conda install fretbursts-c conda-forge Otherwise, see the instruction on the FRETBursts manual.

**Downloading the notebooks.** The <u>multispot_paper github repository</u> [6] contains all the notebooks and results produced for this paper. We suggest to download the ZIP archive from this link. The archive is ~110 MB and contains all the output of the processing (figures and numeric results such as bursts data, fitted parameters, etc.).

**Downloading the data.** Download the dsDNA datasets for single-spot µs-ALEX (link) and for multispot smFRET (link) and put it in the folder *data/singlespot* and *data/multispot* respectively. For the realtime kinetics experiments, download the data files (link) and place them in the folder *realtime kinetics/data/*.

**Reproducibility.** For expert python users, the notebook archive contains a conda environment file that can be used to recreate the exact environment (i.e. the exact version of each library) used during the preparation of this paper.

**Using the notebooks.** The main notebook, index.ipynb (in the root folder of the archive), contains links to all the other notebooks used for the analysis (with a brief explanation of what each notebook does). It also contains links to output data files in CSV format (stored in the results subfolder). The index notebook can also be used to re-execute all the notebooks in a single step and recompute all the paper’s results and figures from scratch. The notebooks for realtime kinetics analysis can be found in the realtime kinetics subfolder of the archive. Therein, the notebook index_realtime_kinetics.ipynb links to all the notebooks of the realtime kinetics analysis.

### 2.2 ALiX Scripts

ALiX is a standalone Microsoft Windows 64 bit standalone application which can be downloaded from its public website (https://sites.google.com/a/g.ucla.edu/alix/), by visiting the Installation webpage for detailed instructions. The website contains an extensive online manual including tutorials. Among the many features of the software is the ability to save settings as well as a list of analysis steps into a “script” text file, which can be reloaded later on to replicate the analysis (or follow the same analysis steps with a different data file). Instruction on how to load and execute a script can be found in the corresponding page of the online manual (General Multispot Analysis).

These scripts have been deposited on Figshare [7] and can be downloaded using the following DOI/link: https://doi.org/10.6084/m9.figshare.3839427. The repository contains a readme.txt file which describes all scripts and the associated data files.

## Appendix 3 Setups and Acquisition Hardware

### 3.1 Single-Spot μs-ALEX Setup

#### 3.1.1 Setup description

The setup used in this work was previously described in ref. [8]. The setup schematic is reproduced in Fig. SI-1.

**Fig. SI-1:**
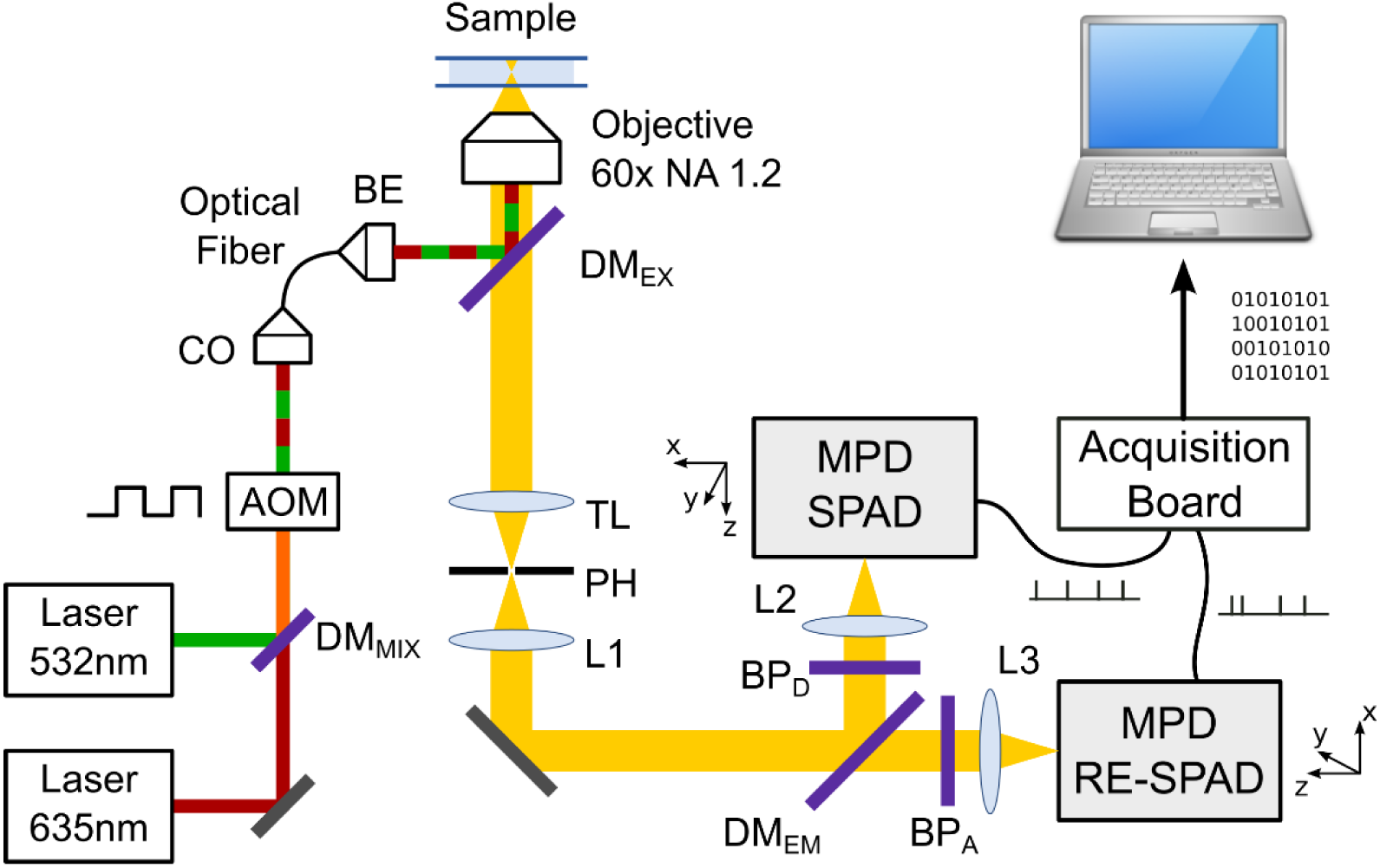
Single-spot µs-ALEX setup. Two CW lasers are alternated using a computer controlled acousto-optic modulator (AOM). The beams are then coupled to a single-mode fiber whose output is expanded using a beam expander (BE) before coupling to the objective lens via a dichroic mirror (DMex). The emitted fluorescence is collected through the objective lens and dichroic mirror and refocused onto a 100 µm pinhole (PH) between the microscope tube lens (TL) and a recollimating lens L1. The two channels (green: D, red: A) are separated using a dichroic mirror (DMem) and focused onto the RE-SPAD using a 75 mm focal length lens (L2 or L3). The TTL output from each SPAD are sent to a counting board (NI-6602) installed in a PC running the LabVIEW acquisition software.

The PDEs of the two types of detectors used in this study are represented in Fig. SI-2.

**Fig. SI-2:**
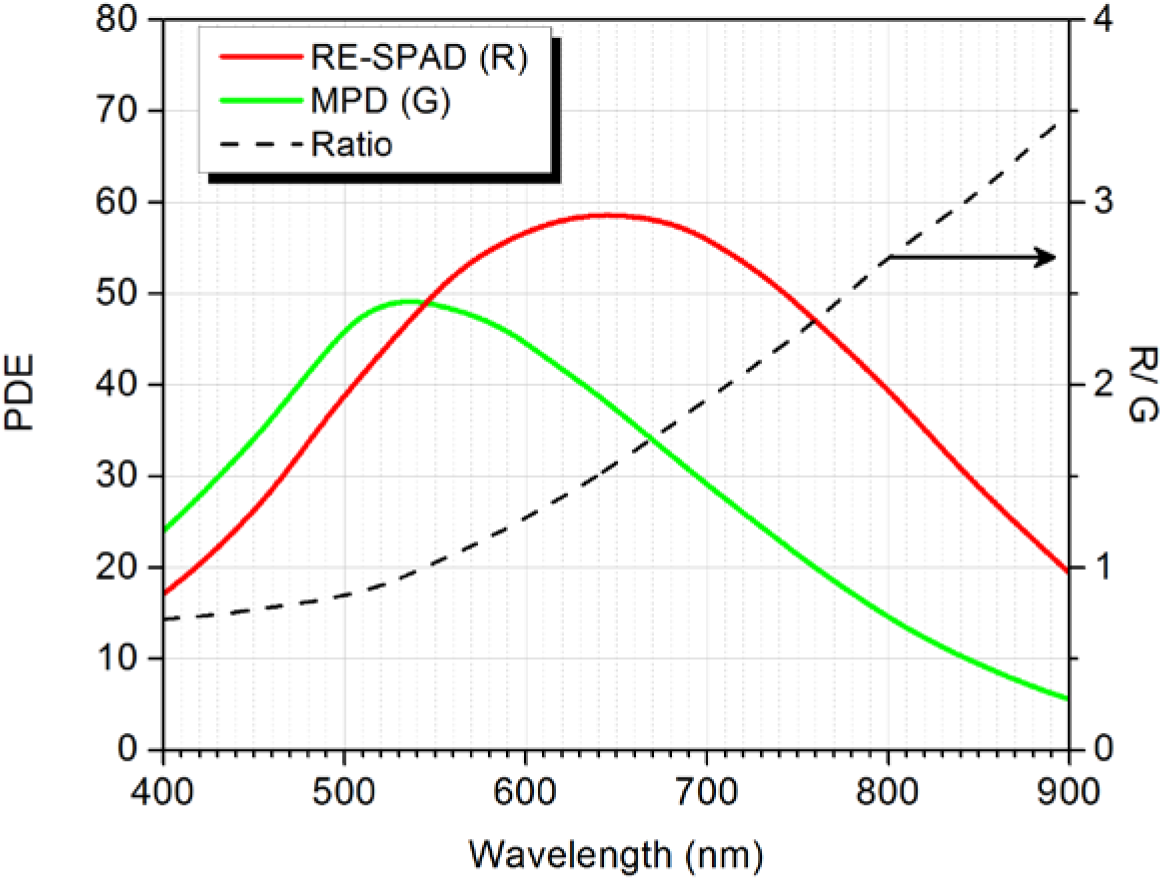
Photon detection efficiency (PDE) of the two types of detectors used in this study. In the single-spot µs-ALEX measurements, a standard technology SPAD from MPD (green) was used for the donor channel, while a red-enhanced SPAD detector from the Polimi group (red) was used for the acceptor channel. The multispot experiments used SPAD arrays manufactured with the same technology as the MPD SPAD and are therefore characterized by the same PDE curve. The ratio of the two PDEs is represented in black (left axis, R/G). The spectral range of the emission bandpass filters (donor: green, acceptor: red) use in the single-spot µs-ALEX measurements are indicated as rectangles on the graph.

#### 3.1.2 Data acquisition

A single digital input/output board (NI PCI-6602, National Instruments, Austin, TX) was programmed using LabVIEW 7.1 (National Instruments) to:

i. send TTL signals (+2.4 V) to each line of the AOM controller, the “on” (resp. “off”) period of each laser alternation corresponding to the +2.4 V (resp. 0 V) state of the corresponding line,
ii. detect and time stamp TTL pulses emitted by each SPAD (one time counter per channel).

TTL inputs and outputs were recorded and generated using a single on-board 80 MHz clock, guaranteeing synchronization of the 12.5 ns resolution time stamps with laser alternation. Each time stamp was recorded with its detector number, and the stream of time stamps recorded in a proprietary binary format (.sm files). Files were converted into the open source HDF5 photon data format (.hdf5 files) [1] and are available as online at URLs indicated in the references [2].

Single spot μs-ALEX measurements for each of the five samples were performed sequentially on a distinct setup from that used for multiple spot measurements. Performing measurements sequentially minimized the possibility that the setup characteristics change from one measurement to another. However, since the measurements were performed on a different day than the multispot measurements, concentration of the samples could be different in the μs-ALEX and multispot measurements.

### 3.2 Multispot setup

A general schematic overview of the setup can be found in Fig. SI-3.

#### 3.2.1 Optics

The 8-spot excitation pattern was generated using a 1 W, 532 nm pulsed laser (IC-532-1000 ps, High Q Laser Production GmbH, Hoheneims, Austria). After laser beam expansion and polarization adjustment, the beam reflects off an LCOS spatial light modulator (LCOS-SLM, model X10468-01, Hamamatsu Corp., Bridgewater, NJ, USA), which generates 8 separate spots on a focal plane a few centimeters from the LCOS surface. The LCOS is a programmable phase modulator, which can impose an arbitrary phase pattern to the incident linearly polarized plane wave (see details below, Section 3.2.5). In the 8-spot setup, the phase pattern creates the same phase modulation as would an array of small lenses (or lenslet array). Whereas a lens imposes a phase difference by putting different amount of material with specific refraction index in the light path (*i.e.* more material in the center of the lens, less at the edges), the LCOS SLM achieves the same result by interposing a thin layer of liquid crystals in the light path, whose light retardation can be controlled by an electric field (electro-optics effect) [9]. By designing a pattern of phase delays similar to that created by a Fresnel lens, the net result is to focus the light incident on that patch of LCOS SLM at a distance (focal length) specified by the pattern (32 mm in our case). Once formed, this pattern of focused beamlets can be manipulated by far-field optics just as any other light pattern.

A recollimating lens L3 (*f* = 200 mm, AC508-200-A, Thorlabs Inc., Newton, NJ, USA) collimates the 8 LCOS-focused spots, directing them toward the objective lens through a dichroic mirror. The unmodulated light, (mostly light incident on the LCOS outside the multispot pattern) is focused at the focal point of L3, where it is stopped by a background-suppressing spatial filter. A 60X water-immersion objective lens (numeric aperture, NA = 1.2, Olympus, Piscataway, NJ) focuses the spots into the sample through a microscope coverslip. The (*x, y*) position, orientation and pitch of the spots can be adjusted by changing the modulation pattern on the LCOS device. A linear arrangement of 8 excitation spots was generated in order to match the linear 8-pixel geometry of the detector.

**Fig. SI-3.**
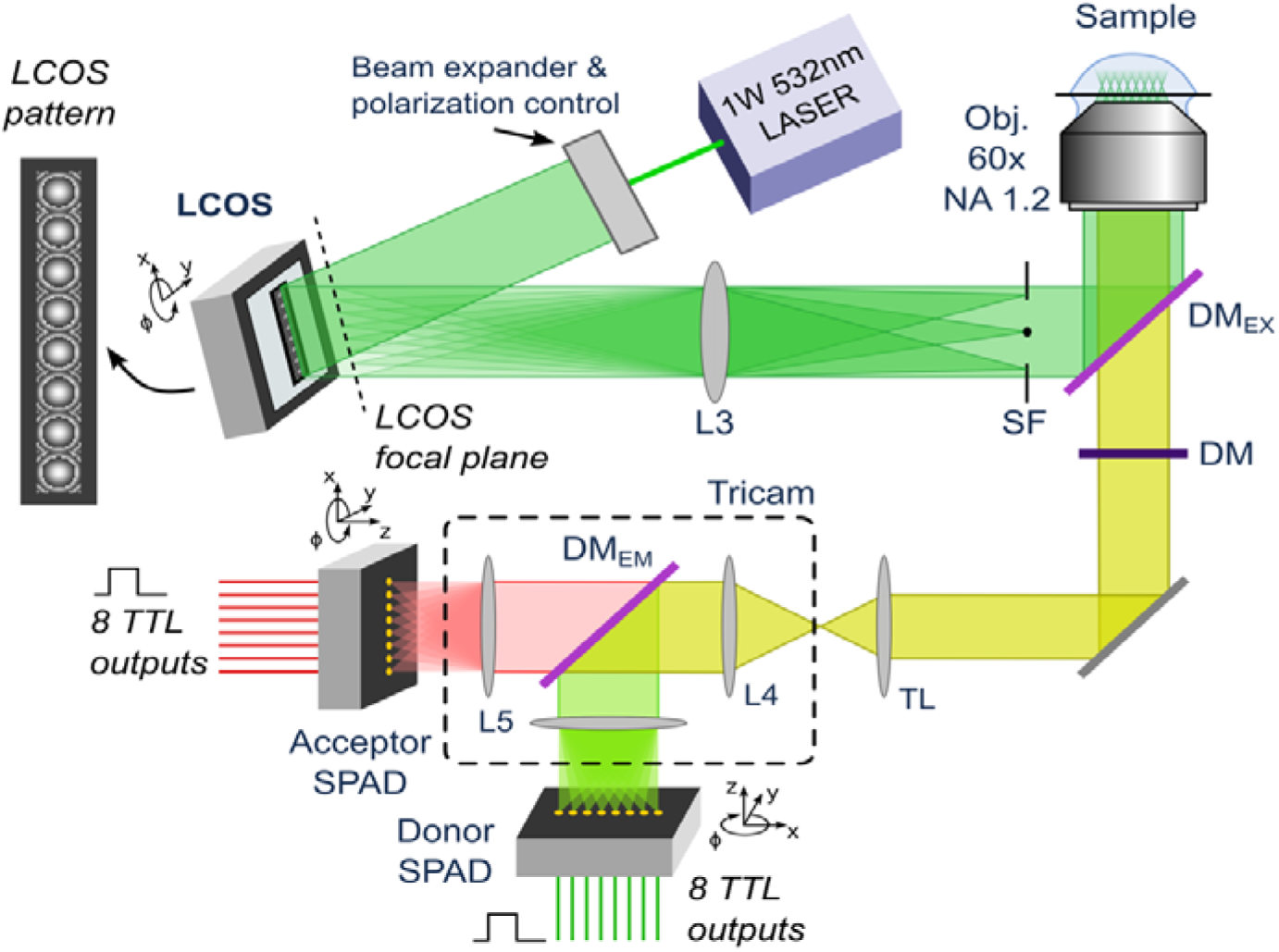
Schematic of the 8-spot single-molecule FRET setup. A freely-diffusing sample is probed via 8 independent excitation spots generated by a 532nm CW laser and an LCOS spatial-light modulator. The fluorescent signal from each excitation spot is optically conjugated to a pair of pixels in the two detectors. The donor and acceptor emission is separated in two distinct spectral bands and detected by two separated SPAD arrays.

#### 3.2.2 Background suppression

The background-suppressing spatial filter used here is an improved version of the pin-dot pattern reported previously [10, 11]. In the original design any light outside the multispot pattern, as well as the small fraction (<10 %) of unmodulated light incident on the multispot pattern, is simply reflected off the LCOS surface. If not blocked, this light generates unacceptable background levels (unfocused sample excitation). The spatial filter is placed at the focal plane of the L3 lens to block this unmodulated light. The pin-dot is a critical component which needs to block a continuous flow of high-power laser and can be easily damaged. Moreover, minimal alignment errors both in the X, Y and Z direction can cause significant increase in background and deterioration of the multispot pattern. To improve this pin-dot design, we use a linear Bragg grating pattern on all the LCOS area outside the multispot pattern.

The Bragg pattern creates a series of diffraction spots on the L3 focal plane. The 0-th order diffraction lays on the optical axis while the higher orders are offset in a direction orthogonal to the Bragg pattern. Using the highest spatial frequency achievable with the LCOS (one line with phase 0 and one with phase **π**), more than 90% of the background light is steered away from the optical axis, into the first and higher diffraction orders. The higher orders are blocked in the L3 focal plane by two narrows stripes of absorbing material. This design achieves better background suppression than the pin-dot-only version and improve the long-term reliability of the pin-dot.

#### 3.2.3 Detection path

Fluorescence emission generated by molecules diffusing through the 8 excitation spots was collected by the objective lens. A dichroic mirror separated the scattered laser light from the dyes emission (Fig. SI-3). An additional long pass filter was inserted to eliminate any residual fraction of scattered laser light transmitted by the dichroic mirror. The 8–spot emission signal was focused by the microscope tube lens and relayed by a multi-camera port system (Tricam, Cairn Research Ltd., Kent, UK) to two 8-pixel SPAD modules [12], after spectral separation. Each module was individually aligned so that each pixel’s active-area received the emission signal from one of the conjugated excitation spots (Fig. SI-3).

The arrays geometry consisted of eight 50 µm-diameter pixels linearly distributed with a 250 µm pitch. The arrays, which have been described previously [10, 12], use a custom process developed by the Polimi group. This process allows obtaining arrays with low dark-counting rates (DCR) and higher PDE than arrays manufactured using CMOS processes (see for instance ref. [13, 14]). However, compared to the thick SPAD process (the detector technology used by the SPCM-AQR detectors of Excelitas or τ-SPAD detectors of Laser Components), the PDE of the custom arrays is about two-fold lower in the red part of the visible spectrum (>600 nm). Therefore, we expected a low sensitivity of the multispot system for low-FRET populations, characterized by very low photon counts in the red part of the spectrum (*i.e.* in the acceptor channel). A more detailed comparison of the different SPAD technologies mentioned above can be found in ref. [15].

#### 3.2.4 Multispot data acquisition

Data acquisition from the two SPAD arrays was performed as described in ref. [16]. Briefly, each SPAD’s TTL output was connected to a single digital input channel of a programmable board (PXI-7813R, National Instruments). The LabVIEW FPGA firmware uploaded on the board detects and time-stamps each signal with 12.5 ns resolution and associates it with its channel number, before transferring the data to a host computer running the LabVIEW acquisition software. Data was saved in binary format recording the time stamp and channel number of each photon in the order they were received, and displayed in real time as time traces of all channels for monitoring purposes. Raw data files were later converted in the Photon-HDF5 data format using the phconvert Python script [1]. These file are available online at URLs indicated Appendix 1.

#### 3.2.5 Alignment

##### 3.2.5.1 Excitation Path

When building the setup, before placing the L3 lens and the pindot filter, the laser needs to be aligned as in a standard confocal setup, using the LCOS as a mirror (the LCOS must be turned on and displaying a uniform black pattern). The expanded beam needs to be centered and aligned along the objective lens’ optical axis. As in standard confocal systems, this alignment can be achieve using a pair of mirrors which are enough to provide both tilting and lateral translation of the beam. The LCOS position should be aligned so that the multispot pattern would lay as close as possible to the peak of the Gaussian beam profile. One the beam is aligned the lens L3 can be inserted. This lens can be easily centered making sure that the confocal spot generated by the objective lens is in the same position after the L3 lens insertion. Finally, the pindot filter can be inserted at a focal-length distance after L3. In this configuration, when the pindot is aligned, it should block the laser excitation almost entirely. For fine alignment of the pindot, we suggest to use a high-concentration dye sample (e.g. 100 nM Cy3B solution) and a camera. Focusing several a few tens of microns inside the sample, it is possible to finely align the pindot by minimizing the emission intensity recorded by the camera. After this point, the multispot LCOS pattern can be activated. The same high concentration dye sample should show the multispot emission pattern. In principle is possible to translate the LCOS pattern to align the emission on the detector. However, this would bring the pattern away from the peak of the beam profile and away from L3 optical axis, causing uneven attenuations and aberrations. Therefore, it is better to align the SPADs instead, leaving the LCOS pattern centered.

##### 3.2.5.2 SPAD Array Alignment

The alignment procedure was performed using a concentrated (100 nM) Cy3B dye solution. Cy3B has a wide enough emission band to be detectable by both the donor and acceptor channels. During alignment the laser power was adjusted such as to achieve a high signal-to-noise ratio (SNR) on the SPADs, making the dark counting rates (DCR) of each pixel negligible.

As a preliminary step, the two detectors are manually aligned to the pattern by maximizing the signal of an 8-spot pattern (formed as described in the next section). After the two detectors are roughly aligned, we perform a fine-alignment procedure described below.

Focusing one detector, we perform a software-controlled cross-hair scan of a single-spot LCOS pattern across the approximate centers of each SPAD pixels and record the corresponding intensities traces from the SPADs. The scan pattern (a cross comprised of a horizontal and a vertical segment) provides intensity profiles which are used to identify the X-Y coordinates of each SPAD pixel. The same procedure is repeated for the second detector. From the coordinates of the 8 pixels in each detector, we compute the X-Y offset and the relative rotation of the two detectors. Next, we adjust the SPAD micro-positioners in order to reduce the offset in X-Y and rotation. The scan procedure is iteratively repeated getting new SPAD coordinates, and adjusting the alignment of one SPAD detector until the offset is minimized.

##### 3.2.5.3 LCOS Pattern Formation

Each excitation spot is generated by modulating the incident plane wave with the phase of a spherical (converging) wave. It is easy to derive the phase from geometric considerations (Fig. SI-4).

The relation between phase delay ∆*Φ* and physical displacement ∆*f* is:

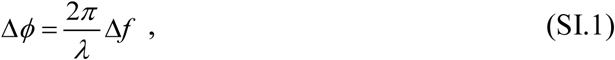

where *λ* is the wavelength. The phase delay to be applied to a plane wave in order to obtain a converging spherical wave is given by:

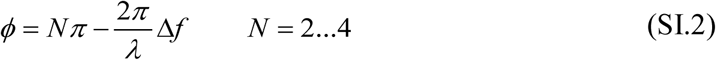

where *N* is an arbitrary constant, which is only used to ensure that the phase remains inside the LCOS dynamic range. To obtain a convergent spherical wave, ∆*f* should follow the profile of a sphere:

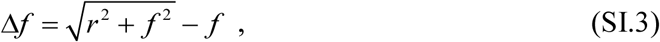

or, equivalently:

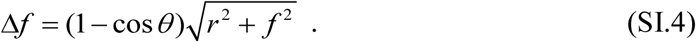

The choice between the two expressions is just a matter of implementation preference.

It is possible to derive a paraxial approximation of the previous formulas which only uses algebraic operations. Using the expansion:

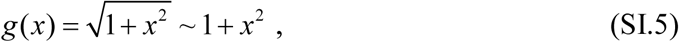

The following approximate expression for ∆*f* and *Φ* are obtained:

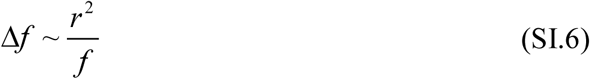

and:

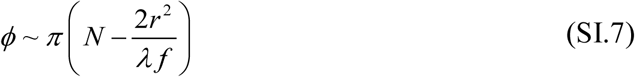

The latter expression is the one reported in [10].

To generate an array of spots, we synthesize an array of 8 adjacent “lens patterns” on the LCOS surface. Each “lens pattern” focuses a spot at a focal length from the LCOS (in our case 32 mm). The pitch of the lens pattern on the LCOS is equal to the pitch of the spots at a distance *f* from the LCOS. The spots are de-magnified by a recollimating lens L3 and the objective lens. Therefore, the pitch of the excitation spots in the sample is controlled by the pitch of the “lens pattern” on the LCOS. The emission pattern on the SPAD plane needs to have the same pitch as the SPAD array, which is 250 µm. This translates into a pitch of 4.1 µm in the sample (60X de-magnification). The overall pattern dimension in the sample is therefore of the order of 30 µm. From the sample to the LCOS, there is a magnification of (250/3 = 83.3), therefore the pitch on the LCOS is nominally 347.2 µm (or 17.4 × LCOS pixels). This value needs to be adjusted experimentally (usually ± 5 %) because of variations of lens focal length from nominal values and slight differences in the collimation of the incoming laser beam. The experimental pitch can be accurately estimated with the scanning procedure described in Section 3.2.5.2.

**Fig. SI-4:**
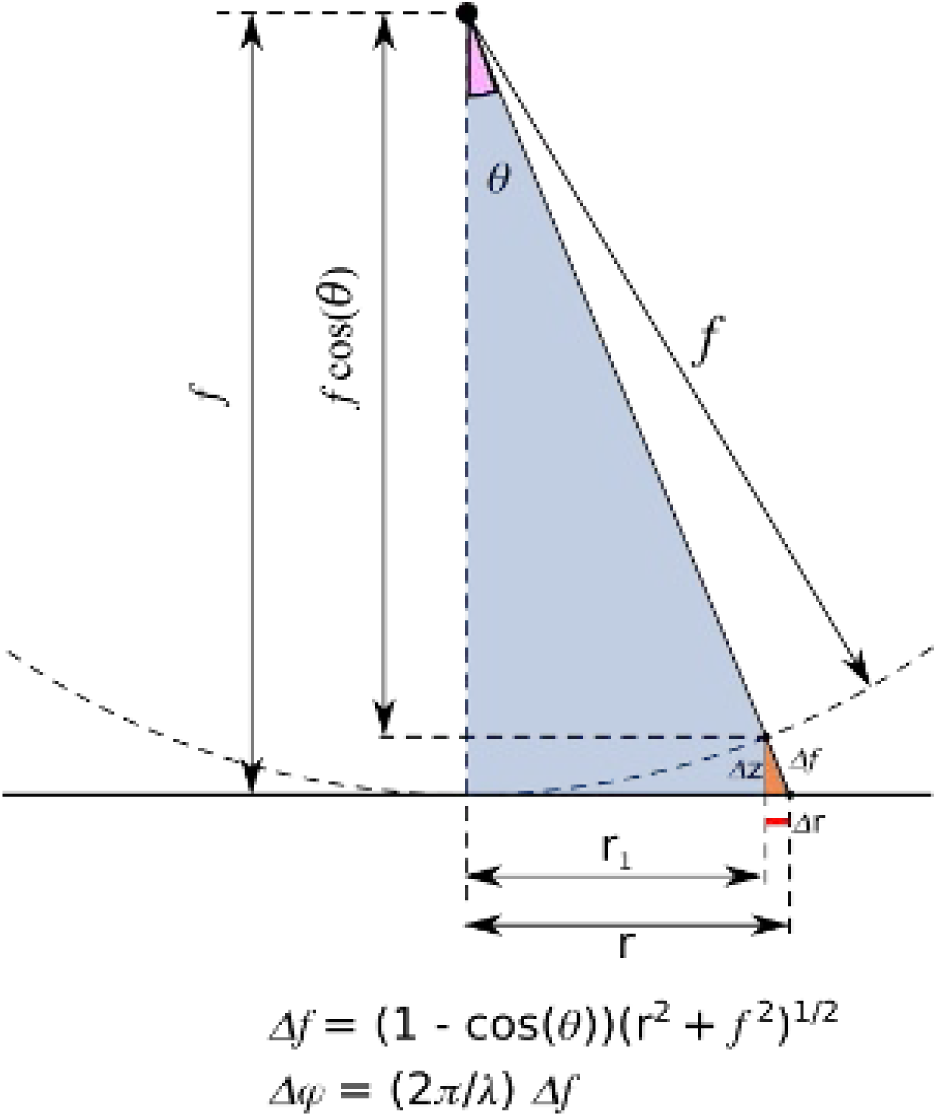
Geometrical construction illustrating how to compute the phase delay of a spherical wave. An incident plane wave propagating top to bottom can be focused at a distance *f* from the reflecting surface (e.g. the LCOS surface) if, after reflection, its wave-front (surface of equal phase) is transformed into a sphere. This happens if the phase is modulated with a radially symmetrical function with suitable dependence on *r*.

Overall the extension of the 8-spot pattern on the LCOS is less than 2.5 mm along the long axis and around 350 µm along the short axis.

Given the pitch and the focal length, the 8-spot LCOS pattern can be generated using equations (SI.1) and (SI.3). In particular, the coordinates of the centers of the *n* spots (in our case, *n* = 8) for a pattern centered around the origin (conventionally chosen as the LCOS center) can be expressed as:

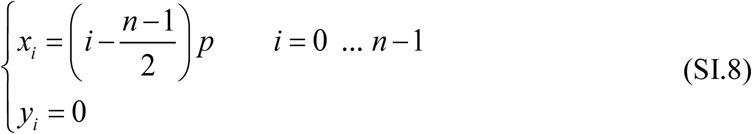

In practice, we need to translate this pattern to an arbitrary position *Δ* on the LCOS, and rotate it with respect to the pattern’s center. The rotational degree of freedom is needed to match the exact orientation of the detector array (we found that the rotational adjustment was around 1 degree). A rigid transformation (translation + rotation) is a linear operation that can be synthetically expressed as:

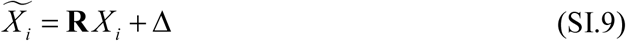

Where *X* = [*x*_*i*_, *y*_*i*_], *X*_*i*_ are the rotated coordinates and **R** is the rotation matrix:

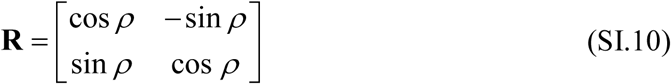

Finally, for each *i*, we evaluate function *Δϕ* (Eq. (SI.1)) centered around each spot position *X*_*i*_ in a region limited to a small square with side equal to the pattern pitch. The LCOS pixels outside this “lens patterns” are filled with a beam steering pattern, which is an alternation of horizontal lines with phase of 0 and π. A zoomed in image of a typical multispot pattern is shown in Fig. SI-5. Readers can find a complete Python implementation for generating multispot LCOS patterns with arbitrary input parameters in the LCOS_pattern folder of the notebooks archive (link).

**Fig. SI-5:**
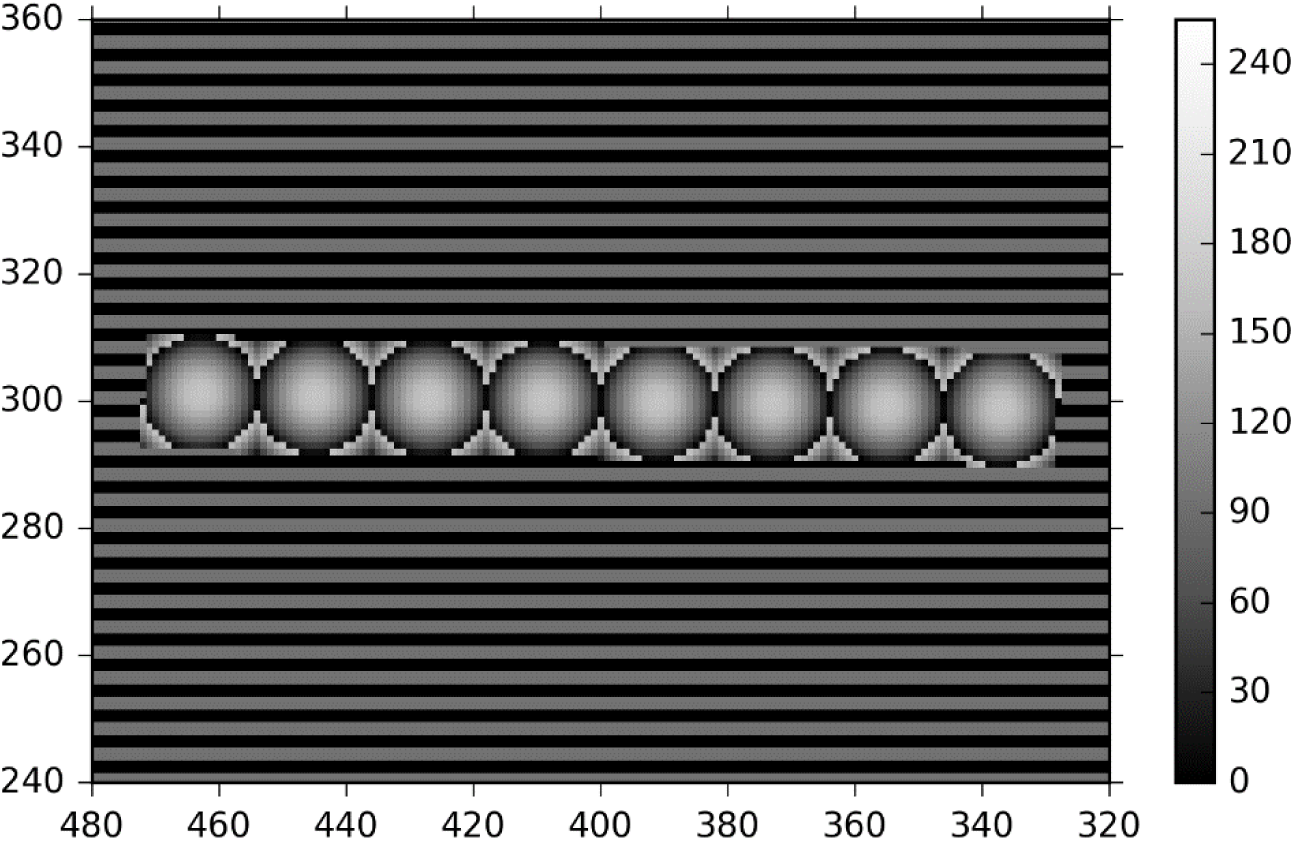
Zoomed in of a typical multispot pattern used to generate 8 excitation spots. The value of each pixel ranges between 0 and 255, matching the grey-scale image format required by the LCOS used in this work. The discontinuity (black to grey transition) inside each spot happens at points where the phase difference from the spot center is larger than 2*π*. The steering pattern, visible outside the lens pattern, is comprised of alternating stripes of phase 0 and *π*. Note that the pattern has a rotation angle of 1 degree applied to it.

## Appendix 4 Crosstalk Analysis

The amount of crosstalk between two SPADs measures the correlation between their signals appearing on top the expected correlation due to diffusion or photophysics, in the case of two detectors detecting the signal emitted from the same spot. Unwanted crosstalk between channels can happen due to different reasons:

- electrical signal pick-up
- emission point spread function (PSF) spillover
- intrinsic optical crosstalk

The first kind of crosstalk happens when the TTL pulse coming from one channel is picked up by another line, or triggers a pulse detection on another line. It is rare in well-designed setups and electronics.

The second kind of crosstalk can happen if the illumination/detection geometry (in particular the size of the image of each excitation spot) is such that a fraction of the signal emitted from one excitation spot, which should optimally confined to a dedicated SPAD, can be detected by a neighboring SPAD, for instance because the SPAD separation is comparable to the PSF extension. This was not the case in these experiments, where the separation between spots was about one order of magnitude larger than the PSF extension.

The third type of crosstalk is specific to SPAD arrays and is due to secondary photon emission during each avalanche. Those isotropically emitted photons can reach neighboring SPADs (either through a direct path or after reflection off the chip’s surfaces) and can trigger an avalanche in those SPADs [17].

The detailed description of crosstalk analysis of the SPAD arrays used in this study is beyond the scope of this article and will be presented elsewhere. The underlying phenomenon is now well understood and has been described in the literature [17, 18]. Here we merely provide the expression used to compute the crosstalk coefficient *l_AB_* between two SPADs A and B, based on the computation of the value of the cross-correlation function of the two signals at time lag 0, CCF_AB_(0):

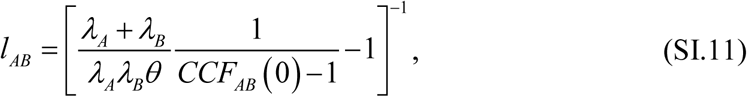

Where*, θ* is the resolution chosen to compute the correlation function and *λ_A_* (resp. *λ_B_*) is the average count rate measured in channel A (resp. B). The only subtlety in this analysis is the choice of the resolution parameter *θ*, which we briefly address here.

The minimum value is set by the timestamp resolution, imposed by the counting electronics. In these experiments, timestamps where obtained using a 80 MHz clock, defining a resolution of 12.5 ns.

While using this value in Eq. (SI.11) and calculating *CCF(0)* with this resolution provided crosstalk coefficients of the correct order of magnitude, it neglects the fact that quenching the avalanche in a SPAD takes a finite amount of time (typically a few tens of ns), during which the probability of secondary photon emission is not zero. During this time, a crosstalk avalanche can be triggered in a nearby SPAD. It is therefore important to compute Eq. (SI.11) using a resolution *θ* covering the time scale of the SPAD avalanche. We found that a value of 37.5 ns (3 times the clock resolution) was appropriate for this calculation (using *θ* = 50 ns provided identical results).

Eq. (SI.11) is equivalent to the expression provided in ref. [18], and can also be expressed in terms of the number of counts detected in each channel during the chosen integration time *θ* (*N_A_* and *N_B_*), as well as the number of coincident counts *C*, using:

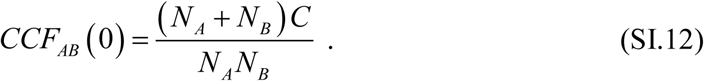

Tables SI-1 and SI-2 report the measured crosstalk values (in percent) for both SPAD arrays. Because of their extremely small values, crosstalk is therefore not a concern in this study.

**Table SI-1:** Crosstalk coefficients (in percent) for the donor channel SPAD array.

| SPAD # | 1 | 2 | 3 | 4 | 5 | 6 | 7 | 8 |
| --- | --- | --- | --- | --- | --- | --- | --- | --- |
| 1 | 0 | 0.11 | 0.13 | 0.12 | 0.12 | 0.11 | 0.09 | 0.13 |
| 2 | 0.11 | 0 | 0.15 | 0.16 | 0.16 | 0.13 | 0.12 | 0.13 |
| 3 | 0.13 | 0.15 | 0 | 0.15 | 0.15 | 0.17 | 0.13 | 0.26 |
| 4 | 0.12 | 0.16 | 0.15 | 0 | 0.15 | 0.17 | 0.17 | 0.26 |
| 5 | 0.12 | 0.16 | 0.15 | 0.15 | 0 | 0.17 | 0.19 | 0.28 |
| 6 | 0.11 | 0.13 | 0.17 | 0.17 | 0.17 | 0 | 0.18 | 0.32 |
| 7 | 0.09 | 0.12 | 0.13 | 0.17 | 0.19 | 0.18 | 0 | 0.46 |
| 8 | 0.13 | 0.13 | 0.26 | 0.26 | 0.28 | 0.32 | 0.46 | 0 |

**Table SI-2:** Crosstalk coefficients (in percent) for the acceptor channel SPAD array.

Table SI-1: Crosstalk coefficients (in percent) for the donor channel SPAD array.
| SPAD # | 1 | 2 | 3 | 4 | 5 | 6 | 7 | 8 |
| --- | --- | --- | --- | --- | --- | --- | --- | --- |
| 1 | 0 | 0.26 | 0.27 | 0.24 | 0.18 | 0.17 | 0.15 | 0.27 |
| 2 | 0.26 | 0 | 0.19 | 0.27 | 0.20 | 0.18 | 0.16 | 0.27 |
| 3 | 0.27 | 0.19 | 0 | 0.21 | 0.25 | 0.24 | 0.21 | 0.26 |
| 4 | 0.24 | 0.27 | 0.21 | 0 | 0.17 | 0.22 | 0.21 | 0.33 |
| 5 | 0.18 | 0.20 | 0.25 | 0.17 | 0 | 0.18 | 0.25 | 0.45 |
| 6 | 0.17 | 0.18 | 0.24 | 0.22 | 0.18 | 0 | 0.19 | 0.41 |
| 7 | 0.15 | 0.16 | 0.21 | 0.21 | 0.25 | 0.19 | 0 | 0.38 |
| 8 | 0.27 | 0.27 | 0.26 | 0.33 | 0.45 | 0.41 | 0.38 | 0 |

## Appendix 5 Photon Streams Definition

In µs-ALEX, photon streams are defined based on a histogram of the “reduced” timestamps of photons in each detection channel (D and A). The reduced timestamp *t*_*i*_ of a photon is simply its timestamp value *t_i_* modulo the alternation period *T*:

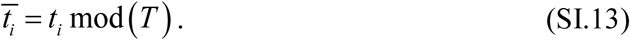

As shown in Fig. SI-6, the resulting alternation period histograms exhibits two regions with different count rates, one of which corresponds to the donor excitation period, the other to the acceptor excitation period.

**Fig. SI-6:**
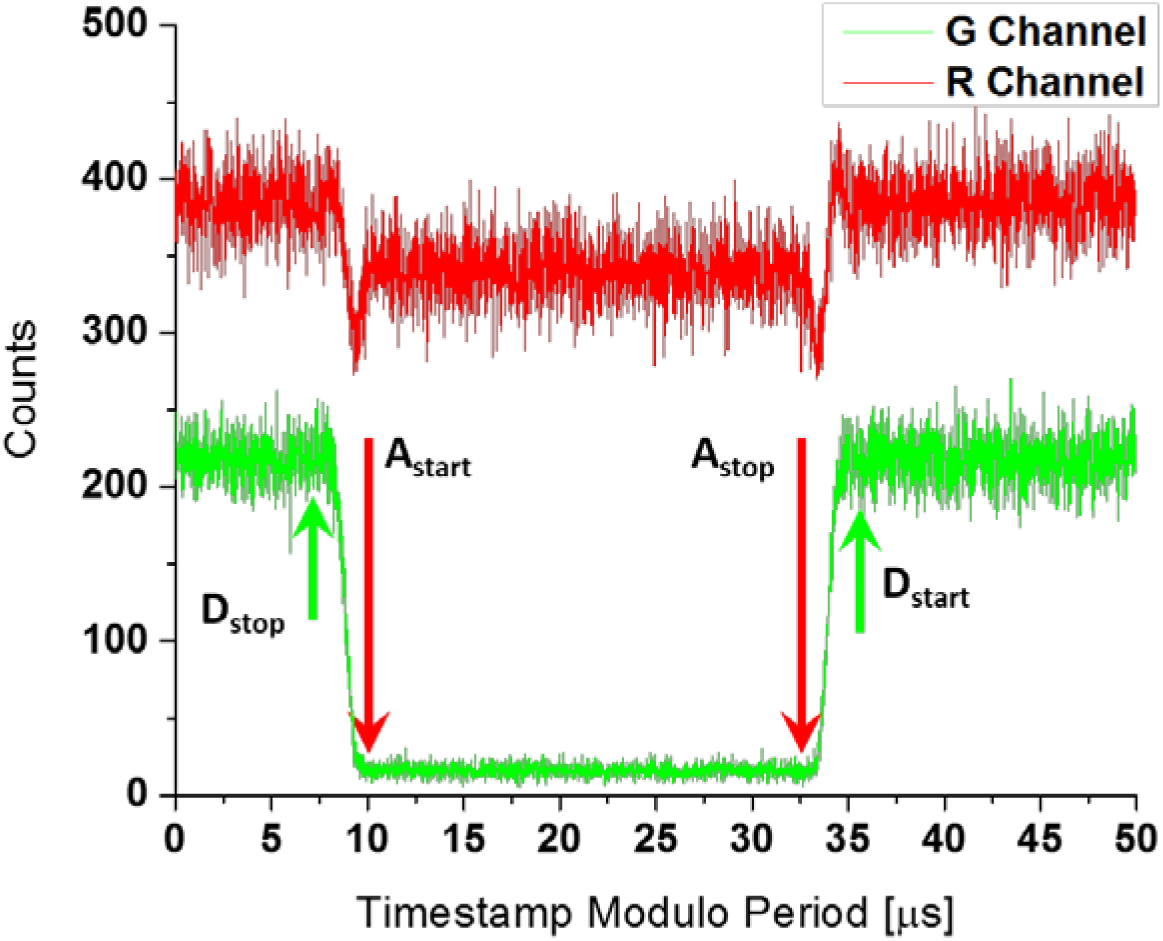
Definition of the donor and acceptor excitation periods within the alternation period in the 12d sample. Histograms of all photons timestamps recorded in the green (G) and red (R) channels are binned with 12.5 ns resolution. The acceptor excitation period is characterized by a lower photon count in the green channel.

The latter is easily identified as the period with almost no donor photon detection (the low detected count rate corresponds to the dark count of the detector and residual background in that channel). Due to the finite response time of the AOM, there are transition regions between donor and acceptor periods, during which both laser lines are partially transmitted. For this reason, two ~2 µs “gap” regions around the donor to acceptor and acceptor to donor transitions are usually rejected. This preprocessing step allows to define the 4 base streams of timestamps: D_ex_D_em_ (D excitation, D emission channel stream), D_ex_A_em_ (D excitation, A emission channel stream), A_ex_A_em_ (A excitation, A emission channel stream) and A_ex_D_em_ (A excitation, D emission channel stream). These streams identify photons detected during the D or A excitation periods (D_ex_ or A_ex_) by the D or A detection channels (D_em_ or A_em_).

In practice, reduced timestamps might be offset by a few microsecond (*t_0_*) with respect to the alternation period “edges” (the time when one laser is turned off and the other on), as illustrated in Fig. SI-6. In this case, it might be easier to define:

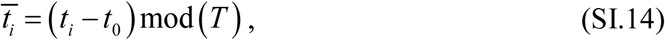

which allows defining each excitation period by a simple condition such as:

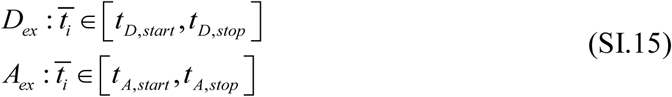

## Appendix 6 Background Rate Calculation

### 6.1 Introduction

#### 6.1.1 Sources of background

Sources of background in single-molecule experiments are multiple and commonly include Rayleigh and Raman scattering, scattering or fluorescent impurities, out-of-focus molecules and detector noise. The first two can be practically eliminated by a proper choice of filters. Buffer contributions can consist of single-molecule-like bursts (which will be treated as single-molecule bursts) or uncorrelated background signal, and can be minimized using ultrapure reagents and filtration. The detector dark count rate is easily characterized but can be non-Poissonian. In ideal situations, background rates can be reduced to less than 1 kHz.

#### 6.1.2 Background rate calculation for each measurement

While it is possible to use a buffer sample and use the measured count rates for that sample as background rates for subsequent samples using the same buffer and experimental conditions, this procedure ignores the contribution of out-of-focus molecules present in most samples. Therefore, a procedure extracting the relevant background rates from each sample file is preferable. A simple approach consists in looking at the inter-photon delay distribution, *φ(τ)*, for each stream.

#### 6.1.3 Inter-photon distribution

For photons generated by a hypothetical background Poisson process, this distribution will be exponential. In the case of single-molecules diffusing through a continuous (non-alternated) excitation spot, this distribution is complex but can be approximated as the weighted sum of two terms, one of which is exponential [19]:

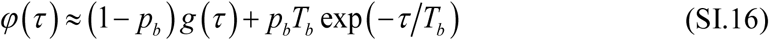

In this expression, *g(τ)* is the inter-photon delay distribution for a single molecule diffusing through the excitation volume (~ *τ*^*-3/2*^ for a 3-dimensional Gaussian PSF, see ref. [20]) and *T_b_* is the mean time between bursts. The first term dominates at short time scales, while the second dominates at long time scales. Since this expression is derived in the absence of external background rate, this means that a background-free single-molecule signal generates an effective Poisson background rate proportional to the concentration, *b* = 1/*T_b_*, to which any additional background source (buffer, detector, etc.)^1^ will be added in real experiments [19].

The background rate used for correction is defined as the exponential rate extracted from the long time scale behavior of the inter-photon delay distribution. This rate can be estimated without computing the inter-photon delay histogram (which can be time consuming for large data sets), using the fact that the maximum likelihood estimator (MLE) of the rate parameter of an exponential distribution is simply equal to the inverse of the mean of that distribution:

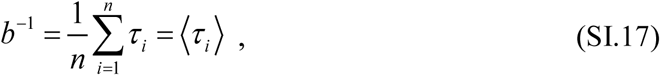

where the *τ_i_*’s are the inter-photon delays. Note that the MLE estimator is only one of many possible estimators. Alternatives are the minimum variance unbiased estimator (MVUE) or the estimator which minimized the mean square error. All these estimators only differ by a multiplicative factor depending on *n*. Because the background rate is estimated with a number of inter-photon delays much larger than 1,000, the difference between the various estimators is in general negligible.

As discussed previously, the inter-photon delay distribution is not exponential, only its large time scale behavior is (Eq. (SI.16)). Therefore, only part of the distribution should be used to estimate the background rate. The MLE of the rate parameter *b* of such a truncated exponential distribution (*τ_i_* > *τ_min_*) is simply given by:

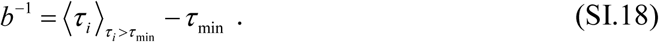

Choosing *τ_min_* involves a trade-off: on one hand, using only the longest time scales of the distribution would result in poor statistics. On the other hand, trying to improve the statistics by extending the range of included time scales towards small values would result in the incorporation of (short) inter-photon delays from within single-molecule bursts, biasing the background rate estimates (towards larger values). Optimization of the lower inter-photon delay used for background rate estimation can be performed systematically, but its study is beyond the scope of this paper. For the experiments in this paper, we computed the *τ_min_* threshold with a previously described, simple iterative algorithm [21]. While it is possible (for example using the FRETBursts software [21]) to perform a global optimization in order to find the optimal *τ_min_* for background estimation, this was not necessary as the procedure only yielded negligible differences.

**Fig. SI-7:**
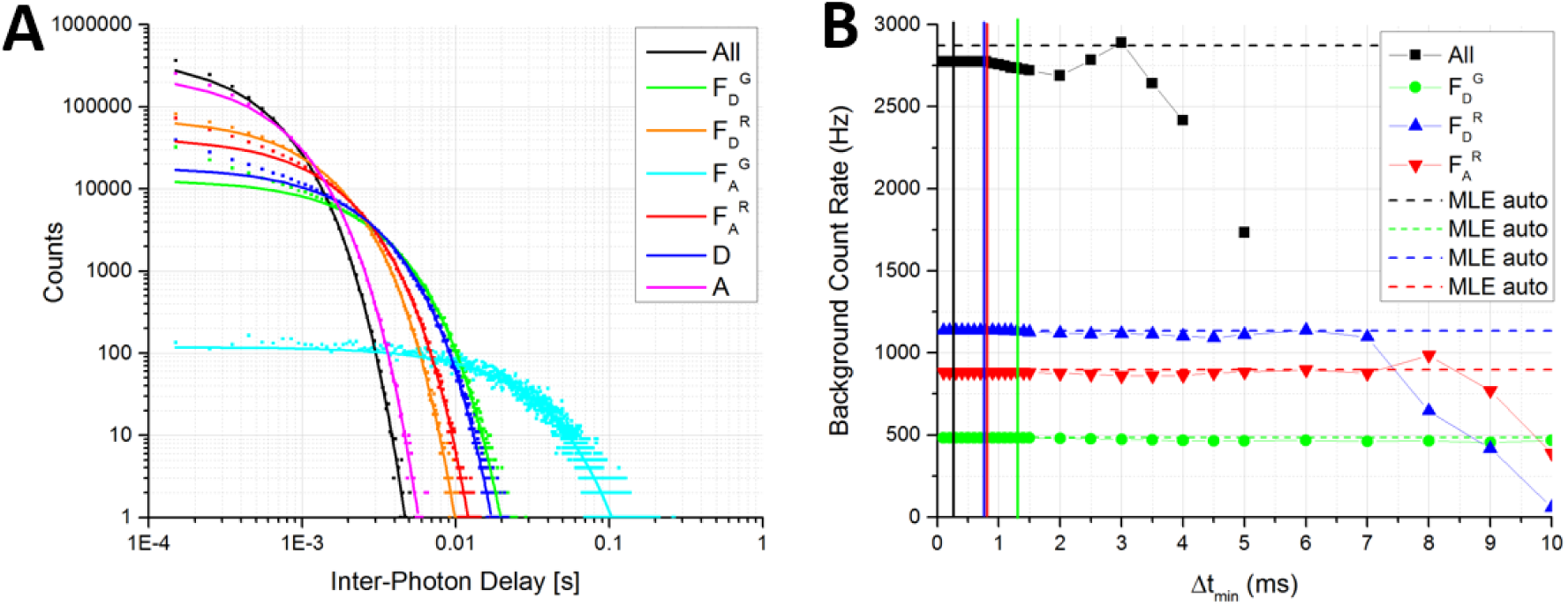
Robust Fit of Background Count Rates. (**A**) Inter-photon delay histograms (bin = 100 µs) for different photons sets of file 006_dsDNA_7d_green100u_red40u.hdf5 (points) and their robust fit with a single exponential (curves). D (resp. A or All) indicates donor (resp. acceptor or all) photons, irrespective of their time of emission and detection channel. *F*_*D*_^*G*^ = Dem, *F*_*D*_^*R*^ = Aem, *F*_*A*_^*R*^ = A_ex_A_em_. Fits were performed with an automatically computed Δt_min_ for each set (Δt_min_ = 1/<count rate>, where <count rate> is the mean count rate for that set, computed over the whole experiment). (**B**) Dependence of the background count rate estimated by robust fit of the tail of the inter-photon delay histogram on the minimum interval. The points correspond to the fitted rate obtained using the specified minimum inter-photon delay Δt_min_ for 4 representative sets of photons. Fits were performed using a single exponential model, statistical weights and the bisquare algorithm implemented in LabVIEW 2015 with default parameters. The vertical lines indicate the location of the automatically determined Δt_min_ for each set of photons. The dashed horizontal lines correspond to the MLE result for each photon set, using the automatically determined Δt_min_ for each set. Except for the “all photons” set, the agreement between MLE and fitted background rates is excellent. Notice how the fitted count rate deviates from the automatically determined value only for very large Δtmin and is otherwise practically constant in the range 0 – 1 ms for the “all photons” set and well past 5 ms for the other sets.

An alternative approach (resulting in similar results) consists in performing a robust fit (e.g. a non-linear least square fit using Tukey’s biweight [22], also known as a bisquare fit) of the inter-photon delay histogram of each stream, for inter-photon delay larger than a minimum inter-photon delay λ_min_. The definition of λ_min_ is such that the resulting background count rate is, to a good approximation, not dependent on it. An example of definition of λmin is the average count rate in the photon stream (as illustrated in Fig. SI-7), but the advantage of this robust fit approach is that the result does not depend on the value chosen for λ_min_ over a large range of values. Its disadvantage is that it requires calculating inter-photon delay histograms, which can be memory and CPU intensive for large files (the fit itself is fast).

### 6.2 Variable background rates

If the background rates vary during the experiment (for instance due to focus drift, evaporation or other reasons), background estimation needs to be performed piecewise over time windows during which rates can be considered as approximately constant, but long enough to ensure proper estimation of this “local” background rate. In this work, a few files exhibited varying background rates justifying using this approach (Fig. SI-8A & B). Note that for the single-spot µs-ALEX measurements, the background levels were low enough that even a significant variation did not result in major differences between results obtained with or without time-dependent background rate calculation (Fig. SI-8C-F). Because background levels were larger in multispot measurements, and for consistency, time-dependent background rates (typical window size: 30 s) were used.

**Fig. SI-8:**
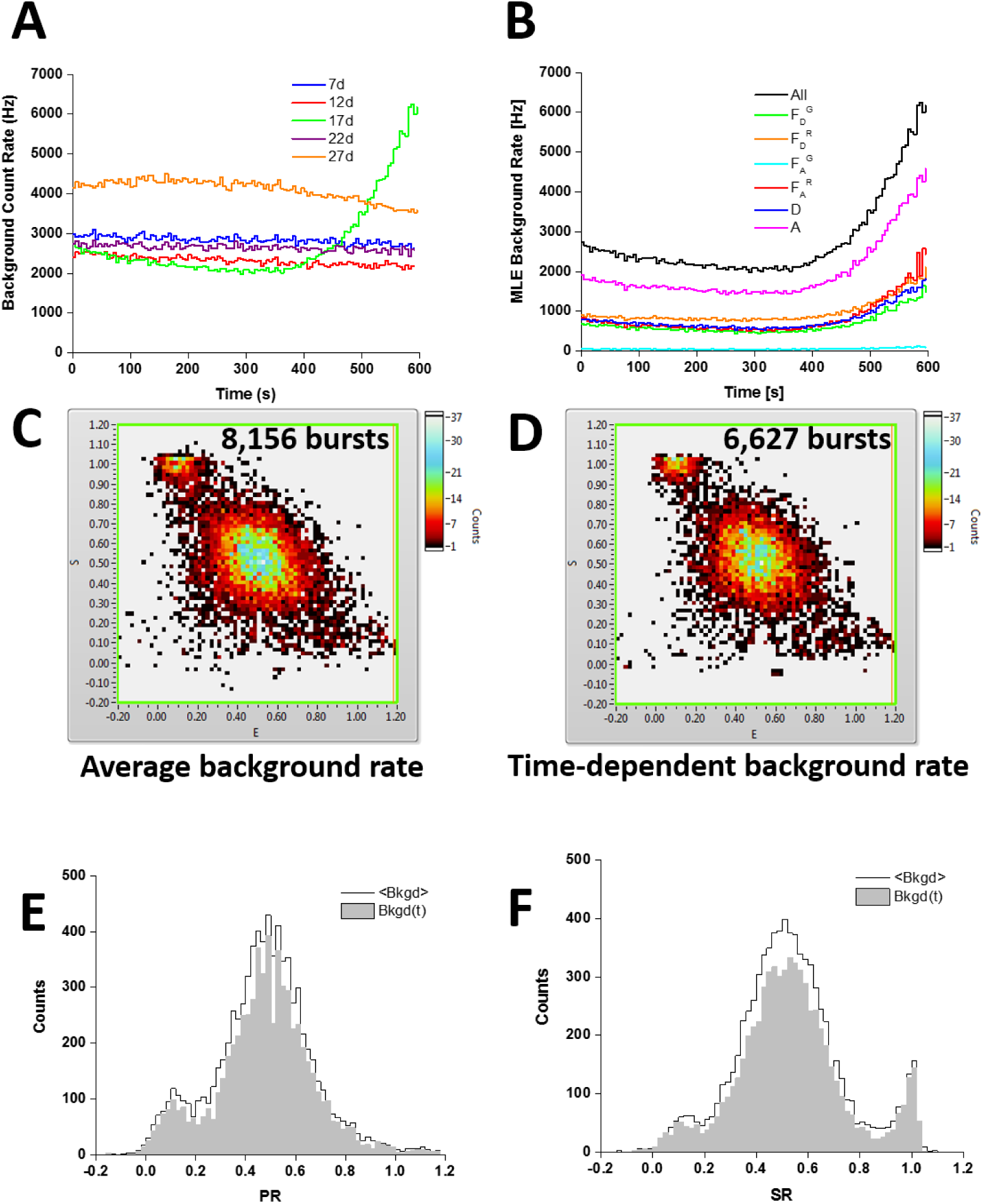
Time-Dependent Background Rate Fit. (**A**) Time-dependent maximum likelihood estimates (MLE) of the total background rates for all single-spot µs-ALEX measurements. The MLE calculation was performed on consecutive windows of 5 s duration, using the automatically determined *Δt_min_* for each set (*Δt_min_* = 1/<count rate>). The large variation for sample 17d is further studied in the other panels. (**B**) Time-dependent MLE of the background rates of different photons streams of the 17d sample. Notice how the background rate first decreases, then increases significantly during the last third of the measurement, most likely due to evaporation. (**C**) ALEX histogram (background corrections only) of the 8,156 burst of total size S > 30 obtained when using constant mean background rates for each photon set throughout the whole acquisition. (**D**) ALEX histogram (background corrections only) of the 6,627 burst of total size S > 30 obtained when using time-dependent background rates for each photon set (as computed in A). (**E**) Proximity ratio histograms of all bursts in C (black) and D (grey). Notice that, while the number of detected bursts is smaller when using a variable background rate, the characteristics of each sub-population are barely affected by the difference in background rates. (**F**) Stoichiometry ratio histograms of all bursts in C (black) and D (grey). As for the proximity ratio histogram, notice that the characteristics of each sub-population are barely affected by the difference in background rates.

### 6.3 Special consideration for µs-ALEX

A final comment is in order regarding µs-ALEX background analysis. Due to laser alternation, photon streams are periodically interrupted. Therefore, the photon and background rates within a stream periodically drop to almost zero (in practice, down to a value close to the detector dark count rate). The background rates fitted as described, are therefore rates averaged over the whole alternation period.

The “instantaneous” background rate alternates between a high value (*hb*_*X*_^*Y*^, *h* > 1, when excitation laser X is on) and a low value ( *lb*_*X*_^*Y*^, *l* < 1, when excitation laser X is off), where *b*_*X*_^*Y*^ is the background rate in channel Y during excitation period X (X and Y are either D or A). The average background rate is a weighted sum of these two rates:

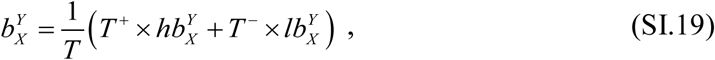

where *T*^*+*^ (resp. *T*^*-*^) is the duration of the X laser-on (resp. -off) period, and *T*^*+*^ + *T*^*-*^ = *T*, the alternation period (generally of the order of a few tens of µs)^2^.

Correspondingly, we can divide a burst duration into alternate X laser-on segments, *{δ T_i_^+^}_i=1,…,p_* and X laser-off segments, *{δ T_i_^−^}_i=1,…,m_*. The expected total background counts during the burst duration are thus:

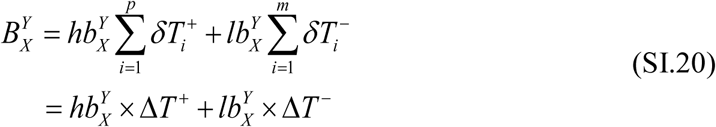

where *ΔT*^*+*^ (resp. *ΔT*^*-*^) is the total duration of X laser-on (resp. -off) periods during the burst.

We can therefore rewrite the background correction term used in this work as a function of the expected background counts as:

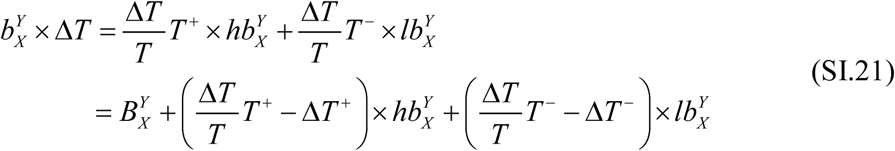

The last two terms are equal to zero when the burst duration *ΔT* is a multiple of the laser alternation period *T*: *ΔT* = *nT* (in other words, the background correction term is equal to the true estimation), since:

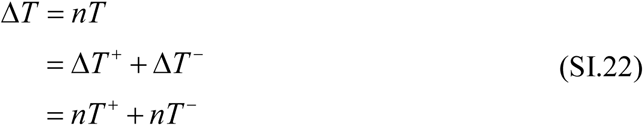

In the special case where the burst duration contains exactly one more complete on-period than off-period (*p* = *m* + 1):

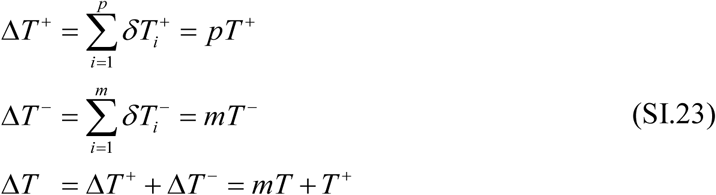

We can thus rewrite:

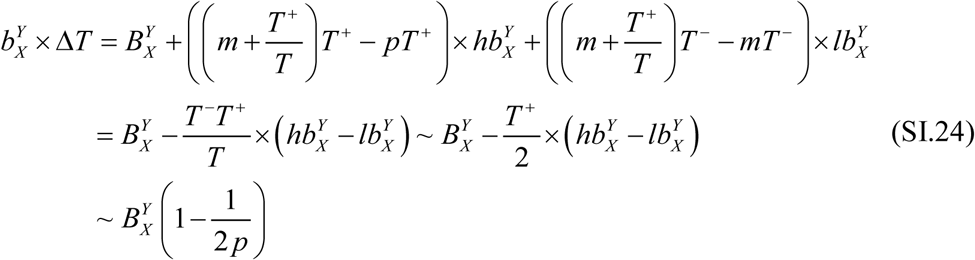

where the last two approximations use the fact that the on and off durations are approximately equal to one half the total alternation period *T*. The difference between the background correction term used in this work (the left hand side term) and the “true” background contribution, *B_X_^Y^*, is thus equal to a fraction of the correction term, and is increasingly negligible for longer burst duration. Bursts with duration comparable to the alternation period contain in general very few counts and are not normally selected for further analysis, justifying using the background correction term *b*_*X*_^*Y*^ × ∆*T* in all cases.

## Appendix 7 Details on Burst Search

### 7.1 Photon Streams

The photon stream is simply the set (or sets) of photons used for the search. The simplest choice consists in using all photons (“all photons burst search”, APBS) [23] but others can be defined to focus the analysis on specific species (footnote ^3^).

While different sets of photon streams can in principle be chosen to identify different types of molecules during burst search, it is often best to perform a generic single-molecule burst search (*e.g.* “all photons burst search”, APBS) and perform the burst selection at a later stage, as discussed in the next subsection. Most analyses presented in this work used APBS (unless specified otherwise), after we verified that the final results were not affected by this choice.

### 7.2 Burst Threshold

#### 7.2.1 Standard Definition of the Burst Threshold

Bursts are defined by their start and end counts, identified respectively as counts in the stream where the observed local count rate, *r_m_*(*t_i_*), computed over *m* consecutive counts starting with time stamp *t_i_*:

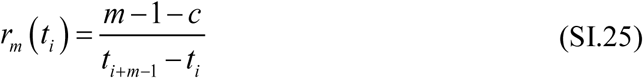

changes from “likely to be due to background only” to “unlikely to be due to background only”, and vice versa [15, 24, 25]. The role of parameter *c* in the previous equation, which was historically set to 0, is discussed in Section 7.2.2.

Burst searches using a minimum SBR criterion compare the value of Eq. (SI.25) to a threshold value defined as *F* times the local background count rate *b*(*t_i_*) [24]:

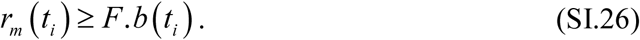

If the background count rate changes over time, the threshold also changes, guarantying a minimum SBR ≥ *F* – 1.

While such a criterion is appropriate for single-spot µs-ALEX experiments of samples characterized by similar background rates, molecular brightness and excitation intensity, it might not always be ideal. For instance, using a minimum SBR criterion might affect the comparison of multiple experiments, if the background rate varies significantly from experiment to experiment. In particular, the large dark count rates of some of the detectors used in our multispot experiments, resulted in very different background rates among spots and detection channels. In these cases, a fixed minimum count rate criterion is preferable, as discussed in Section 4.2.3 of the main text and in Section 7.4 below.

#### 7.2.2 Other Definitions of the Count Rate

The family of count rate estimators defined in Eq. (SI.25) has different properties for different values of *c*. For example, the MLE estimator is obtained for *c* = 0, the minimum variance *unbiased* estimator is obtained for *c* = 1, the estimator minimizing the mean square error is obtained with *c* = 2 while *c* = 1/3 yields an estimator of the median count rate (see this notebook for details).

To verify that the case *c* = 1 yields an unbiased estimator of the rate for a Poisson process, consider *m* consecutive photons generated by a Poisson process with rate *λ*. Let *t_i_*, *t_i_+1*, …, *t_i+m-1_* be the timestamps of the *m* photons, and:

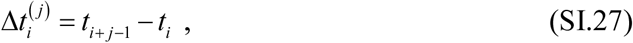

the delay between the last and first photon in a bunch of *j* successive photons. It is well known that the probability distributions of these quantities are Erlang distributions:

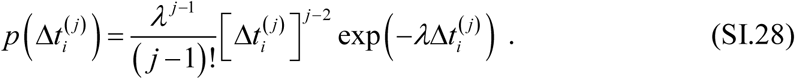

Using this formula, the mean value of the inverse of *t_i+m-1_* - *t_i_* (from which the count rate is calculated) is:

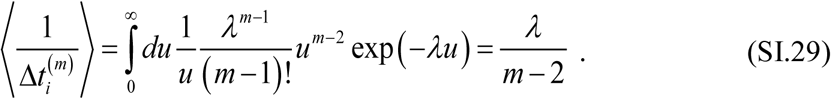

Consequently, to define a count rate from *m* consecutive timestamps in a manner resulting in an average count rate equal to the actual count rate *λ* (unbiased estimator), the following definition is needed (*c* = 1):

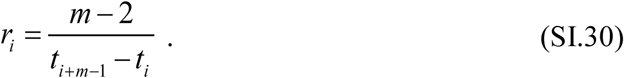

The exact version of the rate estimator (i.e. the value of *c*) used for burst search or burst statistics analysis is not critical, as long as the analysis is carried out consistently, using the same definition. Changing (*e.g.* increasing) *c*, while leaving all other burst search parameters unchanged, will change (*e.g.* decrease) the effective threshold used during bursts search. Note also that Eq. (SI.26) can be equivalently rewritten as a condition on *t_i+m−1_ - t_i_* (the difference between the last and first timestamps in the current set of *m* counts):

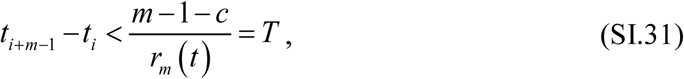

where *T* in the previous equation corresponds to the *T* parameter in the original publication introducing burst search [24], with the definition:

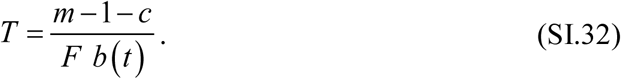

### 7.3 Burst Fusion

A typical burst search performed as described above, yields distributions of burst sizes, burst durations and burst separations which are generally monotonically decreasing. Burst separations, in particular, exhibit two general characteristic time scales: a short time scale corresponding to individual molecule exit from and reentry in the excitation spot, and a longer time scale corresponding to the typical interval between different molecules entering the excitation spot. The latter time scale decreases as concentration increases, while the former scales inversely to the molecule’s diffusion coefficient. Some analyses may benefit from considering bursts separated by short time intervals as independent (see for instance ref. [26]).

In some other cases, it might be preferable to fuse consecutive bursts separated by less than some minimum time, in order to increase the photon statistics of the resulting burst. The counts detected between the two fused bursts are mostly comprised of background, but are taken care of by background subtraction. It is worth mentioning that, even with background correction, the noise associated with these mostly background gap periods is added to the fused burst signal.

All results discussed in this paper were obtained without any burst fusion.

### 7.4 Burst Search Influence on Burst Statistics

The impact on single-molecule analysis of the burst search criteria is a common problem in single-spot measurements. In the case of multispot experiments, an additional complexity arises from the actual differences between each single-spot experiment performed in parallel. Some of these differences are discussed in the main text (Section 4.3). Here, we examine their influence on burst search results by focusing on the variations of a few observables as a function of burst search parameters. The effect of burst selection criteria on these observable is of course important as well, and needs to be considered carefully. The example of the influence of the minimum burst size is discussed in Appendix 8. Details on the definition of the observables discussed next can be found in Appendix 14.

By definition of a burst, the burst count rate (as calculated with Eq. (SI.30)) is initially low, increases and eventually decreases at the end of the burst (with possible fluctuations in between). Therefore, considering a particular burst, increasing the count rate threshold defining the beginning and end of the burst, will reduce its duration (the burst will start later and end earlier, everything else being equal) as well as reduce its background-corrected size. However, this should not affect the peak detected count rate during the burst, since the peak is typically attained well within the burst and not close to its beginning or end.

Such an increase of the effective count rate threshold is exactly what happens when a fixed set of search parameters (*m*, *F*) is used and a larger background rate is encountered in one spot compared to another. In other words, using the same SBR criterion for all spots will result in reduced burst duration and size for spots characterized by a larger background rate (a significant part of which is due to detector dark counts in our case). The peak burst count rate of identical species, however, should not be affected, provided all other acquisition parameters are identical (excitation intensity, alignment and molecule brightness).

**Fig. SI-9:**
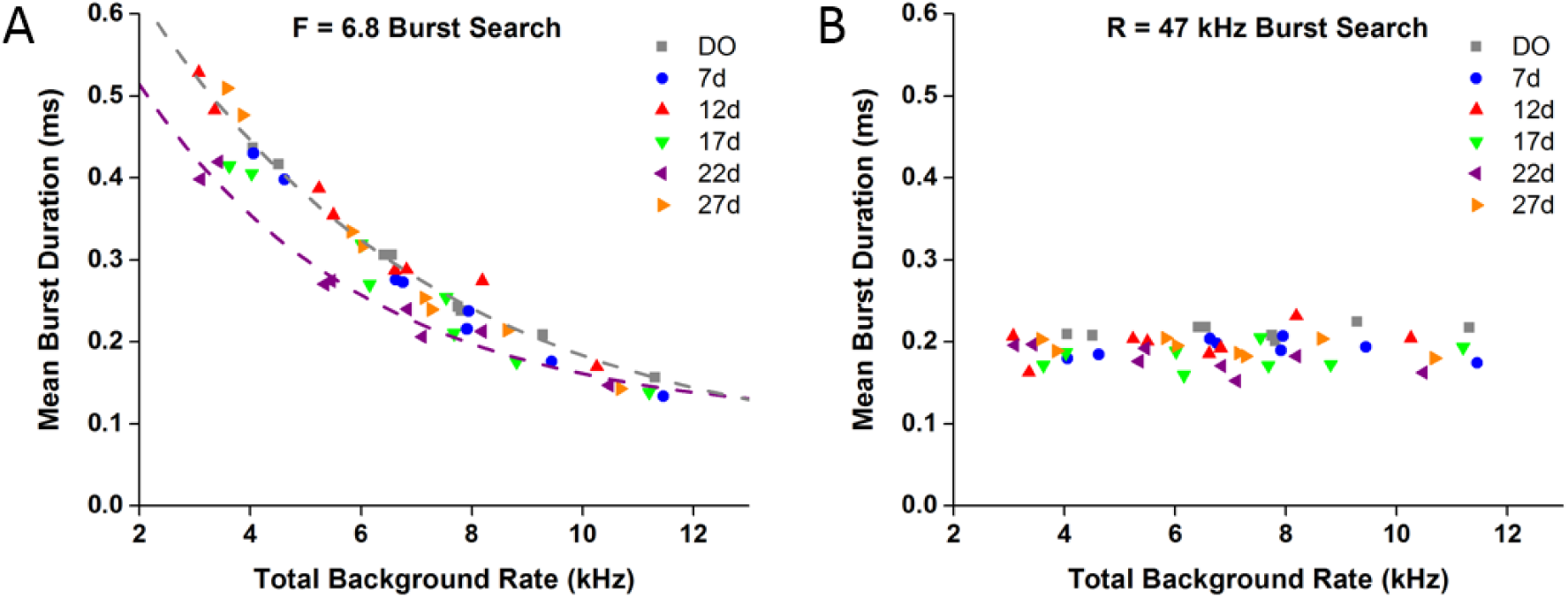
Mean burst duration dependence on total background rate. The mean duration of bursts with brightness-corrected size larger than 30 was computed for all samples (each sample represented with a different color) and each measurement spot within the samples, using two types of burst search. **(A)** *m* = 5, *F* = 6.8. **(B)** *m* = 5, fixed burst threshold rate *R* = 47 kHz. When a constant SBR search is performed (A), the mean burst duration is smaller when the background rate is larger, but does not depend on the sample or spot within a sample, provided the experimental conditions are similar (excitation intensity, alignment, molecular brightness, etc.). Note that the 22d sample trend (purple dashed curve) departs from the trend of all other samples (grey dashed curve), consistent with the observed shorter diffusion time measured for this sample.

As shown in Fig. SI-9A, the first conclusion is supported by our measurements. The average burst duration (irrespective of the sample or spot considered) falls on a universal decreasing curve when a constant minimum SBR criterion is used to detect bursts (*m* = 5, *F* = 6.8). Note that sample 22d, for which FCS analysis detected a shorter diffusion time, is characterized by noticeably smaller average burst durations, confirming the existence of a difference in the experimental conditions for this experiment. By contrast, using a fixed minimum count rate threshold to detect bursts (Fig. SI-9B, *r_min_* = 47 kHz), results in a mean burst duration which is independent of the sample or spot studied, as expected. The mean burst duration is also smaller, because the minimum count rate chosen (*r_min_* = 47 kHz) is close to the largest minimum count rate used across all samples and all spots in Fig. SI-9A. The same conclusion is illustrated on Fig. SI-10, where the complete distributions of burst duration and separation for all spots and all samples are represented. These distributions superimpose only when a *r_min_* = 47 kHz search is performed, while they are offset with respect to one another if a constant SBR search is used.

**Fig. SI-10:**
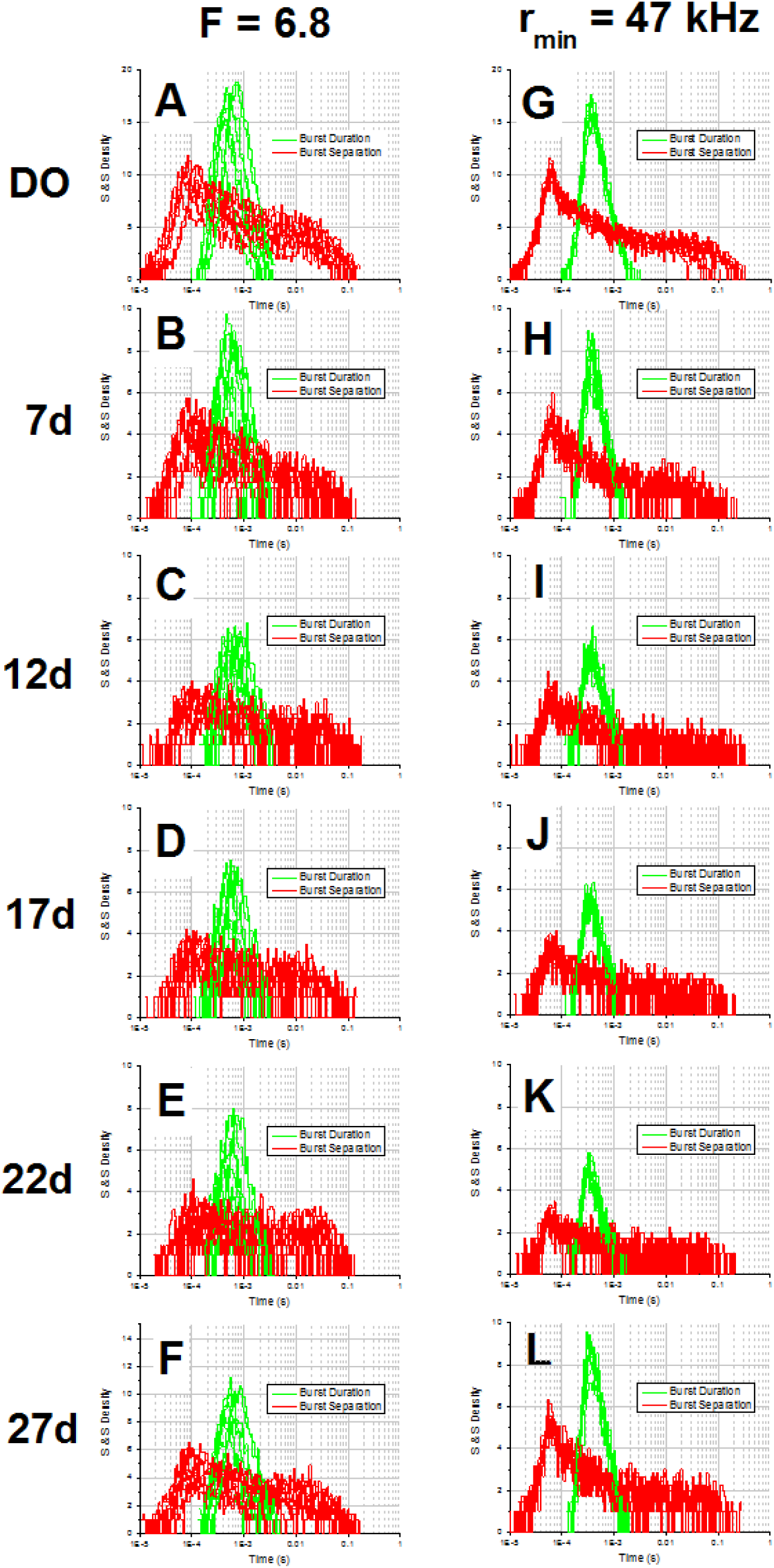
Burst Duration and Separation Distributions. Two types of burst search were performed on the 6 samples studied in this work (donor only: DO, doubly-labeled: 7d to 27d): (**A-F**) a constant (*m*, *F*) = (5, 6.8) search, adjusting the burst count rate threshold to the measured sample/spot background rate (left column), and (**G-L**) a constant burst count rate threshold (*r_min_* = 47 kHz) corresponding to 6.8 times the largest Donor + Acceptor channel dark count rate among all spots. After selection of all bursts with a minimum brightness-corrected size

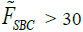

, the burst duration (green plots) and separation (red plots) distribution for each spot were represented using a log-linear representation introduced by Sigworth & Sine [27]. For the spot-dependent search (left column), both distributions clearly depend on the spot studied, while in the other case (constant threshold), the burst duration PDFs of all spots are essentially identical and very similar from sample to sample. A similar observation holds for the burst separation PDFs, with the exception of the long time scale tail, which is underpopulated for spots with large background rates, due to the culling of bursts with insufficient sizes.

The predicted dependence of burst size on background rate, using a minimum SBR criterion (*m* = 5, *F* = 6.8), is also clearly visible in Fig. SI-11A. The larger the background rate, the smaller the mean burst size computed over all detected bursts (filled symbols). Indeed, a larger background rate results in the loss of smaller bursts during the search. This dependence is much more pronounced if small bursts are eliminated by imposing a minimum brightness-corrected burst size selection criterion (*F_SBC_* ≥ 30, open symbols).

**Fig. SI-11:**
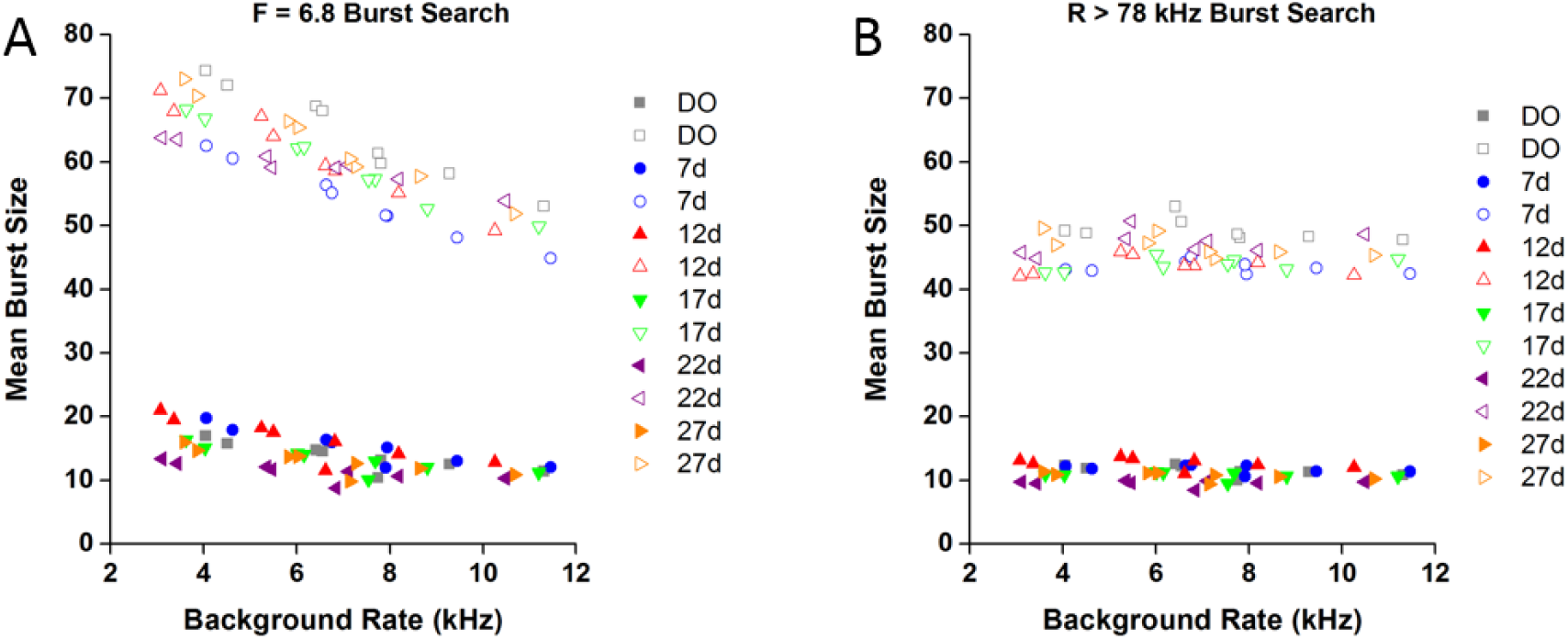
Mean brightness-corrected burst size dependence on total background rate. The mean brightness-corrected size (*F_SBC_*) of bursts was computed for all samples (each sample represented with a different color) and each measurement spot within the samples, using two types of burst search. **(A)** *m* = 5, *F* = 6.8. **(B)** *m* = 5, fixed burst threshold rate *r_min_* = 78 kHz. Two values are reported for each spot and each sample: the mean *F_SBC_* of all burst (plain symbols) and the mean *F_SBC_* of bursts with *F_SBC_* ≥ 30 (open symbols). The latter is larger because small size bursts are not included. When a constant SBR search is performed (A), the mean burst size decreases when the background rate increases, but does not depend on the sample or spot within a sample, provided the experimental conditions are similar (excitation intensity, alignment, molecular brightness, etc). If a common minimum count rate is used for burst search (B), the mean *F_SBC_* does not depend on background rate, spot or sample.

By contrast, using a fixed minimum count rate threshold (Fig. SI-11B, *r_min_* = 78 kHz), results in a mean burst size which is independent of the sample or spot considered. By limiting the analysis to bursts whose brightness-corrected size is larger than a minimum value (set here to 30), the mean burst size naturally increases (open symbols), but remains independent of which sample or spot is considered.

In conclusion, the type of burst search used during analysis can have a significant influence on some burst statistics. This influence needs to be thoroughly characterized when comparing data from difference measurements, or, as in the case of this study, data obtained from different spots during the same experiment and with the same sample.

## Appendix 8 PR and SR Histograms

### 8.1 Standard Histograms^4^

Background correction of raw burst counts *F*_*X*_^*Y*^ (counts collected during X-excitation in the Y-channel) is obtained by subtracting *b*_*X*_^*Y*^ × ∆*T*, where *b*_*X*_^*Y*^ is the photon stream’s background rate computed as described in Appendix 6 and ∆*T* = *t_e_* – *t_s_* is the burst duration, where *t_s_* is the burst start photon timestamp and *t_e_*, the burst end photon timestamp:

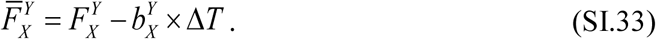

Note that Eq. (SI.33) uses the total duration of the burst and the average background rate for the selected photon stream. As discussed in Appendix 6.3, this formula is appropriate only for bursts long enough compared to the alternation period, which is the case in our analysis.

In the following, all overlined quantities are background-corrected quantities.

A useful quantity is the total background-corrected burst size:

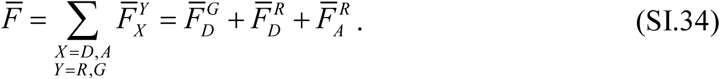

An alternative quantity, convenient in the absence of acceptor excitation laser, is the donor excitation background-corrected burst size:

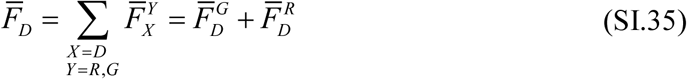

From the background-corrected burst observables

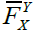

, two important quantities can be computed:

- the *proximity ratio*, *PR*, defined as:

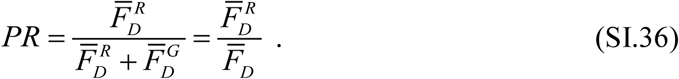

- the *stoichiometry ratio*, *SR*, defined as:

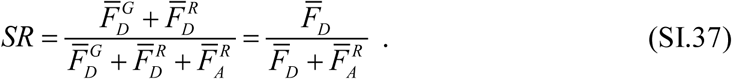

The stoichiometry ratio requires two laser excitations, and allows distinguishing between D-only labeled molecules (characterized by *SR* ~ 1), A-only labeled molecules (characterized by SR ~ 0) and doubly-labeled molecules (characterized by *SR* ~ 0.5).

*PR* and *SR* histograms of these quantities, or the joint (*PR*, *SR*) 2-dimensional histogram, also referred to as the ALEX histogram, are used during different steps of the analysis. Traditionally, they are computed using a bin size of 0.02-0.04, and cover a range slightly larger than the [0, 1] interval in which ratios of quantities not corrected for background would normally fall. In this work, some of the joint (*PR*, *SR*) distributions are represented using hexagonal bins, as produced by the FRETBursts software [21, 28].

### 8.2 Weighted PR Histogram

Rejection of small bursts right after burst search is a standard analysis step. The choice of the minimum burst size *S_min_* to include in any further analysis is important and deserves some care. Increasing *S_min_* obviously reduces the number of selected bursts. This decrease is exponential (Fig. SI-12), with a characteristic size depending on the burst search type and parameters. This decrease in the final number of bursts, detrimental for sampling statistics, is however compensated by a reduction of variance of most observables, such as the PR histogram’s width (Fig. SI-13). This is due to the reduction of shot noise broadening, as more small bursts are rejected. This effect can help separating two nearby *PR* peaks, as illustrated in Fig. SI-13, where the presence of two distinct peaks in the *PR* histograms of sample 17d, measured in spot 2 of the multispot setup, requires *S_min_* ≥ 30 to be clearly distinguishable.

**Fig. SI-12:**
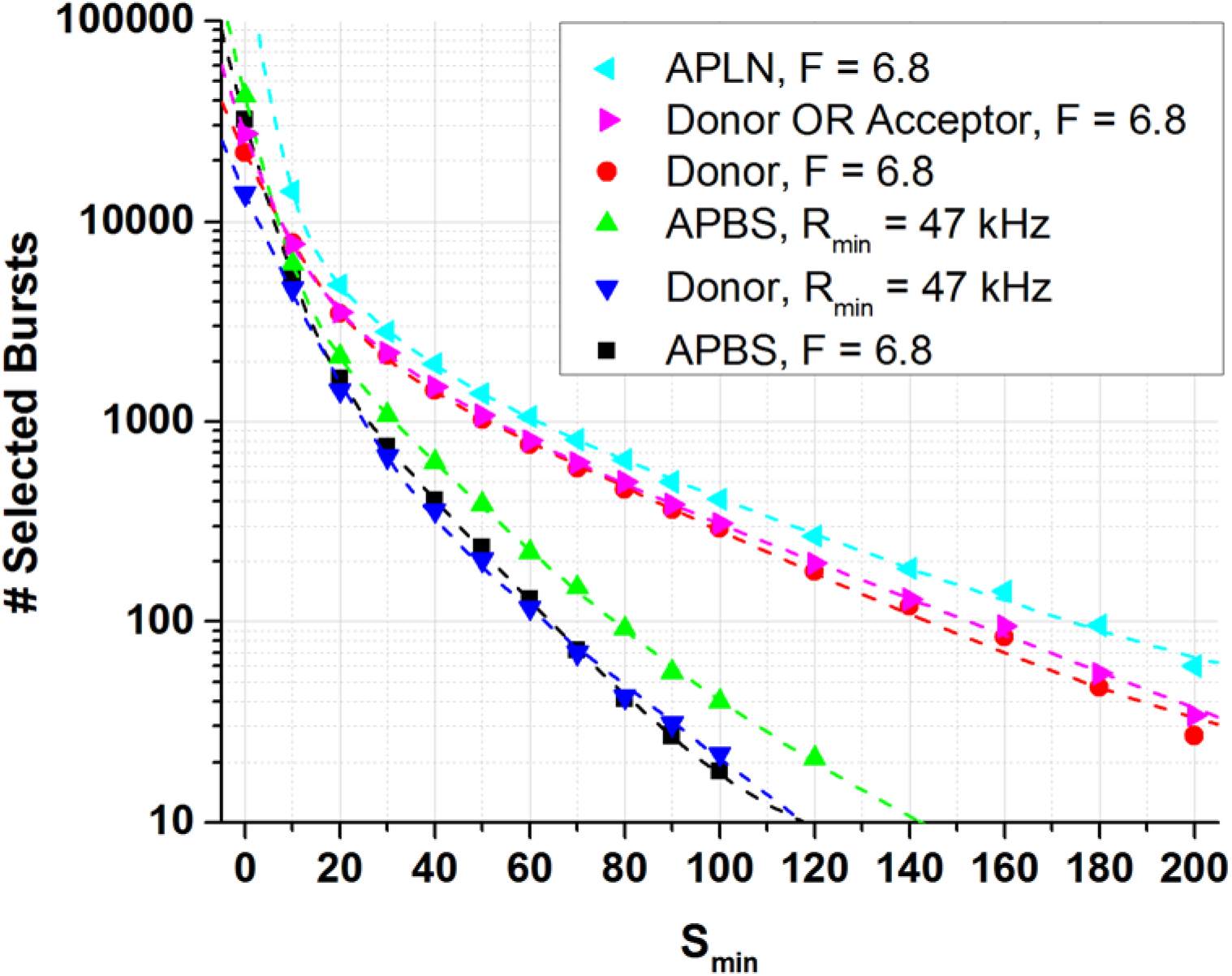
Dependence of burst number on the minimum burst size *S_min_*. Dependence of the total number of selected bursts on the minimum burst size selection parameter *S_min_*, illustrated with sample 17d, spot 2. Results for different types of burst searches are shown. Notice in particular how the standard all photon burst search (APBS) leads to the smallest number of bursts, due to the fact that the large background rate in the donor channel, used for both donor-only or FRET bursts detection, results in a higher loss of FRET bursts. APLN: all-photon burst search, using the lowest of the two donor and acceptor channel background rates; Donor (resp. Acceptor): search limited to the donor (resp. acceptor) channel photons; Donor OR Acceptor: search corresponding to the union of a Donor and an Acceptor search; *F* = 6.8: search using a count rate threshold computed as *F* times the background rate; *R_min_* = 47 kHz: fixed count rate threshold search. *m* = 5 was used for all searches. The dashed curves are multi-exponential fits to the data.

**Fig. SI-13:**
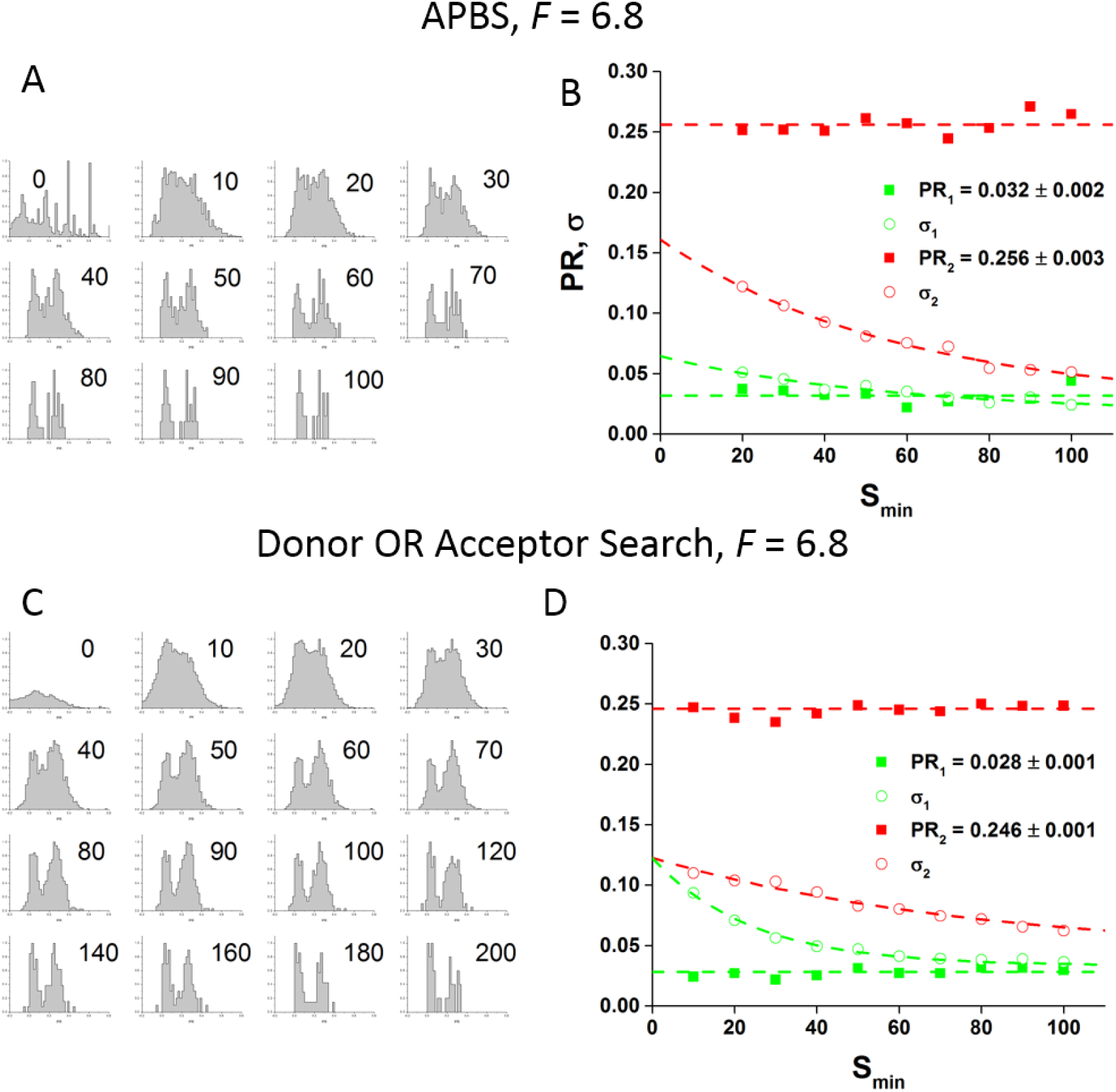
Dependence of proximity ratio statistics on the minimum burst size *S_min_*. Normalized PR histograms of bursts found by an *m* = 5, *F* = 6.8 all photons burst search (APBS) (**A**), or a donor OR acceptor burst search (OR gate of pure donor-channel search and a pure acceptor-channel search) (**C**). Smin is indicated for each plot. The horizontal axis covers [0, 1]. Both searches were followed by selecting bursts with a minimum total burst size

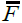

after background correction,

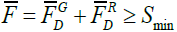

. Most PR histograms (shown on the left) exhibit two peaks which can clearly be identified for *Smin* ≥ 30 (donor-only and FRET population). Fitting the *PR* histogram by a sum of two Gaussian functions returns 6 parameters: the peak positions (*PR_i_*, *i* =1,2) and standard deviations (*σi*, also referred to informally as the “width”), as well as the fraction of each population (*f_i_*), shown on the right plots (**B** & **D**). Increasing the size threshold results in decreased PR histogram widths for both components (open symbols, dashed curves), but little changes in the peak positions (full symbols, plain curves). Dashed curves are constant fits to the peak positions or exponential fits to the peak standard deviations. Red: FRET peak, green: donor-only peak.

The previous conclusions hold whatever search type is used (minimum SBR criterion or fixed minimum count rate threshold), and whatever burst selection criterion is used, including imposing a minimum *γ*-corrected burst size or a minimum peak burst count rate. This similar behavior is due to the strong correlation between all these observables (Fig. SI-14E, F).

**Fig. SI-14:**
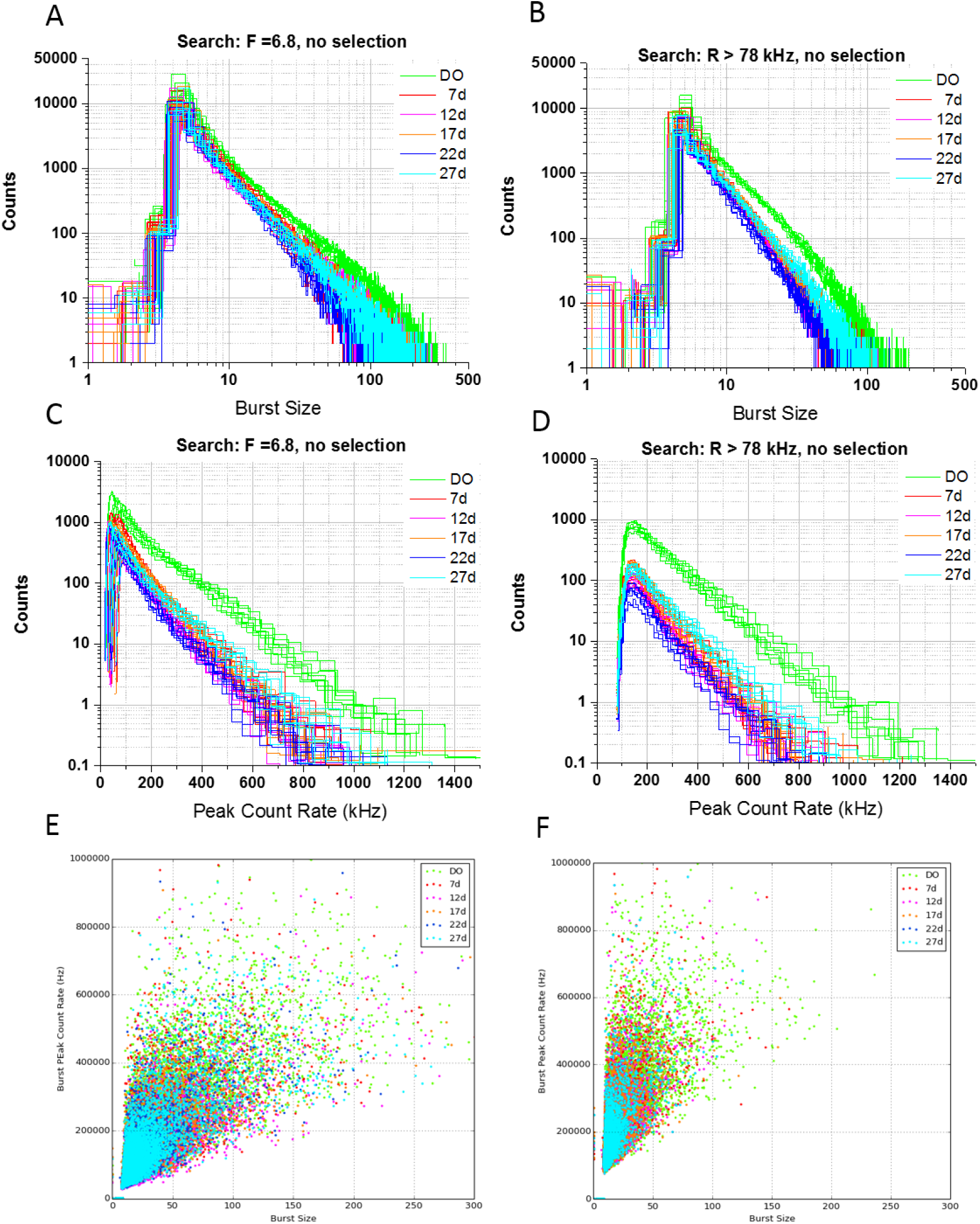
Correlation between burst statistics. (**A**) Burst size distributions for all samples, resulting from an *m*= 5,*F* = 6.8 burst search without further selection. There are 8 distributions per sample. (**C**) Corresponding total peak count rate distributions. (**B**) Burst size distributions for all samples, resulting from an *m* = 5, *R* ≥ 78 kHz burst search without further selection. There are 8 distributions per sample. (**D**) Corresponding total peak count rate distributions. (**E**) Scatterplots of total peak count rate versus burst size for spot 4. The same search (without selection) as in (A, C) was performed. (**F**) Scatterplots of total peak count rate versus burst size for spot 4. The same search (without selection) as in (B, D) was performed.

An alternative approach consists in using a “weighted” *PR* histogram, where each burst is counted not as one unit, but as a number related to its burst size. Formally, the weight factor can be expressed as a function *f* of the background-corrected burst size

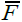

(Appendix S6 of ref. [21], doi:10.1371/journal.pone.0160716.s006):

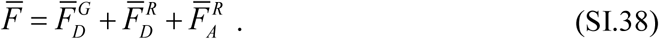

The weighted histogram can be defined formally by its bin contents *h_i_*:

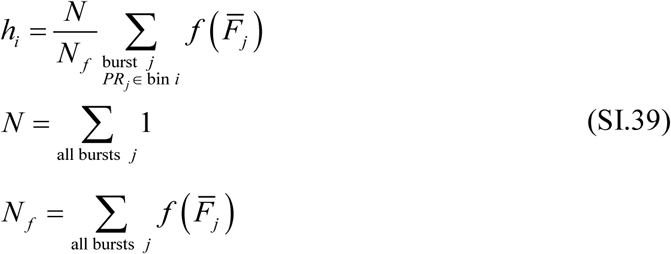

where the *i*^th^ histogram bin boundaries are those of the regular (unweighted) histogram. As discussed in the reference cited above, the choice *f(x) = x* has a simple statistical interpretation and is illustrated in Fig. SI-15, for the same sample and observation spot as used in Fig. SI-13 (17d, spot 2).

**Fig. SI-15:**
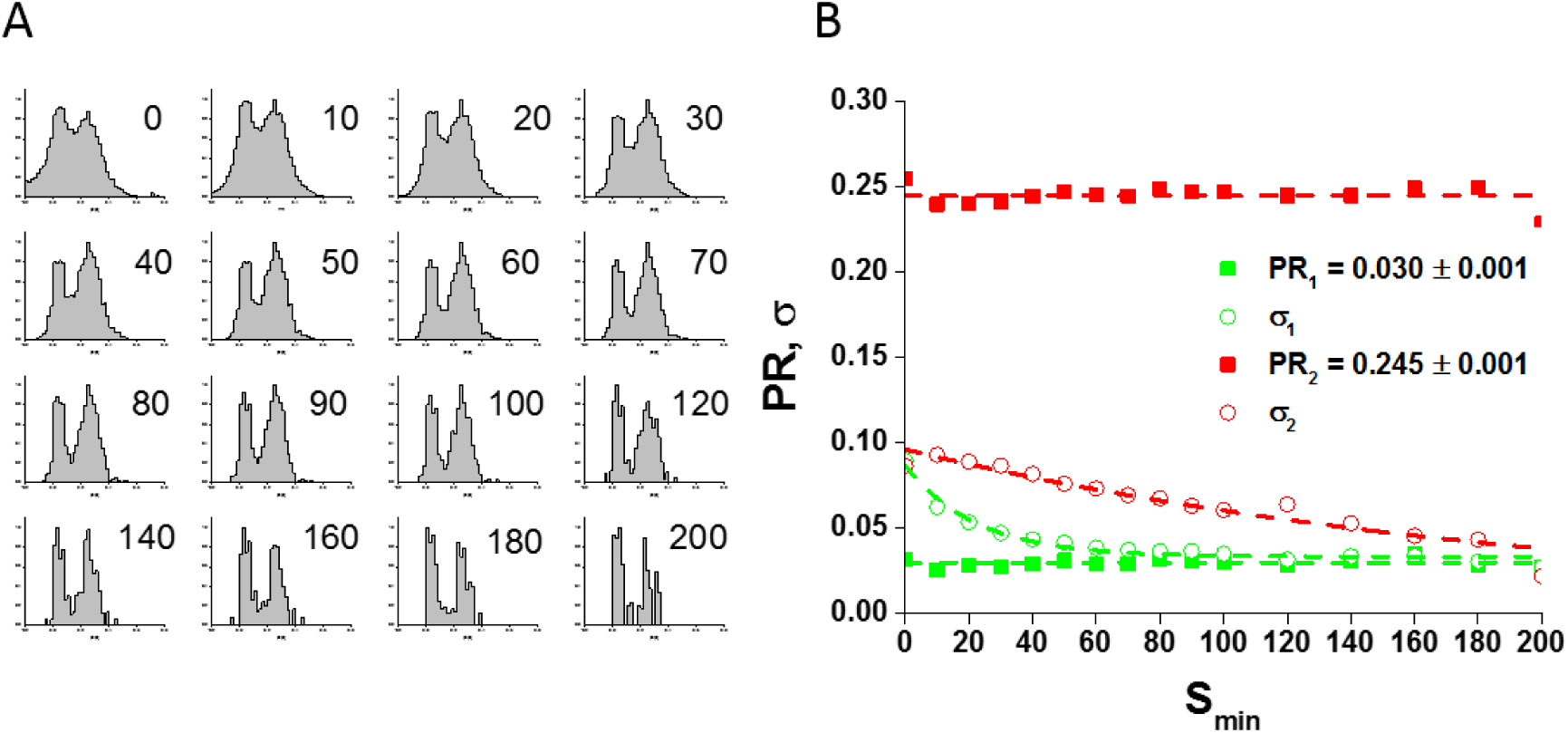
Dependence of weighted proximity ratio statistics on the minimum burst size *S_min_*(weight = 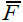). Same analysis as in Fig. SI-13, but using weighted *PR* histograms, each burst being associated with a weight equal to 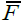, where 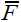 is the total burst size. Notice the identical values of the peak PR, but somewhat reduced width of the *PR* histograms. The weighted *PR* histograms (A) show a clear separation between the two populations even without burst size selection (*S_min_* = 0). (**A**) Normalized PRH, the horizontal axis covers [0, 1]. (**B**) Parameters of a 2-Gaussian Fit of the PRH.

Using a steeper function of the burst size as weight (e.g. *f(x) = x*^*2*^), increasingly suppresses the influence of the smallest bursts on the weighted *PR* histogram (data not shown). This is easily seen by looking at the effect of varying the minimum burst size used to construct these histograms (Fig. SI-15A). Loosely speaking, in this particular case, the first

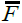

-weighted *PR* histogram (*S_min_* = 0 in Fig. SI-15A) is similar to the unweighted *PR* histogram with *S_min_* = 30 (Fig. SI-13C).

**Fig. SI-16:**
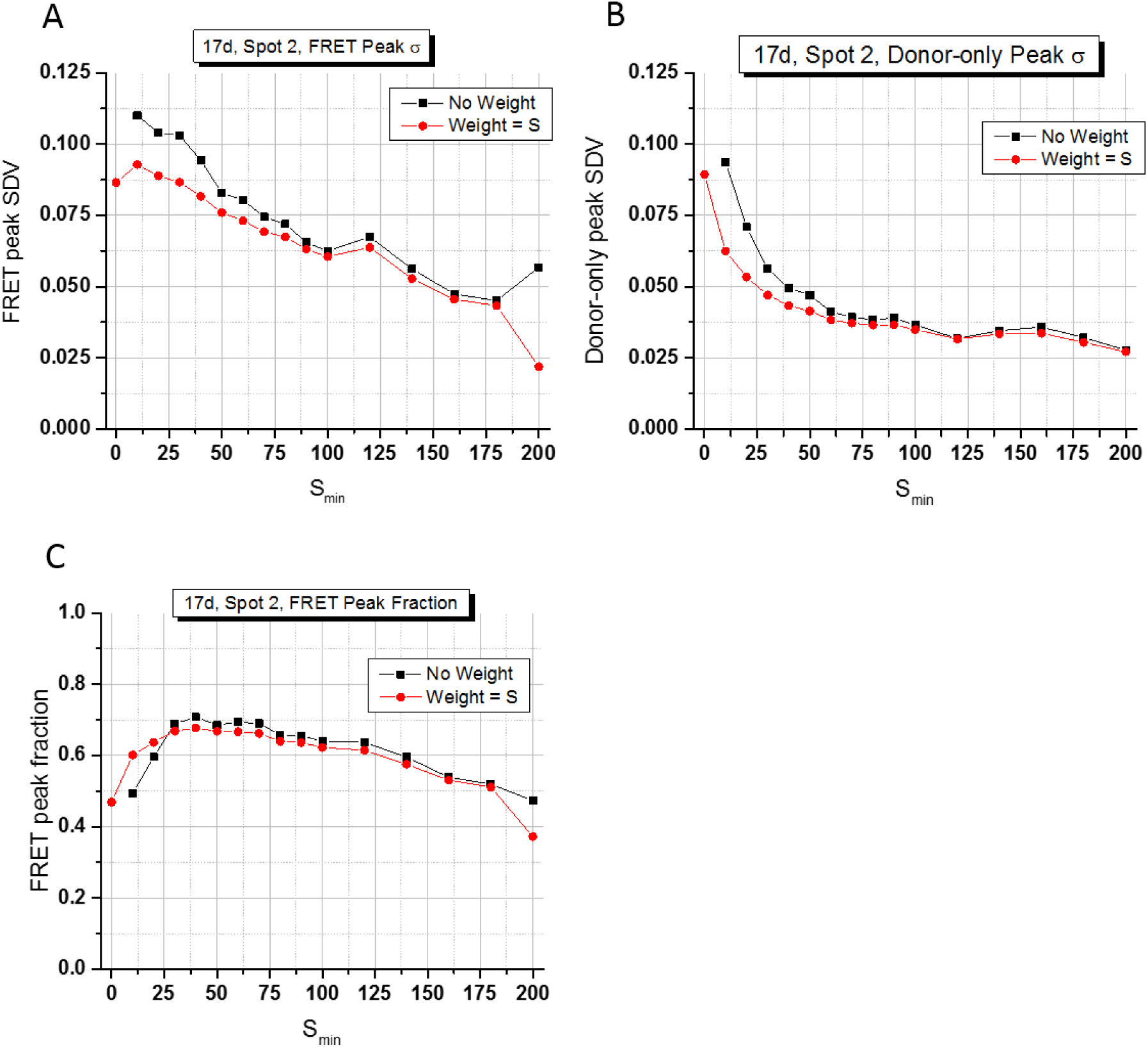
Dependence of weighted proximity ratio statistics on the minimum burst size *S_min_*. Results of the analyses presented in Fig. SI-13 & SI-15 (Donor OR Acceptor burst search). (**A**) Donor-only peak Gaussian fit parameter *σ* (width) as a function of burst size selection threshold *S_min_* for different *PR* histogram weighting schemes; (**B**) FRET peak Gaussian fit parameter *σ* (width); (**C**) FRET peak fraction. The width of both peaks decreases as *S_min_* increases, as expected from the reduction of shot noise broadening due to the rejection of small size bursts. Using a weighted *PR* histograms reduces both peak widths. The fraction of each population is dependent on *S_min_*, with a decrease of the FRET fraction as the size selection threshold increases. This effect is due to the lower brightness of FRET bursts, itself due to the lower detection efficiency in the acceptor channel. Note that this effect is present even when a weighted *PR* histogram is used, and that the measured fraction depends on the weight chosen.

To quantify this comparison, the *PR* histograms can be fitted with a sum of two Gaussians and the evolution of their parameters studied as a function of *Smin* (Fig. SI-16). Each fit returns 6 parameters: the peak positions (*PR_i_*, *i* =1,2) and standard deviations (*σ_i_*, also referred to informally as the “width”), shown on the right plots in Fig. SI-13, as well as the fraction of each population (*f_i_*) (*f_1_*+ *f_2_* = 1). Increasing the size threshold results in decreasing PR histogram widths, with no variation of the peaks positions.

## Appendix 9 Correction Factors

### 9.1 Theory

Four main corrections need to be applied on raw burst counts, in order to compute accurate FRET efficiency:

1. a stream-specific background correction,
2. a correction of the D-excitation R-detection stream for D-emission leakage in the R-channel,
3. a correction of the same stream for direct excitation of the acceptor by the D-excitation laser,
4. a correction for the difference in detection efficiency of G-and R-channels and quantum yields of the donor and acceptor species (*γ* factor correction).

We now briefly recall how the corresponding corrections and factors are computed [29]. A collection of similar derivations using notations used in FRETBursts notebooks can be found in ref. [30]. Background corrections are performed as described in Appendix 6.

#### 9.1.1 Donor Leakage Factor

The donor leakage factor *l* is obtained as the parameter resulting in an *l*-corrected D-only (DO) proximity ratio histogram centered around 0. Using the most likely value *PR_DO_* of the uncorrected proximity ratio of the DO population (see Appendix 10), the relation between *PR_DO_* and *l* is:

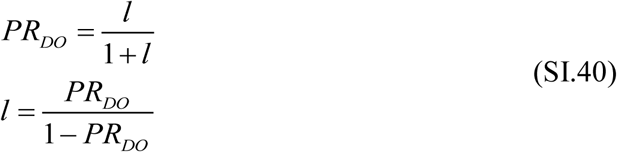

Alternatively, *l* can be defined as the parameter resulting in a D-only, leakage-corrected acceptor channel signal:

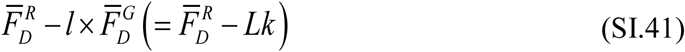

centered around 0.

#### 9.1.2 Acceptor Direct Excitation Factor

The acceptor direct excitation factor *d* can be obtained as the parameter resulting in an A-only (AO) uncorrected stoichiometry ratio centered around 0. Using the A-only, uncorrected stoichiometry ratio *SR_AO_*, the relation between *SR_AO_* and *d* is:

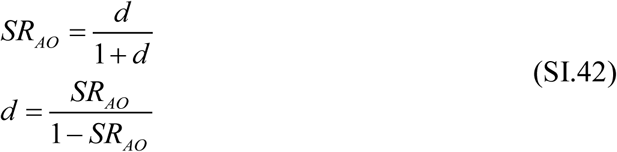

Alternatively, *d* can be defined as the parameter resulting in a direct excitation-corrected A-only, D-excitation, acceptor channel signal:

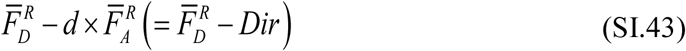

centered around 0.

We will introduce the following notation using a tilde symbol (~), for the D-excitation, A-channel signal corrected for background, leakage and acceptor direct excitation:

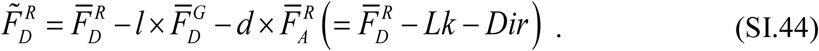

In particular, for FRET species, this quantity will be noted

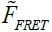

.

In order to use consistent notations, we will also introduce

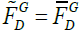

and

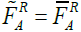

(there is no additional correction for these specific streams beside background correction).

#### 9.1.3 The *γ* Factor

The *γ* factor is obtained as described in Lee *et al.* [29], using the mean corrected proximity ratio *E_PR_* and mean corrected stoichiometry ratio *S* of the series of dsDNA FRET samples described in Section 4 of the main text.

As a reminder, the *l-* and *d-*corrected proximity ratio *E_PR_* and the *l-* and *d-*corrected stoichiometry ratio *S* are defined as follows:

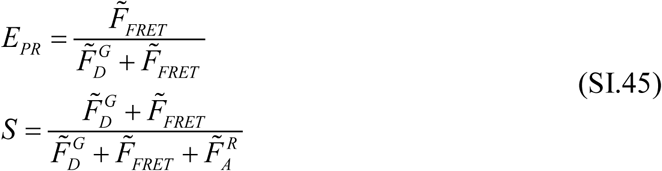

Using the most likely values of these two quantities for the FRET subpopulation of each sample, {(*E*_*PR*,*i*_, *S*_*i*_)} and fitting the following relation to this set of points:

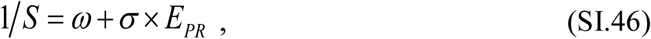

one obtains experimental parameters *ω* and *σ*, from which the *γ* factor can be computed as:

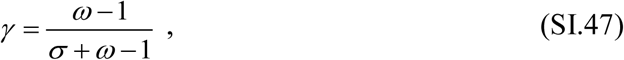

and the associated factor *β*:

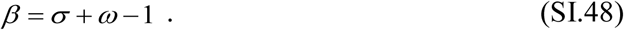

The corrected FRET efficiency *E* and stoichiometry *S_γ_* for each sample are then obtained using the following relations:

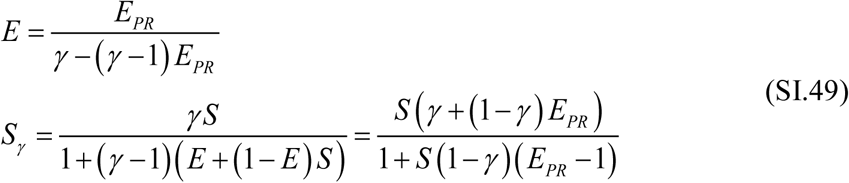

### 9.2 Single-Spot μs-ALEX Experiments

#### 9.2.1 Donor Leakage Factor

Donor leakage factors were obtained from analysis of the donor-only (D-only) population *PR* histogram of each sample, using three different approaches:

1. fit with a normal distribution;
2. analysis by kernel density estimation (KDE), yielding the peak value, which was converted into a leakage factor using Eq. (SI.40)
3. shot noise analysis (SNA) of the same *PR* was performed assuming an underlying Gaussian distribution of FRET efficiency with center and standard deviation (*E*, *σ_E_*). Eq. (SI.40) was used with *PR_DO_* = *E* to obtain the leakage factor. In this latter calculations, the best fit parameters was *σ_E_* = 0, indicating that the *PR* distribution was shot noise limited.

Table SI-3 reports the results of these analyses, which show a negligible negative bias of the standard *PR* histogram analysis results compared to shot noise analysis. The average value of the KDE results was used in subsequent analyses. Details of the analysis can be found in the μs-ALEX: Leakage coefficient section of the main Jupyter notebook [6].

**Table SI-3:** Donor-leakage factors obtained using kernel density analysis (KDE) or shot noise analysis (SNA) of the D-only *PR* histogram. Burst used for the analysis were obtained by an APBS *m* = 5, *F* = 6.8 search,

|  | 7d | 12d | 17d | 22d | 27d | All |
| --- | --- | --- | --- | --- | --- | --- |
| Normal | 0.106 | 0.106 | 0.111 | 0.096 | 0.098 | 0.103 ± 0.006 |
| KDE | 0.098 | 0.106 | 0.111 | 0.092 | 0.089 | 0.099 ± 0.009 |
| SNA | 0.115 | 0.115 | 0.120 | 0.110 | 0.104 | 0.113 ± 0.006 |
| # Bursts | 702 | 412 | 604 | 394 | 184 | 459 ± 201 |

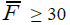

, *PR* ≤ 0.3, *SR* ≥ 0.85 selection criteria. Their number is indicated in the last row. All: mean and sample standard deviation of all samples. Results obtained using ALiX Scripts provided in ref. [7].

#### 9.2.2 Acceptor Direct Excitation Factor

A similar approach could in principle be used for the acceptor direct excitation factor provided enough acceptor-only (A-only) bursts are detected in each sample. Since this was not the case, the A-only bursts of all samples (APBS *m* = *5*, *F* = 6.8 search,

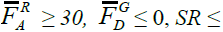

0.2 selection criteria) were accumulated and their *SR* data was analyzed by normal fitting and kernel density estimation. The peak position (*SR* = 0.044 (Normal fit), 0.041 (KDE)) was converted into a acceptor direct excitation factor using Eq. (SI.42), resulting in a acceptor direct excitation factor of 0.045. Note that the value quoted in the main text (*d* = 0.06) corresponds to a different set of search parameters (A_em_BS, *m* = 10, *F* = 7,

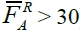

. The difference is small but points to the fact that it is difficult to obtain correction factors with an absolute precision of better than a few percents.

In general advisable to use dedicated D-only and A-only samples to estimate these correction factors reliably.

#### 9.2.3 γ Factor

The *γ* factor was obtained as described in Section 9.1.3, resulting in a value *γ* = 1.02.

### 9.3 Multispot Experiments

#### 9.3.1 Donor Leakage Factors

Leakage factors were measured as described, using 3 different methods. The results are presented in Fig. SI-17 and Table SI-4 to SI-5, and discussed in the main text.

**Fig. SI-17:**
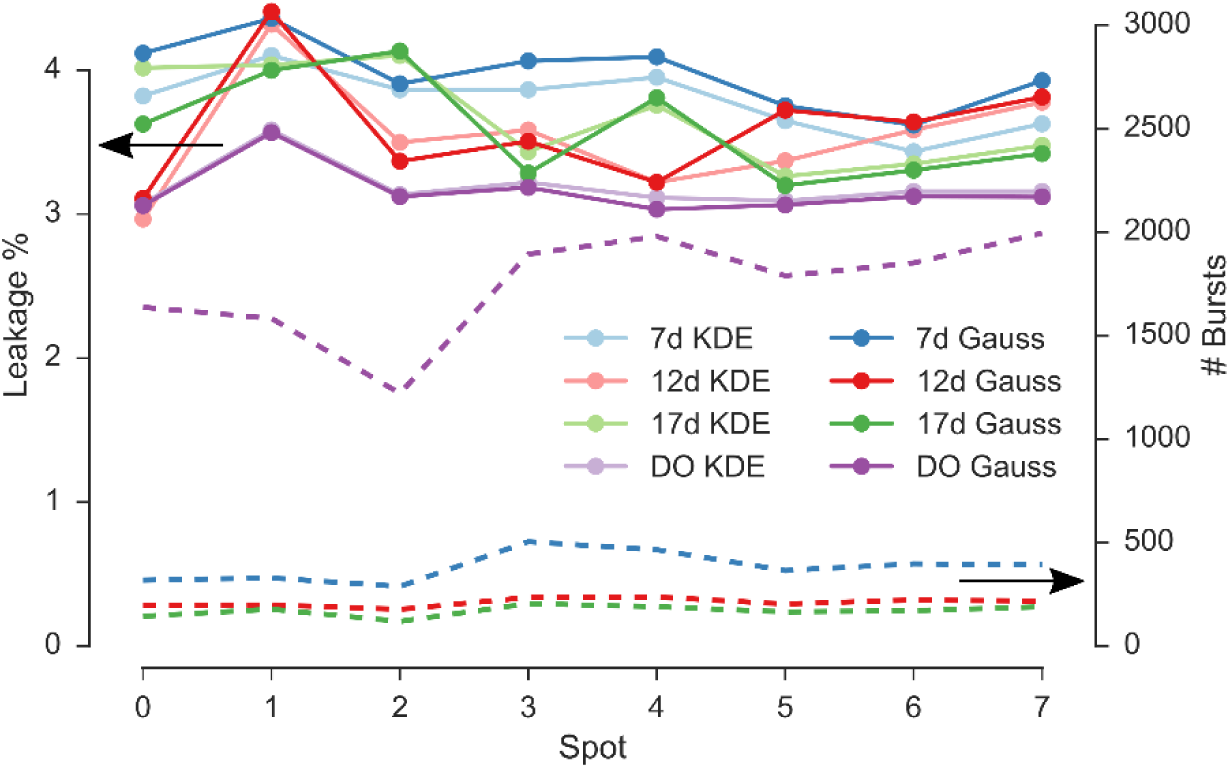
Leakage factor versus spot computed with different fitting methods in different samples. Leakage factor (*solid lines, left vertical axis*) estimated for each spot in 4 dsDNA samples (7d, 12d, 17d and DO). The leakage coefficient is computed from the DO peak position in the background-corrected *PR* distribution. The DO peak position is estimated either by kernel density estimation (*label KDE, dark color*s) or using a Gaussian fit (*label Gauss, light colors*). *Dashed lines*, *right vertical axis*: number of bursts in the DO population for the different samples (colors are identical to those used for the leakage factor). For computational details (including the numerical values used in the above figure) see section Leakage coefficient of the main Jupyter notebook [6].

**Table SI-4:** Multispot Measurement. Donor-leakage coefficients obtained for the 7d sample using Gaussian fit, kernel density analysis (KDE) or shot noise analysis (SNA, 1 replicas per burst). Bursts used for the analysis were obtained by a D_em_ burst search with *m* = 10, *F* = 6, and

| Spot | 1 | 2 | 3 | 4 | 5 | 6 | 7 | 8 | All |
| --- | --- | --- | --- | --- | --- | --- | --- | --- | --- |
| Gaussian | 0.046 | 0.137 | 0.045 | 0.044 | 0.048 | 0.054 | 0.048 | 0.044 | $0.058 \pm 0.014$ |
| KDE | 0.023 | 0.103 | 0.030 | 0.034 | 0.034 | 0.040 | 0.028 | 0.029 | $0.040 \pm 0.026$ |
| SNA | 0.054 | 0.140 | 0.044 | 0.048 | 0.046 | 0.050 | 0.058 | 0.058 | $0.062 \pm 0.032$ |
| # Bursts | 425 | 463 | 361 | 623 | 599 | 464 | 546 | 513 | $499 \pm 88$ |

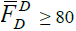

selection criterion. All: mean (sample standard deviation) of all spots. The values obtained by pooling bursts from all 8 spots (instead of performing a spot by spot analysis) were *l* = 0.051 (Gaussian fit), *l* = 0.032 (KDE) and *l* = 0.068 (SNA).

**Table SI-5:** Multispot Measurement. Donor-leakage coefficients obtained for the 12d sample using Gaussian fit, kernel density analysis (KDE) or shot noise analysis (SNA, 1 replica per burst). Bursts used for the analysis were obtained by a Dem Burst Search, *m* = 10, *F* = 6,

| Spot | 1 | 2 | 3 | 4 | 5 | 6 | 7 | 8 | All |
| --- | --- | --- | --- | --- | --- | --- | --- | --- | --- |
| Gaussian | 0.040 | 0.111 | 0.043 | 0.036 | 0.042 | 0.039 | 0.045 | 0.046 | $0.050 \pm 0.025$ |
| KDE | 0.017 | 0.075 | 0.030 | 0.031 | 0.034 | 0.022 | 0.030 | 0.037 | $0.034 \pm 0.018$ |
| SNA | 0.028 | 0.11 | 0.036 | 0.040 | 0.036 | 0.046 | 0.038 | 0.040 | $0.047 \pm 0.026$ |
| # Bursts | 233 | 235 | 206 | 270 | 287 | 247 | 261 | 247 | $248 \pm 25$ |

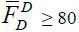

and *PR* ≤ 0.3 selection criteria. All: mean (sample standard deviation) of all spots. The values obtained by pooling bursts from all 8 spots (instead of performing a spot by spot analysis) were *l* = 0.046 (Gaussian fit), *l* = 0.033 (KDE) and *l* = 0.048 (SNA).

**Table SI-6:** Multispot Measurement. Donor-leakage coefficients obtained for the DO sample using Gaussian fit, kernel density analysis (KDE) or shot noise analysis (SNA, 1 replica per burst). Bursts used for the analysis were obtained by a D_em_ Burst Search, *m* = 10, *F* = 6,

| Spot | 1 | 2 | 3 | 4 | 5 | 6 | 7 | 8 | All |
| --- | --- | --- | --- | --- | --- | --- | --- | --- | --- |
| Gaussian | 0.030 | 0.067 | 0.035 | 0.034 | 0.034 | 0.033 | 0.019 | 0.034 | $0.036 \pm 0.014$ |
| KDE | 0.030 | 0.064 | 0.029 | 0.030 | 0.026 | 0.029 | 0.028 | 0.028 | $0.032 \pm 0.013$ |
| SNA | 0.026 | 0.050 | 0.032 | 0.032 | 0.028 | 0.028 | 0.030 | 0.030 | $0.032 \pm 0.008$ |
| # Bursts | 1618 | 1571 | 1197 | 1852 | 1952 | 1760 | 1819 | 1958 | $1716 \pm 252$ |

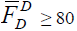

and *PR* ≤ 0.26 selection criteria. All: mean (sample standard deviation) of all spots. The values obtained by pooling bursts from all 8 spots (instead of performing a spot by spot analysis) were *l* = 0.036 (Gaussian fit), *l* = 0.029 (KDE) and *l* = 0.034 (SNA).

#### 9.3.2 Acceptor Direct Excitation Factor(s)

In the absence of acceptor excitation laser, the contribution of the direct acceptor excitation of the acceptor by the donor laser cannot be corrected using Eq. (SI.44). As shown in ref. [29], it is for instance possible to express it as a function of the donor signal coming from direct excitation by the donor laser,

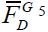

.^5^ The expression provided there (Eq. (27) in ref. [29]) was derived for a *doubly-labeled, zero-FRET sample*, for which:

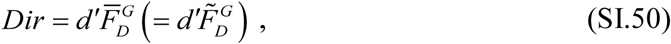

with:

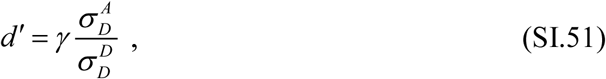

where *σ_X_^Y^* is the absorption cross-section of species Y at the wavelength of excitation laser *X*.

Formally, using the notations of ref. [29], it is possible to express *Dir* as:

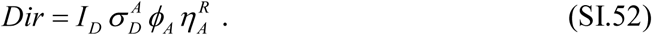

Using the definitions of the corrected donor and acceptor signals, 
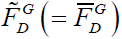
 and

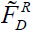

(Eq. (SI.99)), one obtains:

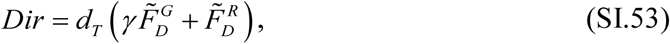

with:

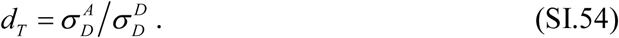

Eq. (SI.53) expresses *Dir* as a function of the *γ*-corrected burst size (Eq. (SI.101)) and is valid forany doubly-labeled molecules^6^. The direct excitation coefficient *d_T_* is an intrinsic property of the dye pair, being the ratio of D and A absorption cross-sections at the D excitation wavelength. Since Eq. (SI.53) involves the leakage and direct-excitation corrected quantity

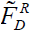

(Eq. (SI.44)), we can rewrite it in a way that isolates *Dir*, to obtain an expression involving only background-corrected burst quantities

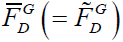

and

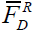

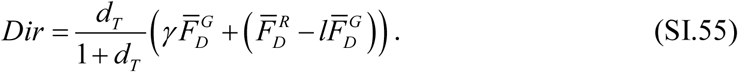

Note that for small values of parameters *d_T_*, this expression will in practice differ very little from Eq. (SI.53) where

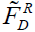

is replaced by the leakage-only corrected quantity

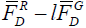

.

Another expression for *Dir* can be obtained as a function of the background-corrected burst size

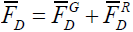

(defined in Eq. (SI.35)):

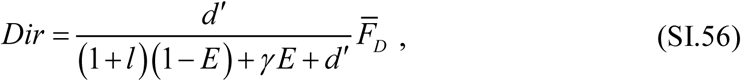

This relation is used to estimate the effect of direct acceptor excitation in Shot Noise Analysis (Appendix 11). Other relations between *Dir* and different burst quantities can be found in ref. [30].

Using factor *d_T_*, the FRET efficiency *E* can be expressed as:

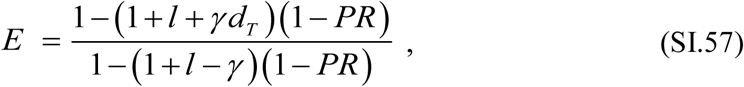

which involves correction factor *γ*, D-leakage factor *l* (both spot-specific), the A-direct excitation factor*dT* and the proximity ratio *PR* calculated without any leakage or direct excitation correction (Eq. (SI.36), *E*_*PR*_^*raw*^ in ref. [29]).

The A-direct excitation factor

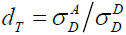

can in principle be computed from the normalized absorption spectra of the donor and acceptor dye (measured in the environment they are found in in the sample) at wavelength

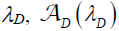

and their respective extinction coefficients *ε*_*D*_ and *ε*_*A*_:

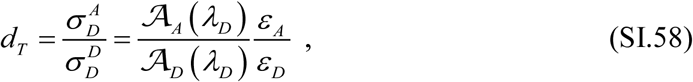

where *λ_D_* is the donor excitation laser wavelength.

Alternatively, since *d_T_* is a characteristic of the sample, not of the measurement method, it can be obtained from μs-ALEX measurements. Simple algebra shows that:

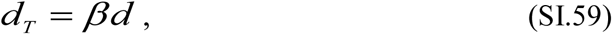

where parameter *β* is defined by Eq. (SI.104) and *d* is the direct acceptor excitation factor defined in 9.1.2.

While *d_T_* can be obtained from μs-ALEX measurements, the *γ* factor to be used in all multispot calculations is that characterizing the multispot setup, *γ_m_*. In particular, when using parameter *d’* (e.g. in Eq. (SI.56)), the following formula applies:

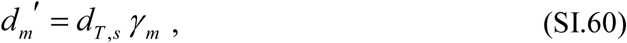

where the subscripts *s* and *m* indicate which measurement the parameter is obtained from (*s*: single-spot μs-ALEX measurement, *m*: multispot, non μs-ALEX measurements).

#### 9.3.3 γ Factor

In multispot experiments the absence of acceptor laser excitation prevents determining *γ_m_* with the procedure used for μs-ALEX data. The value has been instead determined by comparison of *PR* for one sample (chosen as calibration sample) with the *γ*-corrected FRET efficiency obtained from μs-ALEX measurement of the same sample. Formally, *γ_m_* can be computed as:

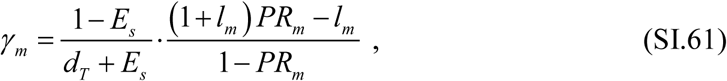

where the subscripts *s* and *m* indicate the setup with which the quantity is measured (*s*: single-spot µs-ALEX, *m*: multispot setup). Parameter *d_T_* (Eq. (SI.54)) is a characteristic of the sample and can be computed from single-spot µs-ALEX measurements. Eq. (SI.61) is obtained from Eq. (SI.57), by solving for *γ*. As discussed next, it is possible to determine the value *PR_m_* minimizing the uncertainty on *γ_m_*. In this work, sample 12d was used for that purpose.

Note that the multispot *γ* factor (*γ_m_*) could in principle be estimated on a spot-by-spot basis. This could be necessary if the *PR* peak position showed significant variation across channels, which was not the case for the measurements reported in this work. For computational details of how the *γ_m_* coefficient is computed see section PR and FRET analysis of the main Jupyter notebook.

#### 9.3.4 Error on factor *γ* estimation

Differentiating Eq. (SI.61) with respect to its two dependent variable *l* and *PR*, we obtain:

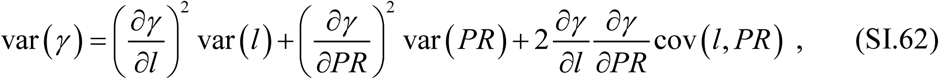

with obvious notations. Assuming independence between the estimates of *l* and *PR*:

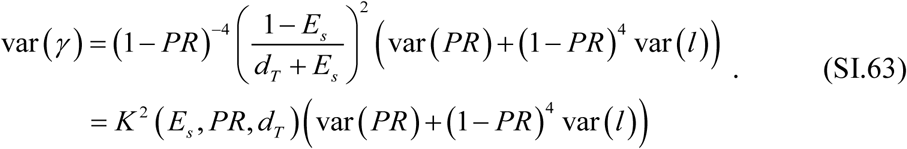

Any *PR* variance will clearly be magnified for *PR* values close to 1, while any value of *Es* small enough will amplify the right hand side of Eq. (SI.63) due to the factor (*d_T_* + *E_s_*)^-2^, where *d_T_* is in general small.

In the case where the term involving the variance of *l* is negligible compared to the term involving the variance of *PR*, and assuming that *PR* ~ *Es~ x*, the error on *γ* will minimal when Var ( *PR* ) ^1/2^ ⨯[ (*d_T_* + *x*) ( 1-*x* )]^-1^ is minimal. Assuming furthermore (which is the case here) that *d_T_* << 1, the minimum of [*x*(1-*x* )]^-1^ is obtained for *x* = ½. In other words, in order to minimize the error on the estimation of factor *γ* using Eq. (SI.61), a sample with FRET efficiency close to 1/2 is preferable, assuming that var(*PR*) is similar for all samples. Using the above approximations, the error on this estimate is given by:

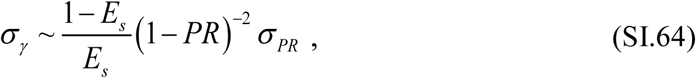

where *σ_X_* designates the standard deviation of *X*.

In general, when var(*PR*) and var(*l*) are similar for all samples, the sample minimizing the uncertainty on factor *γ* is the one minimizing *K* in Eq. (SI.63):

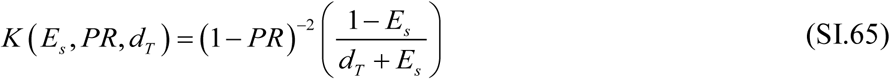

#### 9.3.5 Proximity Ratios

Proximity ratios estimated by single peak Gaussian fit, KDE analysis and SNA analysis are reported below for each sample. Details on SNA analysis can be found in Appendix 11. The values reported here are those obtained using the ALiX scripts provided in [7]. For similar values computed with the FRETBursts software see section PR and FRET analysis of the main Jupyter notebook [6].

**Table SI-7:** Multispot Measurement. PR values obtained for the 7d sample using Gaussian fit, kernel density analysis (KDE) or shot noise analysis (SNA, mean value, 1 replica per burst). Bursts used for the analysis were obtained by an APBS *m* = 10, *F* = 7 search,

| Spot | 1 | 2 | 3 | 4 | 5 | 6 | 7 | 8 | All |
| --- | --- | --- | --- | --- | --- | --- | --- | --- | --- |
| Gaussian | 0.87 | 0.86 | 0.83 | 0.84 | 0.83 | 0.85 | 0.84 | 0.85 | $0.85 \pm 0.01$ |
| KDE | 0.88 | 0.87 | 0.84 | 0.85 | 0.84 | 0.86 | 0.84 | 0.85 | $0.85 \pm 0.01$ |
| SNA | 0.91 | 0.90 | 0.86 | 0.88 | 0.88 | 0.90 | 0.88 | 0.88 | $0.82 \pm 0.01$ |
| # Bursts | 548 | 625 | 515 | 793 | 768 | 253 | 1040 | 1269 | $726 \pm 318$ |

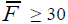
 and PR ≥ 0.5 selection criteria. All: mean ± sample standard deviation of all spots. The values obtained by pooling bursts from all 8 spots (instead of performing a spot by spot analysis) were *PR*= 0.85 (Gaussian fit), *PR* = 0.85 (KDE) and *PR* = 0.82 ± 0.05 (SNA: mean ± standard deviation). Data computed using ALiX Scripts 6-8.

**Table SI-8:**
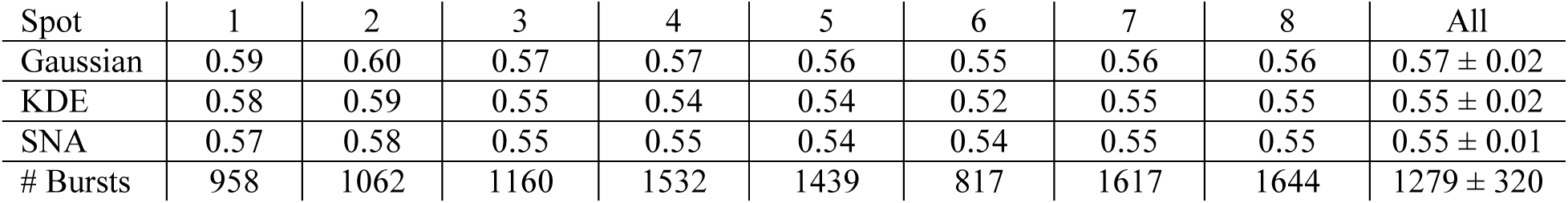
Multispot Measurement. PR values obtained for the 12d sample using Gaussian fit, kernel density analysis (KDE) or shot noise analysis (SNA, mean value, 1 replica per burst). Bursts used for the analysis were obtained by an APBS *m* = 10, *F* = 7 search,

| Spot | 1 | 2 | 3 | 4 | 5 | 6 | 7 | 8 | All |
| --- | --- | --- | --- | --- | --- | --- | --- | --- | --- |
| Gaussian | 0.59 | 0.60 | 0.57 | 0.57 | 0.56 | 0.55 | 0.56 | 0.56 | $0.57 \pm 0.02$ |
| KDE | 0.58 | 0.59 | 0.55 | 0.54 | 0.54 | 0.52 | 0.55 | 0.55 | $0.55 \pm 0.02$ |
| SNA | 0.57 | 0.58 | 0.55 | 0.55 | 0.54 | 0.54 | 0.55 | 0.55 | $0.55 \pm 0.01$ |
| # Bursts | 958 | 1062 | 1160 | 1532 | 1439 | 817 | 1617 | 1644 | $1279 \pm 320$ |

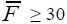

and *PR* ≥ 0.24 selection criteria. All: mean ± sample standard deviation of all spots. The values obtained by pooling bursts from all 8 spots (instead of performing a spot by spot analysis) were *PR*=0.57 (Gaussian fit), *PR* = 0.56 (KDE) and *PR* = 0.56 ± 0.05 (SNA:: mean ± standard deviation). Data computed using ALiX Scripts 9-11.

**Table SI-9:** Multispot Measurement. PR values obtained for the 17d sample using Gaussian fit, kernel density analysis (KDE) or shot noise analysis (SNA, mean value, 1 replica per burst). Bursts used for the analysis were obtained by an APBS *m* = 10, *F* = 7 search,

| Spot | 1 | 2 | 3 | 4 | 5 | 6 | 7 | 8 | All |
| --- | --- | --- | --- | --- | --- | --- | --- | --- | --- |
| Gaussian | 0.30 | 0.31 | 0.28 | 0.28 | 0.29 | 0.29 | 0.28 | 0.29 | $0.29 \pm 0.01$ |
| KDE | 0.29 | 0.30 | 0.26 | 0.27 | 0.28 | 0.27 | 0.26 | 0.26 | $0.27 \pm 0.02$ |
| SNA | 0.29 | 0.29 | 0.27 | 0.28 | 0.29 | 0.28 | 0.27 | 0.28 | $0.28 \pm 0.01$ |
| # Bursts | 505 | 505 | 496 | 663 | 676 | 391 | 757 | 811 | $600 \pm 147$ |

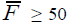

and *PR* ≥ 0.12 selection criteria. The larger burst size threshold was used in order to better separate the D-only population from the FRET population. All: mean ± sample standard deviation of all spots. The values obtained by pooling bursts from all 8 spots (instead of performing a spot by spot analysis) were *PR* = 0.28 (Gaussian fit), *PR* = 0.27 (KDE) and *PR* = 0.29 ± 0.03 (SNA). Data computed using ALiX Scripts 12-14.

**Table SI-10:** Multispot Measurement. PR values obtained for the 22d sample using Gaussian fit, kernel density analysis (KDE) or shot noise analysis (SNA, mean value, 1 replicas per burst). Bursts used for the analysis were obtained by an APBS *m* = 10, *F* = 7 search,

| Spot | 1 | 2 | 3 | 4 | 5 | 6 | 7 | 8 | All |
| --- | --- | --- | --- | --- | --- | --- | --- | --- | --- |
| Gaussian | 0.12 | 0.15 | 0.10 | 0.11 | 0.10 | 0.13 | 0.11 | 0.11 | $0.12 \pm 0.02$ |
| KDE | 0.11 | 0.14 | 0.08 | 0.08 | 0.07 | 0.11 | 0.08 | 0.07 | $0.09 \pm 0.02$ |
| SNA | 0.11 | 0.14 | 0.09 | 0.10 | 0.09 | 0.11 | 0.09 | 0.09 | $0.10 \pm 0.02$ |
| # Bursts | 876 | 825 | 946 | 1310 | 1092 | 744 | 1243 | 1386 | $1053 \pm 240$ |

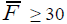

selection criterion. All: mean ± sample standard deviation of all spots. The values obtained by pooling bursts from all 8 spots (instead of performing a spot by spot analysis) were *PR* = 0.11 (Gaussian fit), *PR* = 0.09 (KDE) and *PR* = 0.10 ± 0.04 (SNA). Data computed using ALiX Scripts 15-17.

**Table SI-11:** Multispot Measurement. PR values obtained for the 27d sample using Gaussian fit, kernel density analysis (KDE) or shot noise analysis (SNA, mean value, 1 replicas per burst). Bursts used for the analysis were obtained by an APBS *m* = 10, *F* = 7 search,

| Spot | 1 | 2 | 3 | 4 | 5 | 6 | 7 | 8 | All |
| --- | --- | --- | --- | --- | --- | --- | --- | --- | --- |
| Gaussian | 0.10 | 0.13 | 0.09 | 0.09 | 0.09 | 0.10 | 0.09 | 0.09 | $0.10 \pm 0.01$ |
| KDE | 0.08 | 0.10 | 0.07 | 0.07 | 0.07 | 0.08 | 0.07 | 0.08 | $0.08 \pm 0.01$ |
| SNA | 0.09 | 0.12 | 0.08 | 0.08 | 0.08 | 0.08 | 0.08 | 0.08 | $0.08 \pm 0.01$ |
| # Bursts | 1459 | 1370 | 1721 | 2063 | 2111 | 1498 | 2147 | 2286 | $1832 \pm 361$ |

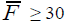

selection criterion. All: mean ± sample standard deviation of all spots. The values obtained by pooling bursts from all 8 spots (instead of performing a spot by spot analysis) were *PR* = 0.09 (Gaussian fit), *PR* = 0.07 (KDE) and *PR* = 0.08 ± 0.02 (SNA). Data computed using ALiX Scripts 18-20.

## Appendix 10 Determination of Proximity Ratio and FRET Efficiency

A common problem in single-molecule burst analysis consists in determining the “characteristic” value of an observable for a selected burst sub-population. An example is provided by the proximity ratio (or the FRET efficiency) of a doubly-labeled population.

The common way of approaching this question has long been to compute a population-specific histogram of the observable and fit this histogram with an ad-hoc function (oftentimes a normal distribution) and report the function’s peak position.

The first problem with this approach is that it is not always obvious which histogram bin size to choose for an optimal representation [23]. Secondly, a normal (Gaussian) distribution is not necessarily the best model function to fit the resulting histogram. While a Gaussian distribution might provide a reasonable fit for FRET efficiencies in the range 0.2 ≤ *E* ≤ 0.8, asymmetric distributions are often encountered for small or large values of *E*. Finally, for asymmetric functions, it might not be obvious which quantity (mean, mode or median) to report in order to characterize the distribution.

Moreover, the distribution of an observable is affected by shot noise, whose magnitude depends on burst size. Small bursts generally result in asymmetric distribution for ratiometric observables (e.g. the proximity ratio), while large bursts may result in large variance for some observables (e.g. the ratio of donor and acceptor signal in a FRET sample). Since an experiment is characterized by a distribution of burst sizes, the overall effect on the distribution of the observable of interest can be complex, rendering the choice of a model function problematic. In particular, even with a knowledge of the effect of various burst quantity distributions (size, duration, etc.), it is rarely feasible to obtain a satisfactory “fit” of an observed histogram with a few parameters only.

For all these reasons, it would seem preferable to use a model-free approach.

The simplest model-free approach consists in reporting simple statistical measures of the observable such as the mean, median, mode, standard deviation, etc. However, these statistical measures can be biased or have a large variance. While these characteristics can be determined theoretically for model functions, incorporating shot noise effects in the equation rapidly makes this determination a complex problem.

Another model-free and almost parameter-free approach is provided by kernel density estimation (KDE) [31]. In KDE, a normalized function is used as kernel (typically a normal distribution) and scaled by a bandwidth parameter to represent each observable data point in the sample. The sum of these “replicas” of the kernel density is then taken as an approximation of the underlying probability distribution function (PDF) of the observable. The result can be described as a smoothed version of a standard histogram representation of the observable with a particular bandwidth. Because it is a continuous function, the median of the distribution can be computed unambiguously. However, because the bandwidth parameter affects the shape of the distribution, there is no guarantee that a mode value exists (there could be several local maxima in the KDE distribution) or, if it exists, that it is not biased.

All these methods (fit by *ad hoc* model functions, calculation of statistical measures, KDE analysis) ignore that, in practice, the observable distribution is due to an underlying distribution of molecular properties, convolved with an “instrument response function” (IRF) involving the whole optical setup and its detectors, the main feature of which is that observable values are obtained from small count numbers and therefore are affected by shot noise. The way to account for shot noise effects on burst observable is well understood, and involves taking into account the joint distribution of burst size and burst duration [23, 32]. There is some debate as to which additional parameters (molecular, setup characteristics) need to be added to the convolution in order to account for the observed distributions. Appendix 11 discusses these issues.

## Appendix 11 Shot Noise Analysis

Shot noise analysis (also sometimes referred to as probability distribution analysis or PDA in the literature) [23, 32], in its simplest form used in this work, consists in comparing the measured proximity ratio histogram (PRH) to a predicted one based on the knowledge of burst counts, background rates and correction factors, and some adjustable model parameters describing the sample’s properties. Ideally, a single parameter, the FRET efficiency, should be needed to account for the PRH of a single, static population of doubly-labeled molecules. In practice, multiple sources of imperfection render this assumption too simplistic. Among others, the presence of D-only molecules, bleaching or blinking of the donor or acceptor molecules, dependence of the excitation and detection efficiencies on each molecule’s trajectory, and finally, possible distance distributions or dynamic fluctuations, are some of the possible effects to take in to account in practical situations.

Here, we extend the model described in ref. [23] to:

- include the acceptor-excitation signal provided by µs-ALEX alternation (for correction of the FRET signal for direct excitation of the acceptor by the donor excitation laser)
- take into account the effect of a *γ* factor significantly different from 1 in the multispot measurements
- introduce a variant of the Gaussian distribution of distances to account for additional PRH broadening.

### 11.1.1 Using the acceptor excitation information in µs-ALEX measurements in SNA

In the Monte-Carlo approach used in ref. [23], each burst was characterized by its size *S* and duration *τ*, where *S* designates the total signal detected during donor excitation. Taking into account that there is a donor excitation period as well as an acceptor excitation period in µs-ALEX measurements, each burst can be characterized by its total signal during each excitation period, *F_D_* and *F_A_*, with:

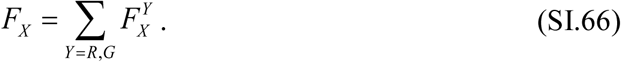

With these notations, and using the definitions introduced in the main text, the algorithm used to generate a shot noise limited PRH based on the selected bursts is as follows:

i. Choose an oversampling factor *N* (integer number) and a FRET efficiency *ε*,
ii. For each burst, compute the

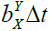

’s, where *Δt* is the burst duration.
iii. Draw one random number

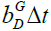

, where Δt is the burst duration.
iv. Draw one random number *a* from the Poisson distribution of mean

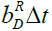

,
v. Draw one random number *r* from the Poisson distribution of mean

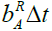

,
vi. Draw one random number *g* from the Poisson distribution of mean

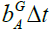

,
vii. Draw one random number *Dir* from the Poisson distribution of mean *d*(*FA– r-g*) [direct acceptor excitation],
viii. Draw one random number

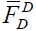

from the binomial distribution of size *FD– d – a – Dir* and probability

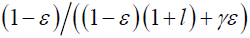

[donor signal not due to direct excitation],
ix. Define

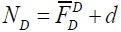
 and 
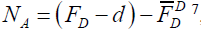

^7^
x. Add the value

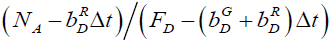

to the PRH^8^,
xi. Repeat (iii)-(ix) *N* times,
xii. Repeat (ii)-(ix) for each burst,
xiii. Divide the final PRH by *N*,
xiv. Compare the result to the measured PRH (*e.g.* using the mean square error) and improve *ε*.

When the FRET efficiency is modeled not by a single parameter, but several (as is the case when it is assumed that there is an underlying distribution of distances accounting for the observed PRH), the search for the optimal set of parameters was performed by a systematic exploration of the parameter space depending on the model chosen, as described in later sub-sections. The reason for this brute force approach is twofold:

- The range of parameters explored in this manner is well-circumscribed.
- The mean square error (MSE) map resulting from this approach is useful to gauge the quality of the minimum found.

Note that for PRH with poor statistics (small number of bursts, say, <100), choosing a large oversampling factor *N* is counterproductive, because the final PRH obtained in step (xii) is then much smoother than the actual PRH. On the other hand, when the PRH contains a large number of bursts (say, > 10,000), using a small *N* is recommended, due to the linear increase of computation time with *N*. A value *N* = 1 to 10 appears to be a good compromise.

### 11.1.2 Algorithm for shot noise analysis in multispot measurements

In the absence of an acceptor excitation laser, and of a *γ* factor markedly different from 1, we used a modified algorithm as follows, based on the only available burst size *F_D_*:

i. Choose an oversampling factor *N* (integer number) and a FRET efficiency *ε*,
ii. For each burst, compute the

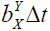

’s, where *Δt* is the burst duration.
iii. Draw one random number *d* from the Poisson distribution of mean

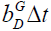

,
iv. Draw one random number *a* from the Poisson distribution of mean

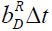

,
v. Draw one random number *Dir* from the Poisson distribution of mean

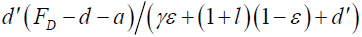

[direct acceptor excitation, Eq. (SI.56)],
vi. Draw one random number

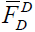

from the binomial distribution of size *FD– d – a – Dir* and probability

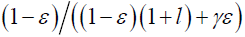

[donor signal not due to direct excitation]
vii. Define

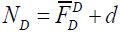
 and 
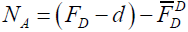
,
viii. Add the value

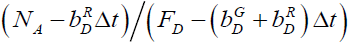

to the PRH,
ix. Repeat (iii)-(ix) *N* times,
x. Repeat (ii)-(x) for each burst,
xi. Divide the final PRH by *N*,
xii. Compare the result to the measured PRH (*e.g.* using the mean square error) and improve *ε*.

### 11.1.3 Normal FRET efficiency distribution model

While broadening of the PRH beyond shot noise can be attributed to many underlying phenomena, the simplest way to quantify this broadening is by interpreting it as due to an underlying distribution of FRET efficiencies. This is particularly useful for the analysis of the donor-only population, where the notion of a distance distribution is meaningless in the absence of an acceptor fluorophore. The simplest conceivable model is a normal distribution, centered on a value *E*_0_ and characterized by a standard deviation *σ*_*E*_. Since FRET efficiency values smaller than 0 or larger than 1 are meaningless, the actual distribution is truncated:

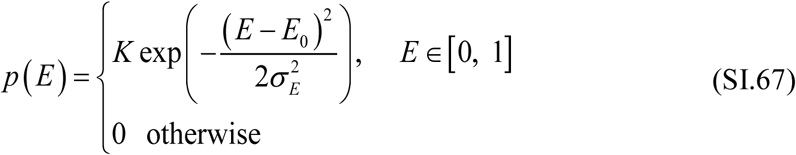

where *K* is a normalization constant ensuring that the integral of *p(E)* over [0, 1] equals 1. With this definition, parameter *σ*_*E*_ is not strictly the standard deviation of the distribution (in particular if *E*_0_ ~ 0) but can still be interpreted conveniently.

This model was only used to analyze the donor-only population and verify that the observed PRH was compatible with *E*_0_ ~ 0,*σ*_*E*_ ~ 0. For other samples (or subpopulations), the distance model discussed in the next section was used.

### 11.1.4 Beta distribution of FRET efficiencies

A slightly more complex model of FRET efficiency distribution is provided by the family of beta distributions of parameters (*a, b*):

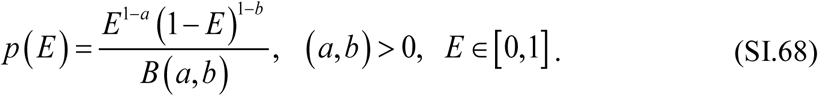

Parameters (*a, b*) can be expressed in terms of the mean *E_0_* and standard deviation *σ_E_* of the distribution:

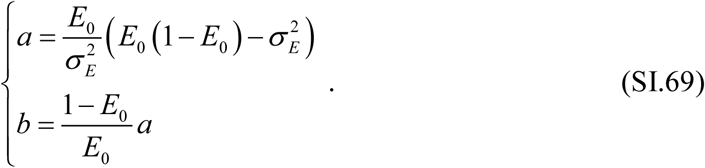

It turns out that the beta distribution *p*(*E*) with parameters (*E_0_*, *σ_E_*) differs very little from the third (geometric) model discussed in the next section.

### 11.1.5 Fuzzy dumbbell model of distance distribution

In ref. [23], a simple model of normal distribution of distances was used to quantify the small departure from the shot noise limited PRH. Calling

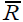

the mean distance between the two fluorophores and *σ*_*R*_ the standard deviation of the distribution, the PDF was defined as:

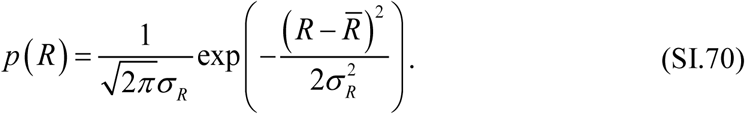

While this distribution is convenient, it is not based on any physical model, which should reflect the fact that two fluorophores attached to two flexible linkers are involved in the calculation of *R*. In this respect, a slightly more natural model consists in assuming that each dye’s location is characterized by an average position

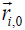

(*i* = 1 or 2) and standard deviation *σ*_*i*_. Noting

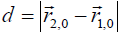

the distance between the two average dye positions, a simple calculation yields the interdye distance PDF [33]:

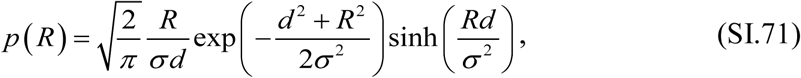

where parameter σ is defined by:

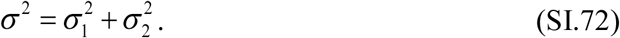

Because this model described two normally distributed objects separated by a fixed distance between their average locations, we refer to this model as the “fuzzy dumbbell” model.

This model has a few useful properties. First, it has effectively only 2 free parameters, *σ* and *d*, even though the underlying picture involves 3 parameters (*d*, *σ*_1_ and *σ*_2_). This means that very different situations could result in identical outcomes, as far as the PRH is concerned. In other words, once optimal model parameters *σ* and *d* have been obtained, there remain some flexibility (Eq. (SI.72)) to describe the system, without the need for additional model parameters. Second, contrary to the normal distribution of distance, the fuzzy dumbbell model (Eq. (SI.71)) can yield asymmetric distance distributions, which might be a desirable feature in some situations. Note that both models result in asymmetric FRET efficiency distributions for low and high FRET efficiencies, as is experimentally observed:

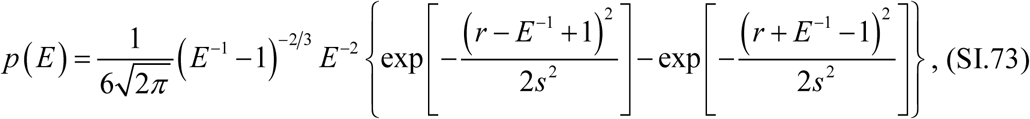

where *r* = *d/R_0_* and *s* = *σ/R_0_* are two reduced parameters expressed in terms of *d*, *σ* and the Förster radius *R_0_* (note that the Förster radius is not needed, unless the fit parameters *r* and *s* need to be converted to real units).

For all models, the search for the best fit parameter set was performed by an exhaustive grid search between min and max values for each parameters. For each set, a value *ε* from the corresponding *p(E)* PDF was used in each step (i) of the procedure described above.

Results of SNA for the single-spot measurements are represented on Fig. SI-18. Results of the 3 types of analysis (Gaussian fit, KDE and SNA) for the multispot measurements are provided in Table SI-7 to SI-11.

**Fig. SI-18:**
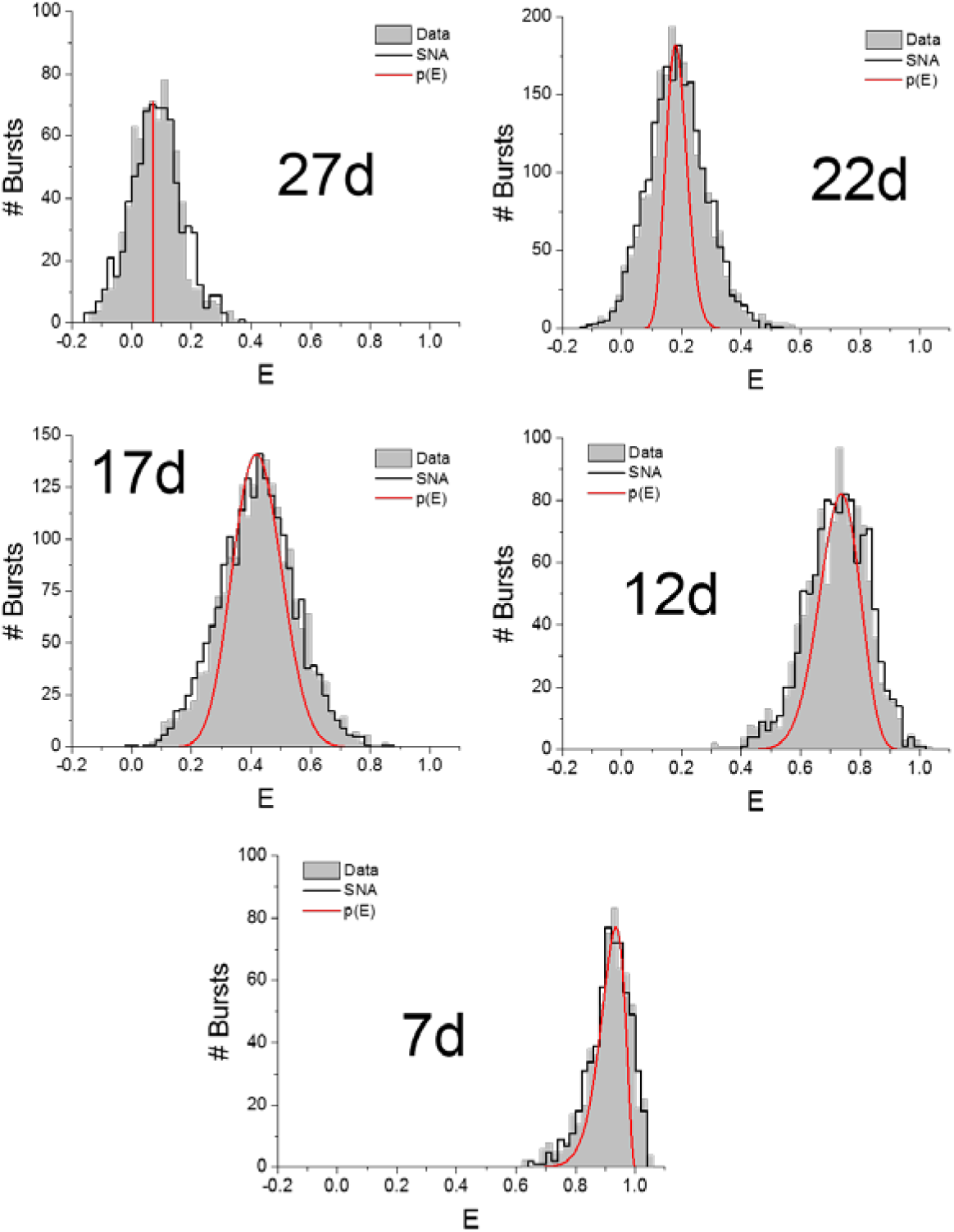
SNA results for the μs-ALEX measurements. For each sample, the corrected FRET histogram is represented in dark gray, the fitted histogram is represented in black. Superimposed to these two histograms is the E distribution used to account for the histogram (Eq. (SI.73)). The corresponding parameters (*E_0_*, *σ_E_*) of the beta distribution model are:

| Sample | 7d | 12d | 17d | 22d | 27d |
| --- | --- | --- | --- | --- | --- |
| $E_0$ | 0.913 | 0.725 | 0.423 | 0.185 | 0.080 |
| $\sigma_E$ | 0.045 | 0.068 | 0.080 | 0.035 | 0 |

The value *σ_E_* = 0 obtained for the 27d sample simply means that there is practically very little difference between this choice of parameter and larger values (such as those obtained for other samples).

## Appendix 12 FCS Analysis

This Appendix first presents details on the FCS analysis performed on both single-spot and multispot data (using ALiX), followed by results mentioned in the main text.

There are two main differences between the single-spot and multispot data as far as FCS analysis is concerned:

1. The single-spot data was obtained in the presence of laser alternation, which needs to be handled specifically in order to compare ACF and CCF curves to standard models.
2. The multispot data suffered in some cases from much larger afterpulsing, which made some of the standard ACF corrections inadequate.

This explains the presence of two distinct sections for the FCS analysis part.

### 12.1 Single-Spot µs-ALEX FCS

Fluorescence correlation spectroscopy (FCS) is a powerful tool to analyze molecular diffusion coefficients, brightness and stoichiometry and in some cases, short time scale dynamics [34]. In particular, the typical diffusion time through the observation volume, *τ_D_*, can be extracted from a fit of the autocorrelation function (ACF) to the appropriate model function. The practical and quantitative implementation of this approach is usually perilous, due to a number of potential artifacts and necessary approximations [35, 36]. Since the molecules studied here are doubly-labeled, it is also possible to compute the cross-correlation function (CCF) of the signals detected in each channel corresponding to a spot.

The ACFs and CCF of the donor and acceptor channel signals upon donor excitation (photon streams D_ex_D_em_ and D_ex_A_em_) together with the ACF of the A_ex_A_em_ stream (acceptor excitation, acceptor channel detection) were calculated on a multitau time scale using the recorded arrival times using published algorithms [37]. Because of µs laser alternation, the raw ACFs, *ACF_ALEX_*(*τ*), exhibit a periodic modulation, which can be cancelled out by a simple renormalization by the alternation period histograms’ ACFs (*ACF_Period_*(*τ*)):

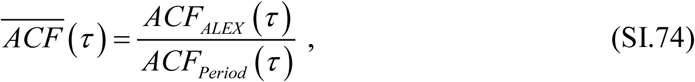

where the alternation period histograms obtained in Appendix Appendix 5 are extended over the whole duration of the experiment using their periodicity property.

As usual for SPADs, the short time scale part of the ACFs is contaminated with afterpulsing and can be corrected using the simple approach described in ref. [38], provided the time scales of interest are sufficiently well separated:

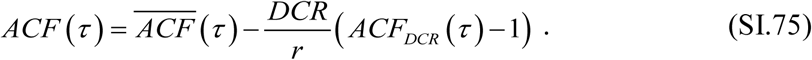

In this expression, *DCR* is the dark count rate of the detector (measured in a separate experiment), *ACF_DCR_*(*τ*) its autocorrelation function and *r* is the actual count rate of the photon stream (computed over the period defining the photon stream under consideration, not the whole alternation period).

ACFs obtained after these corrections were fitted by a simple model of 2-dimensional diffusion through a Gaussian observation volume (beam waist: *ω*), with an additional blinking component (fraction *λ*, time scale *τ_bl_*):

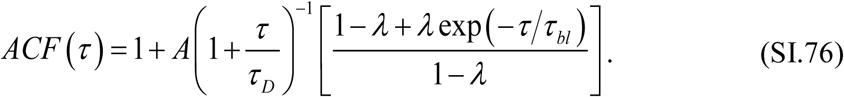

In this expression, the ACF amplitude *A* is related to:

i. the mean occupancy *N* of the observation volume,
ii. the signal-to-uncorrelated-background ratio for the photon stream under consideration,
iii. the fraction as well as brightness of different populations,

in a complex manner requiring careful calibrations to be properly computed [39]. Since we are not interested in this information, no attempt was made to compute *N*.

The diffusion constant, *D*, common to all species in solution, is related to the beam waist parameter *ω* and diffusion time *τ_D_* by the standard relation:

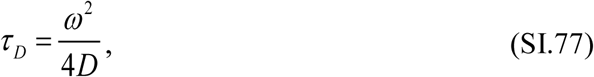

where *ω* depends on the excitation and detection channel under consideration, as well as the experiment [40]. The blinking time scale *τ_bl_*, generally due to single excited state to triplet state transitions, is a dye characteristic and is globally fit to all autocorrelation functions related to a given dye. To identify which stream an ACF corresponds to, we use the following indices for the fitted parameters:

- D excitation, D emission channel: DG (e.g.*τ*_*DG*_)
- D excitation, A emission channel: DR (e.g. *A*_*DR*_)
- A excitation, A emission channel: AR

Similarly to ACFs, the raw CCFs, *CCF_ALEX_*(*τ*), exhibit a periodic modulation, which can be cancelled out by a simple renormalization by the alternation period histograms’ CCFs (*CCF_Period_*(*τ*)):

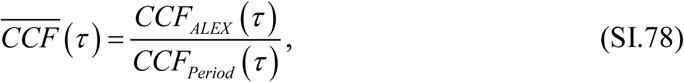

where the alternation period histograms obtained in Appendix Appendix 5 are extended over the whole duration of the experiment using their periodicity property. Because afterpulses of different detectors are uncorrelated, there is no need for further corrections, and we will simply use the notation CCF for the demodulated cross-correlation function.

CCFs were fitted with a simple 2-dimensional diffusion model:

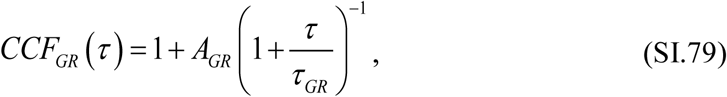

where the diffusion time *τ_GR_* (G: green, or donor, R: red, or acceptor) can in principle be obtained from the donor and acceptor channel ACF diffusion times, *τ_DG_* and *τ_DR_* by:

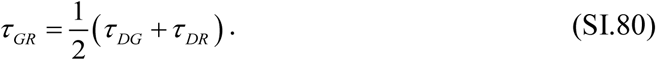

In practice, diffusion time *τ_GR_* was fitted and compared to its theoretical value, Eq. (SI.80). As for the ACF, the CCF amplitude depends in a complex manner on a variety of parameters [39], which were not computed in this study.

Fits were performed in Origin 9.1 (OriginLab Corp., Northampton, MA) using the built-in Levenberg-Marquard algorithm with statistical weights (ACF and CCF timelag range: 1 µs – 1 s).

### 12.2 Multispot FCS

The ACFs and CCF of the donor and acceptor channel signals upon donor excitation (photon streams D_ex_D_em_ and D_ex_A_em_) together with the ACF of the A_ex_A_em_ stream (acceptor excitation, acceptor channel detection) were calculated on a multitau time scale using the recorded arrival times as described in ref. [37].

To remove afterpulsing contamination from ACFs, the standard approach consisting in subtracting a component proportional to the ACF obtained with an uncorrelated, non-fluctuating sample proved ineffective [38]. Instead, we used a model function comprised of a sum of 3 exponentials in addition to a 2-dimensional diffusion contribution [10], to fit each spot’s ACF, *ACF*_*Y*_,_*i*_ (*τ* ):

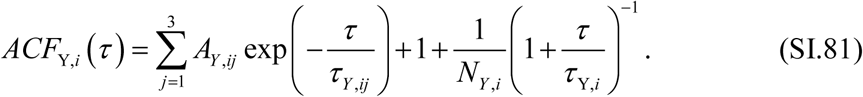

Here *i* indicates which spot’s data is fitted, and Y the detection channel under consideration (Y = G or R), while {*A*_Y,*ij*_, *τ*_*Y*_,_*ij*_ } (*j* = 1…3) are the afterpulsing components for spot *i*, channel Y. Since there is no acceptor excitation laser, there is no need to specify which excitation period is considered in the notations (the excitation period is the donor excitation period).

Note that, because of the large number of exponential involved to account for afterpulsing, we did not attempt to fit a dye triplet state blinking component. In practice, the short time scale multi-exponential fitting parameters “fit out” afterpulsing as well as triplet state blinking in a global manner.

The parameters of interest are the molecular occupancy, *N*_Y,*i*_ and the diffusion time,*τ*_Y,*i*_. Note that the true occupancy value would require a precise knowledge of both excitation and detection PSFs, which we do not possess [39]. Therefore, the term “occupancy” (and the corresponding parameters *N*_Y,*i*_) should be understood as the product of a fixed concentration (common to all spots within a sample) and an “effective” volume, which might vary from spot to spot. Comparison of “occupancies” between spots will thus report on excitation/detection volume differences between spots, not on concentration.

Afterpulsing contamination is not a problem with the CCFs, since the afterpulsing of two separate channels are uncorrelated, simplifying the fit model:

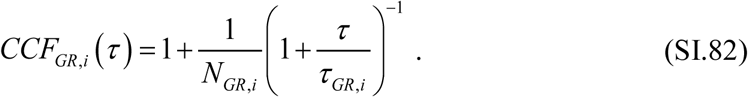

To account for a possible lateral^9^ shift *d* between donor and acceptor observation volumes, the expected functional form of the CCF is modified into [41]:

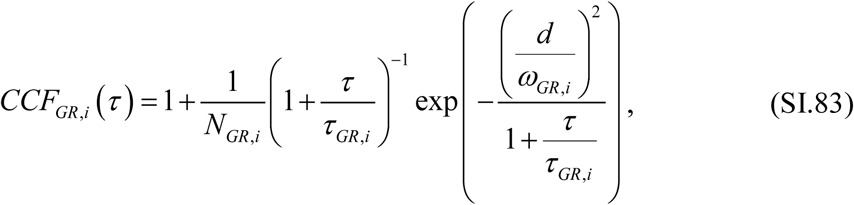

where *ω_GR, i_* is the effective Gaussian parameter of the CCF observation volume:

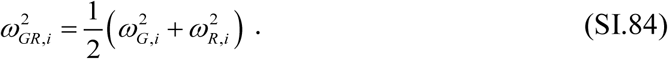

In the expression above, *ω_G, i_* (resp. *ω_R, i_*) is the Gaussian parameter of the donor (resp. acceptor) observation volume, and:

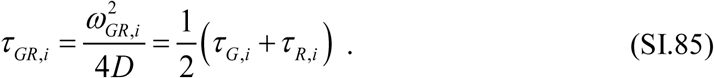

ACF and CCF curves were generated in ALiX using a standard multitau binning scheme (8 bins per chunk, 16 bins of 12,5 ns duration for the first chunk). Fits were performed in Origin 9.1 (OriginLab Corp., Northampton, MA) using the built-in Levenberg-Marquard algorithm with statistical weights (ACF timelag range: 500 ns – 1 s, CCF: 50 µs – 1 s).

Raw occupancy parameters for ACFs were corrected for background attenuation using the standard formula:

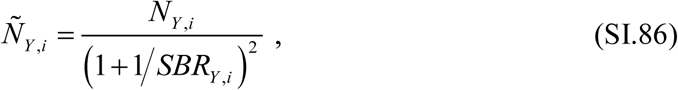

where the signal-to-background ratio is defined as:

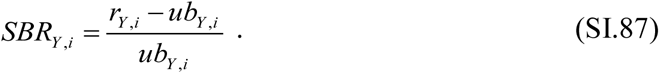

In the formula above, *r*_*Y*_,_*i*_ is the average count rate in channel *Y* for spot *i* and *ub*_*Y*_,_*i*_ is the uncorrelated background rate obtained with a buffer-only sample.

Note that Eq. (SI.81) - (SI.86) do not take into account the presence of multiple species in solution (donor only, acceptor only and doubly labeled molecules, all characterized by the same diffusion coefficient), making the occupancy parameter only useful for comparison between spots within an experiment, but not between experiments.

### 12.3 Single-spot FCS Results

The single-spot setup detection path was designed in such a way that the size of the image PSF was smaller than the detector area (50 µm) [8]. In these conditions, the detector plays no role in defining the effective observation volume, and the emission path pinhole, common to both channels, is expected to be the only source of collection efficiency reduction. In this case, the measured diffusion times increases with wavelength: *τ_AR_ > τ_DR_ > τ_DG_*. This is confirmed by the fit parameters of individual ACFs (Fig. SI-19) reported in Table SI-12.

**Fig. SI-19:**
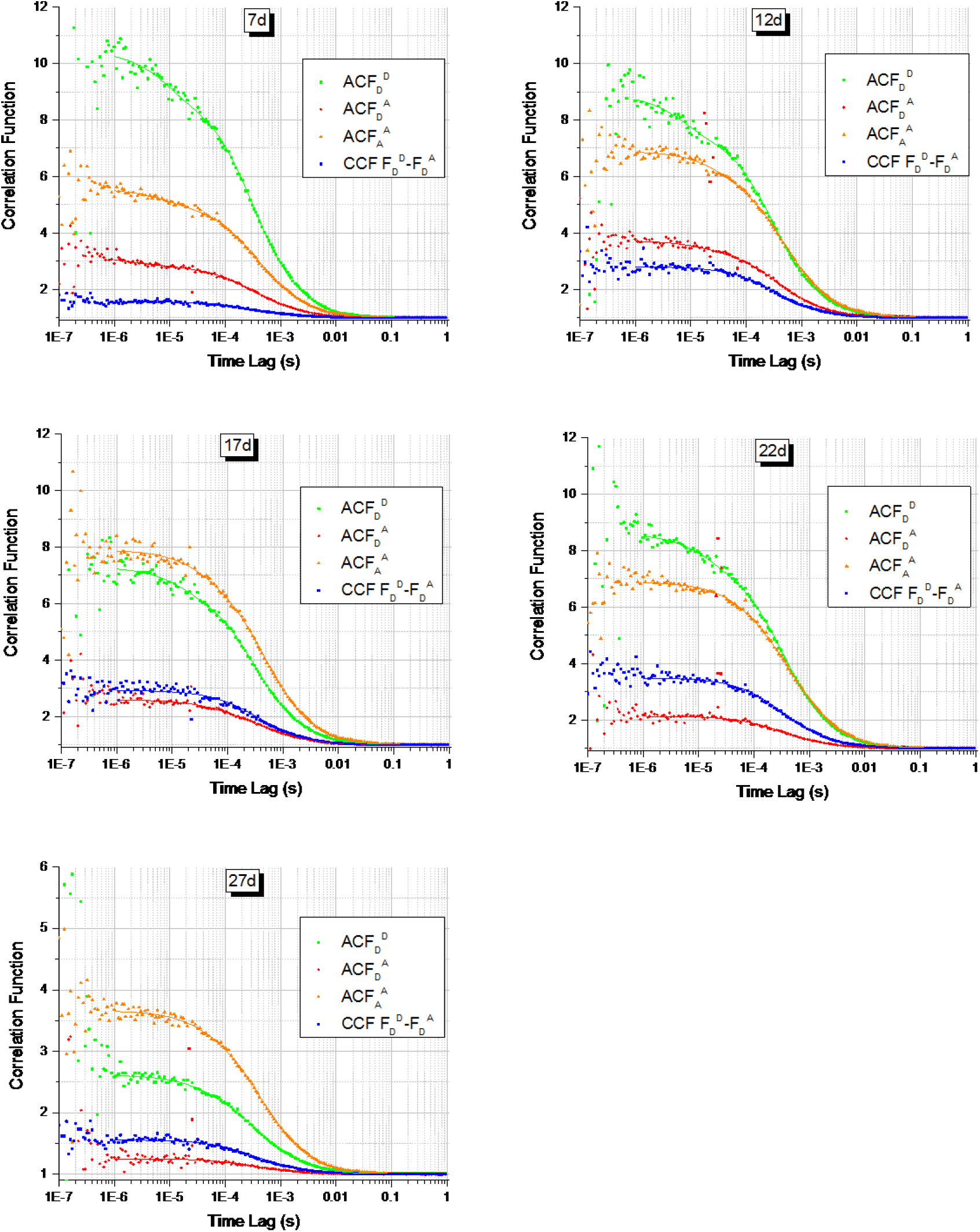
Single spot µs-ALEX FCS analysis. ACFs and CCF of the different samples used in this study. For each sample, the ACF of the 3 streams DG, DR and AR were computed and fitted with a 2-dimensional diffusion plus triplet blinking model. The CCF of the DG and DR streams was computed and fitted with a 2-dimensional diffusion plus PSF lateral offset model (Eq. (SI.83)).

**Table SI-12:** Amplitude*A*and diffusion times*τ_D_*(in µs) (Eq.) extracted from a fit of the D-excitation, D-emission channel autocorrelation function (DG), D-excitation, A-emission channel ACF (DR), and D-excitation, A-emission channel ACF (AR). Triplet state blinking (*λ* ~ 10-15%, data not shown) is observed for both donor (*τ_bl_* = 9 µs) and acceptor (*τbl* = 21 µs). (*) Only the first 400 s of dataset 17d were retained, due to the increasing background and signal level in the remainder of the trace.

| Sample | $A_{DG}$ | $\tau_{DG}$ | $A_{DR}$ | $\tau_{DR}$ | $A_{AR}$ | $\tau_{AR}$ |
| --- | --- | --- | --- | --- | --- | --- |
| 7d | 7.9 | 323 | 1.8 | 384 | 4.0 | 416 |
| 12d | 6.6 | 316 | 2.5 | 382 | 5.3 | 470 |
| 17d* | 5.6 | 306 | 1.5 | 353 | 6.3 | 447 |
| 22d | 6.8 | 327 | 1.1 | 336 | 5.5 | 460 |
| 27d | 1.5 | 355 | 0.25 | 358 | 2.6 | 396 |

Some variability in diffusion times is observed among samples, but not at a level that could lead to any suspicion of severe misalignment. However, a significant difference in all correlation amplitudes is observed for sample 27d, consistent with an increased uncorrelated background rate. Such increased background rate is most likely due to improper focus distance (lower than usual), resulting in larger scattering from the sample holder’s bottom coverslip.

### 12.4 Multispot FCS Analysis Results

Results are discussed in the main text. ACF and CCF curves can be found in Fig. SI-20.

**Fig. SI-20:**
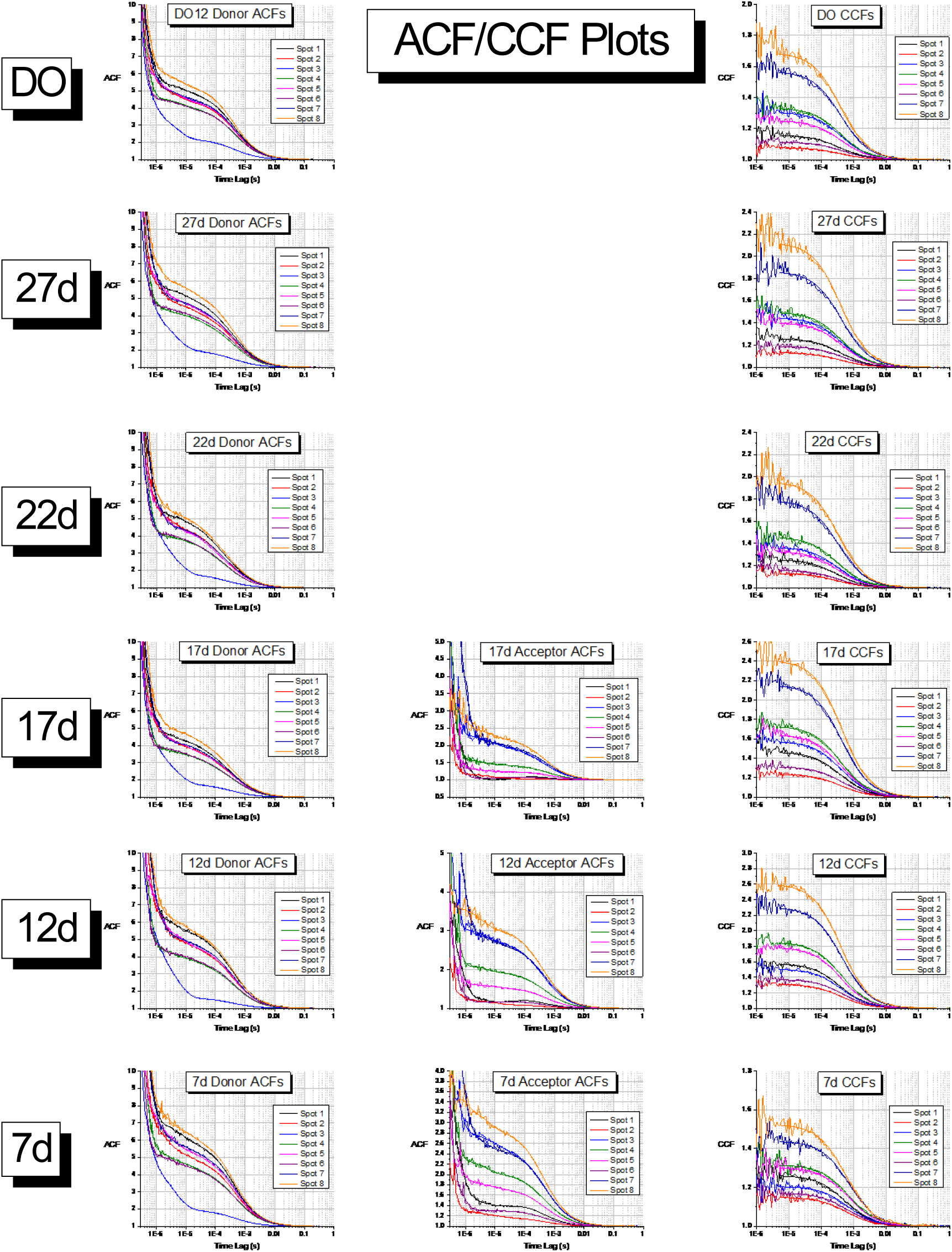
Donor and acceptor ACF and CCF plots for multispot measurements. FCS analysis for the samples studied in this work (donor only: DO, doubly-labeled: 27d to 7d). Each row in the Figure shows data from a single sample measurement (whose name is indicated to the left). Each graph in the Figure shows the donor channel autocorrelation function (ACF, left column), acceptor ACF (center column) and donor-acceptor cross-correlation function (CCF, right column) of all 8 spots in the measurement, with their corresponding fits to the models described in the main text. The acceptor ACFs for sample DO, 27d and 22d are not shown due to the low to non-existent acceptor channel signal for these samples, which prevented any reliable fit.

## Appendix 13 Samples Description

### 13.1 dsDNA FRET Samples

A set of 5 different FRET samples and their corresponding singly-labeled counterparts was used (Fig. SI-21). All samples consisted of a common 40 base-pair (bp) long doubly-labeled double-stranded DNA (dsDNA) with the donor (ATTO 550, ATTO-TEC GmbH, Heidelberg, Germany) on one strand and the acceptor (ATTO 647N, ATTO-TEC GmbH) on the other. The acceptor dye was bound to the 5' end of the top strand (Fig. DNA), while the donor dye was attached to the bottom strand at different positions from the 3’ end (7, 12, 17, 22 and 27 bp away from the acceptor, respectively). The sequence is identical to that used in previous work [29] and is designed in such a way that the environment of the donor dye is similar for all molecules, in order to minimize variations in donor quantum yield between samples.

**Fig. SI-21:**
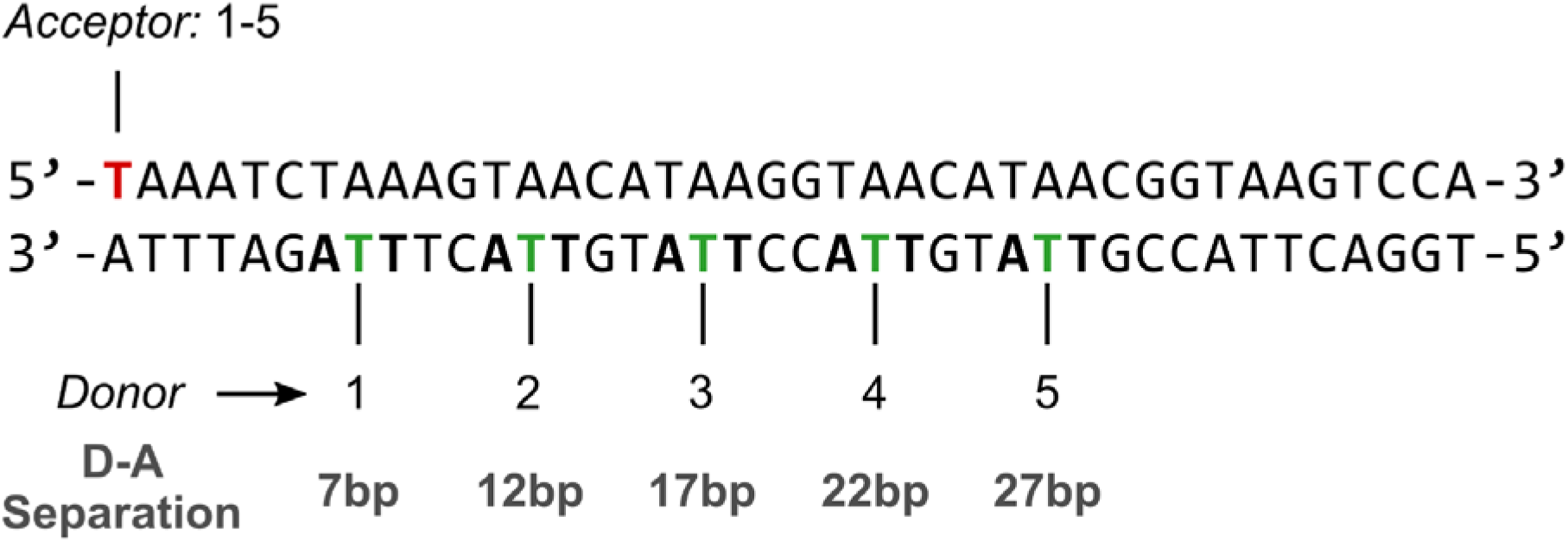
DNA sequence used in this work, with location of the dyes indicated for each sample.

Dyes were attached to a dT residue through a C6 linker using NHS-ester chemistry. Dual HPLC purified singly-labeled ssDNA samples were purchased from IDT (Coralville, IO, USA) and used without further purification.

ssDNA molecules were hybridized to their complementary strand in a 1:1 stoichiometry to form doubly-labeled samples, and with two-fold excess of unlabeled complementary strand for singly-labeled samples.

dsDNA samples were prepared with filtered, freshly prepared TE50 buffer (Tris-EDTA 50) and kept on ice until observed. 5 μl of sample at single-molecule concentration (<100 pM) were deposited in a sealed chamber consisting of a polymer gasket sandwiched between two glass coverslips. 10 to 30 min measurements were performed at room temperature (~24 ºC).

### 13.2 RNAP Transcription Samples

RNA polymerase (RNAP)-promoter initiation complex (RPO) solution was prepared as described[42]:

- 1 μl *E. coli* RNAP holoenzyme (NEB, Ipswich, MA, USA, M0551S; 1.6 μM)
- 10 μl 2X transcription buffer (80 mM HEPES KOH, 100 mM KCl, 20 mM MgCl2, 2 mM dithiotreitol (DTT), 2 mM 2-mercaptoethylamine-HCl (MEA), 0.02% Tween 20, 2 mM 5 min UV-illuminated Trolox, 200 μg/ml Bovine Serum Albumin (BSA), pH 7)
- 8 μl of water
- 1 μl of 0.1 μM lacCONS promoter DNA [43] doubly-labeled with donor and acceptor dyes labeling bases in the transcription bubble in intiation: NT(-8)ATTO647N – T(-5)ATTO550 (purchased from IBA, Germany).

RPO was then incubated in solution at 37 ºC for 30 min. To remove unreacted and nonspecifically-bound RNAP, 2 μl of 100 mg/ml Heparin-Sepharose CL-6B beads (GE Healthcare, Little Chalfont, Buckinghamshire, UK) was added to the RPO solution together with 10 μl of pre-warmed 1X transcription buffer. The mixture was incubated for 5 min at 37 ºC and centrifuged for at least 45 s at 6,000 rpm. 20 μl of the supernatant containing RPO wass transferred into a new tube containing 10 μl of pre-warmed 1X transcription buffer (heparin challenge[43, 44]).

The heparin challenged RPO solution was then incubated with 1.5 μl of 10 mM Adenylyl(3′-5′) adenosine (ApA; Ribomed, Carlsbad, CA, USA) at 37 ºC for 20 min to form a stable initially transcribed complex of up to two RNA bases (RP_ITC=2_) solution. The result was a stock of RP_ITC=2_ at promoter concentration of 2 nM. smFRET experiments were performed at 100 pM promoter concentration.

## Appendix 14 Single-Spot Setup Characterization

### 14.1 Mean Count Rates and Background Rates

Mean count rate and background rate provide a simple way to characterize a sample and its measurement. Table SI-13 reports these rates for all samples studied here (see also Fig. SI-8 for temporal variation of the total background rates).

**Table SI-13:** Background 
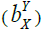
 and mean 
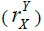
 count rates (in Hz) recorded for different photon streams XY (X: excitation period, Y: emission channel) for the 5 samples. All: All counts; D (resp. A): donor (resp. acceptor) excitation; G (resp. R): donor/green (resp. acceptor/red) channel detection. (*) For sample 17d, only the first 400 s of the measurement were used, due to the subsequent increase discussed in Fig. SI-8.

| | $b_{All}$ | $b_D^G$ | $b_D^R$ | $b_A^R$ | $r_{All}$ | $r_D^G$ | $r_D^R$ | $r_A^R$ |
| --- | --- | --- | --- | --- | --- | --- | --- | --- |
| 7d | 2,773 | 489 | 1,156 | 871 | 3,713 | 773 | 1,347 | 1,219 |
| 12d | 2,255 | 444 | 962 | 654 | 3,191 | 676 | 1,192 | 1,002 |
| 17d* | 2,149 | 554 | 843 | 588 | 3,138 | 869 | 980 | 966 |
| 22d | 2,612 | 636 | 889 | 809 | 4,095 | 1,176 | 1,039 | 1,457 |
| 27d | 3,986 | 1,523 | 996 | 1,102 | 4,905 | 1,983 | 1,061 | 1,367 |

From this data, sample 27d appears to have a significantly larger background than the other samples (*b_All_* column), the increase being particularly noticeable in the donor-excitation donor-detection stream (*b*_*D*_^*G*^ column). Sample 22d on the other hand, comes as a close second in terms of total mean count rate (*r_All_* column), while its background rate is not significantly different than most other samples. This suggest that these two samples exhibit these larger rates for distinct reasons.

When all samples are characterized by the same molecular brightness (which is a good approximation in these samples, as will be discussed later), both mean and background count rates will increase proportionally to sample concentration or excitation intensity. To distinguish which of these two effects is responsible for the observed differences, an observable which does not depend on concentration is needed.

### 14.2 Peak Burst Count Rate

The maximum peak burst count rate would seem to provide such an information. Indeed, a larger excitation intensity will result in a proportionally larger peak burst count rate, as long as saturation is negligible. Increase in the observation volume without any change in the peak excitation intensity should not affect it either. However, it is an elusive quantity to measure. In particular, it is sensitive to the presence of rare multiple molecule events, which can pass as very large single-molecule bursts.

As discussed in the Appendix 14.3, the mean value of the peak burst count rate < *r_max_* > for bursts above a threshold value *r_max,0_*, is a good alternative observable, in the sense that it does not, to a large extent, depend on the burst search and selection parameters, and scales proportionally to the peak excitation intensity. In order to obtain a similar value for all samples, the peak total count rate during donor excitation, *r_D, max_*, was chosen. Table SI-14 shows the values of < *r_D, max_* >, *r_D, max_* > 300 kHz, for all 5 samples and for two different types of burst search (APBS, *m* = 5, *F* = 6.8 and APBS, *m* = 5, *r_min_* = 27 kHz). The first search uses a common minimum SBR criterion (resulting effectively in different searches for samples characterized by different background rates), while the second uses a common burst count rate threshold criterion, equal to the threshold used for the noisiest sample in the first search.

**Table SI-14:** Average peak burst count rate during donor excitation (in kHz),*r_D, max_*, computed for bursts with*r_D, max_*>300 kHz, in two different burst searches: 1) *F* = 6.8, constant SBR burst search, using *m* = 5; 2) *rmin* = 27 kHz, constant burst count rate threshold, *m* = 5. (*) Only the first 400 s of data set 17d were retained, due to the increased background rate in the rest of the trace.

|  | 7d | 12d | 17d* | 22d | 27d |
| --- | --- | --- | --- | --- | --- |
| $F = 6.8$ | 391 | 405 | 418 | 411 | 418 |
| $r_{min} = 27$ kHz | 393 | 406 | 417 | 409 | 419 |

We conclude that the peak burst count rate during donor excitation is similar for all samples. This suggests that, as originally intended, there are no significant difference in excitation intensity and detection efficiency between measurements, and that the reason for the variability in background count rate and mean count rate needs to be attributed to other causes. This leads to the investigation of another measurement observable, the detected burst rate (number of bursts per unit time).

### 14.3 Burst Rate *n(t)*

The number of detected bursts per unit time (burst rate) depends on many parameters, including burst search and burst selection parameters. However, if sample and measurement characteristics are similar, or burst search and selection parameters are chosen in such a way as to compensate for differences, we can expect to be able to reliably compare measurements using this observable.

Table SI-15 reports the number of bursts per second detected in each measurements, using two different burst search and selection parameters.

**Table SI-15:**
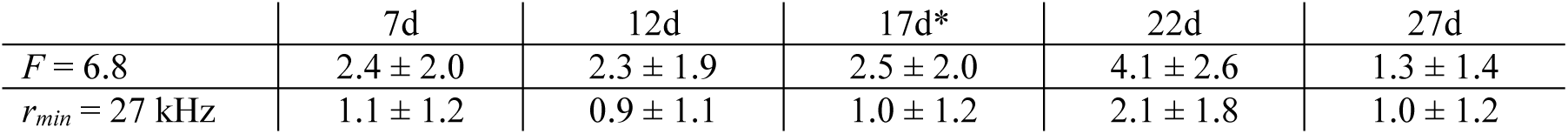
Number of bursts detected per second (± standard deviation) computed for bursts with total *γ*-corrected size *F_SBC_* ≥ 30 in two different burst searches: (i) *F* = 6.8, minimum SBR burst search, using *m* = 5; (ii) *rmin* = 27 kHz, constant burst count rate threshold, *m* = 5. (*) Only the first 400 s of data set 17d were retained, due to the increased background rate in the rest of the time trace.

|  | 7d | 12d | 17d* | 22d | 27d |
| --- | --- | --- | --- | --- | --- |
| $F = 6.8$ | $2.4 \pm 2.0$ | $2.3 \pm 1.9$ | $2.5 \pm 2.0$ | $4.1 \pm 2.6$ | $1.3 \pm 1.4$ |
| $r_{min} = 27$ kHz | $1.1 \pm 1.2$ | $0.9 \pm 1.1$ | $1.0 \pm 1.2$ | $2.1 \pm 1.8$ | $1.0 \pm 1.2$ |

The results for sample 27d are almost identical in both cases, as expected, since both burst searches are essentially equivalent for that sample (27d is the noisiest sample of the series). For all other samples, once the effect of background rates is properly compensated (*i.e.* when a constant count rate threshold is used, as used for the second row of Table SI-15), it appears that approximately twice as much bursts are detected in sample 22d than in the other samples. From this information, combined with the peak burst count rate identity among samples, we can infer that the increased mean count rate for sample 22d is likely due to a larger concentration in that sample.

What could have caused the simultaneous increases of background rate and mean count rate in sample 27d? To address this question, we need to resort to yet another independent observable obtained by fluorescence correlation analysis.

### 14.4 FCS analysis

The single-spot FCS analysis results were discussed in Appendix 11.4. There, we concluded that the 27d sample was characterized by an increased uncorrelated background rate, most likely due to laser scatter off the sample’s bottom coverslip.

## Appendix 15 Burst Statistics Definitions

While bursts are defined by their start and end photons, and can be characterized by a few quantities such as raw photon counts *F*_*D*_^*G*^, *F*_*D*_^*R*^, *F*_*A*_^*R*^, etc., as well as burst duration *ΔT*, it is possible to compute various other characteristics using the arrival time and detection channel of each photon comprising the burst. Quantities such as the proximity ratio (*PR*) and stoichiometry ratio (*SR*) or their corrected equivalents (*E_PR_*, *S* or *E*, *Sγ*) which have been introduced in Appendix 8 & Appendix 9, are used to identify and characterize FRET populations.

Other quantities are useful to characterize population concentrations, diffusivity, brightness or setup characteristics, such as relative excitation/detection efficiencies. The following quantities will be used in the remainder of this work.

### 15.1 Peak Burst Count Rate

The peak burst count rate in stream XY,
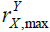
, where X represents the excitation period (D or A) and Y the detection channel (G or R), is defined by:

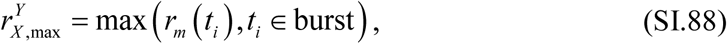

where *rm*(*ti*) is defined by Eq. (SI.25) with *c* = 1. In this work, a value of *m* = 10 was used to define the local count rate. If Y is omitted, the peak burst count rate during excitation period X is computed, including counts from both R and G channels.

Note that Eq. (SI.25) for the count rate needs to be modified in the presence of µs-ALEX alternation, as discussed in the next section.

### 15.2 Count Rate in the Presence of Laser Alternation

The previous count rate formula (Eq. (SI.25)) does not take into account which excitation period the timestamps belong to. In other words, this formula is only valid in the absence of alternation. In the presence of alternation, computing a count rate based on this formula yields problematic results if the analysis is limited to a specific excitation period, as would be the case if we wanted to compute the count rate in stream DG, for instance. This situation is illustrated in Fig. SI-22A.

**Fig. SI-22:**
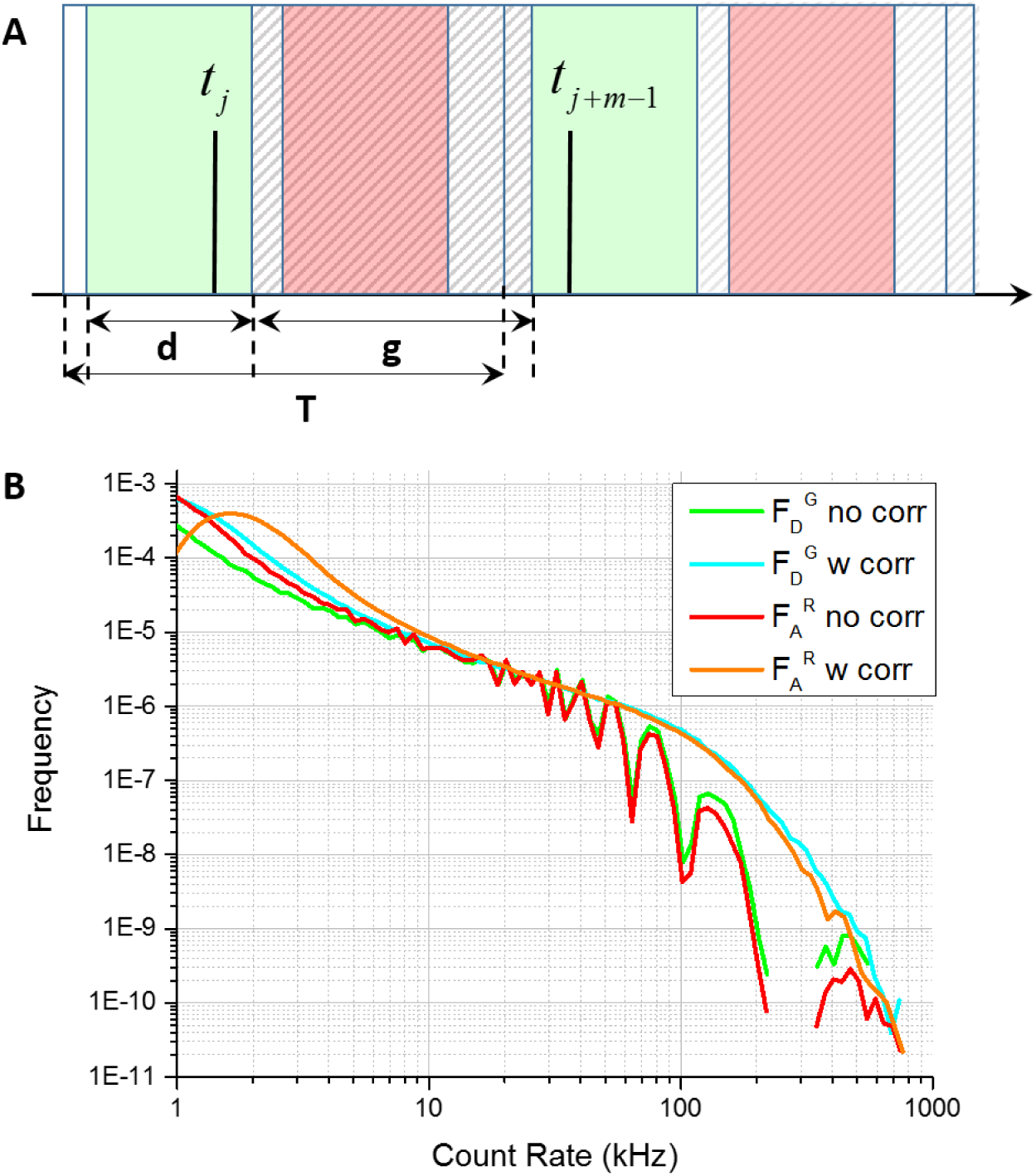
µs-ALEX corrections to the count rate. (**A**) Illustration of the gap between successive D-excitation periods (green rectangles). The total duration of the alternation period, *T*, can be decomposed into that of the D-excitation period, *d*, and its separation (or gap), *g*, from the next D-excitation period. What happens during this gap is irrelevant to compute the count rate relative to D-excitation period photons, and needs to be subtracted from the final inter-photon delay, as described in Appendix 14.1. (**B**) Illustration of the effect of using the uncorrected formula (Eq. (SI.88), green and red curves) or the µs-ALEX-corrected formula taking into account emission gaps (Eq. (SI.92), blue and orange curves). Data from file 7d. *m* = 10 was used. Notice the dips corresponding to a gap of approximately 25 µs between consecutive D-excitation (resp. A-excitation) window in the uncorrected DG (resp. AR) count rate curve (green (resp. red)).

To simplify the discussion, we only represented the first and last photons of a particular bunch of *m* consecutive photons in the DG stream, with respective timestamps *tj* and *tj+m-1*. From the figure, it is apparent that the minimum separation between both timestamps is equal to the gap between two successive D-excitation periods, *g*. Similarly, if the two timestamps were two alternation periods apart, the minimum separation would be 2*g* + *d = g + T*, where *d* is the D-excitation window duration, and *d + g = T*, the alternation period duration. More generally, if the two timestamps were *p* alternation periods apart, their minimum separation would be *g* + (*p-1*) *T*, creating artificial dips in the count rate distribution, as illustrated in Fig. SI-22B.

From Fig. SI-22A, it is clear that to ignore the gaps during which no DG photons is detected, the following *apparent* photon separation should be used:

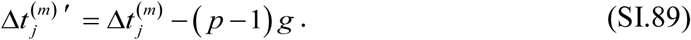

Calling the D-excitation period offset *t0*, it is easy to show that the number *p* of D-excitation periods separating the two timestamps is:

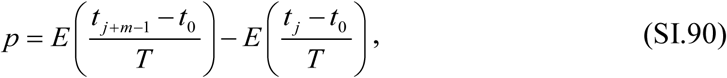

where *E(x)* designate the integer part of *x*. The final µs-ALEX-corrected formula for the count rate is thus:

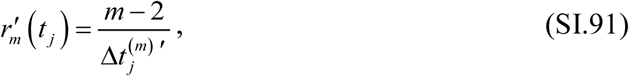

where Δ*t*_*j*_^(*m*)'^ is defined by Eq. (SI.89).

The µs-ALEX-corrected formula for the peak count rate is therefore modified into:

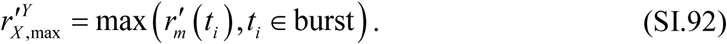

Note that in the previous derivation, the fact that donor photons were considered did not play any role, therefore the previous formula applies irrespective of which emission channel is considered. If A-excitation period photons had been considered instead, the same formulas would apply, with the simple replacement of *g* by the gap between successive A-excitation periods and *t_0_* by the A-excitation period offset. The result of these substitutions is illustrated in Fig. SI-22B, which shows continuous count rate distributions after correction, and a concomitant increase in the average count rate, as expected from the removal of gaps.

### 15.3 Proxy for the Maximum Peak Count Rate

In this section, we discuss how a *typical* peak burst count rate of a sample can be defined in a way which does not depend on the number of bursts or on the sample concentration and reflects the excitation intensity used during a measurement.

Because of the finite number of detected bursts and the stochastic nature of diffusion, the maximum peak count rate observed in the sample is not a reliable measure. Moreover, the maximum will most likely correspond to a multiple-molecule burst, and therefore artificially bias the result when using concentrated samples.

Computing the average peak count rate of *all bursts* is not a good option either, because the low peak count rate bursts (gray boxed region in Fig. SI-23A) are usually those which barely made it during the burst search or burst selection steps of the analysis, and are therefore very sensitive to search and selection parameters, including background rates.

**Fig. SI-23:**
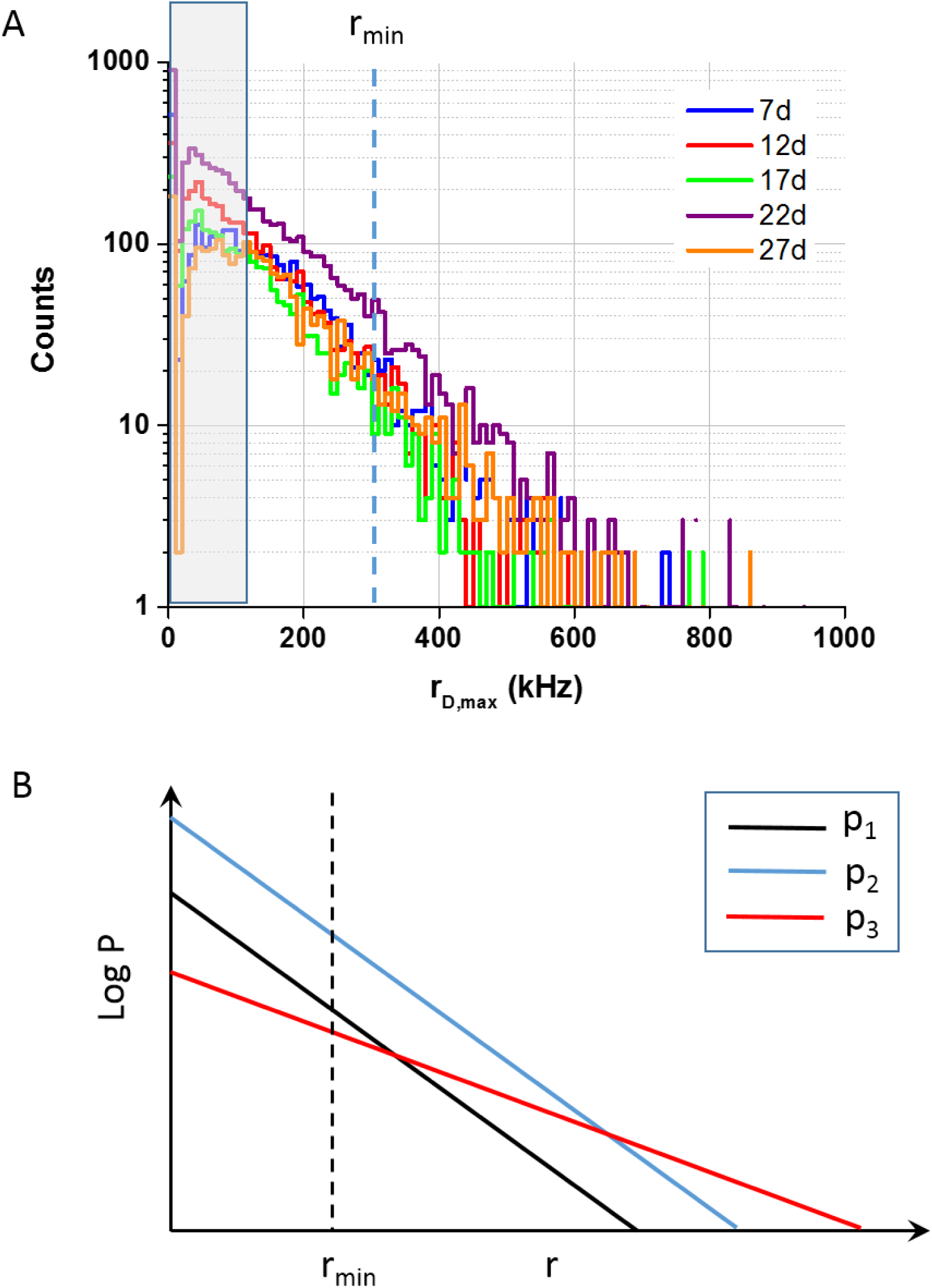
Single-spot μs-ALEX peak burst count rate histograms. Histograms of peak burst count rate (total signal during donor-excitation), whose shape should be independent of which species is observed, due to the similar brightness of all species, provided the excitation intensity and detection efficiencies are similar in all measurements. *r_min_* = 300 kHz is the minimum value used to compute the mean peak burst count rate discussed in the text. Notice that while the 22d sample measurement collected more bursts (due to a higher concentration), its average peak burst count rate was similar to that of the other samples.

To stay away from this region, a peak count rate threshold *r_min_* needs to be chosen, which is neither too close to the minimum peak count rate, nor too close to the maximum peak count rate, in order to have enough statistics to compute a reliable average.

We will now show that this bounded, mean peak count rate has the required properties.

Let’s define *p1(r)* as the peak rate histogram in a given measurement (we do not specify which stream is considered, because this is irrelevant, as long as the same choice is kept for all other measurements, as assumed here). Another measurement of the same sample resulting in more (resp. less) bursts will have a peak rate histogram *p2(r) = A p1(r)*, where *A* > 1 (resp. < 1), while a measurement of the same sample performed with a different excitation intensity (or detection efficiency), will be characterized by a peak rate histogram *p3(r) = B p1(θr)* (Fig. SI-23B).

The mean peak rate computed above *rmin* is, for each case *i* = 1, 2, 3:

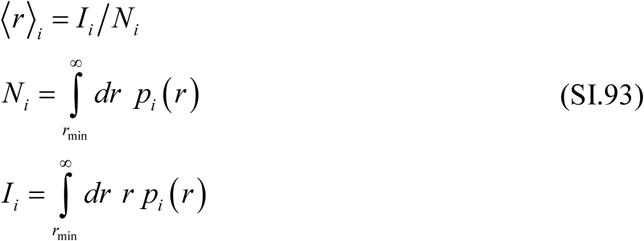

From these definitions, it follows immediately that 〈*r*〉_1_ = 〈*r*〉_2,_ no matter what functional form *pi(r)* has. This demonstrates that, as long as *rmin* is chosen in such a way that *N1* and *N2* are not too small (large enough sampling), samples characterized by the same excitation/detection properties will yield a comparable mean peak rate 〈*r.*〉

On the other hand, for the measurement characterized by either a different excitation intensity or detection efficiency, we have:

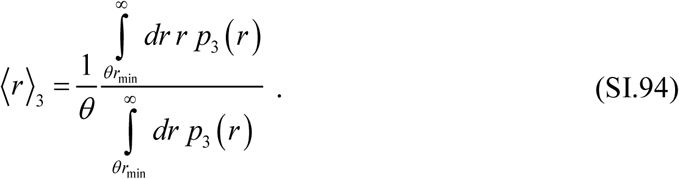

To compare this quantity to the other two, we need to assume a particular functional form. As shown in Fig. SI-23A, peak count rates distribution tails are generally well approximated by an exponential distribution:

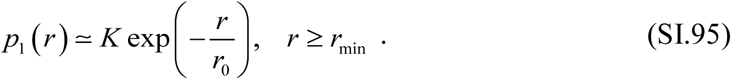

Using this approximation, we obtain:

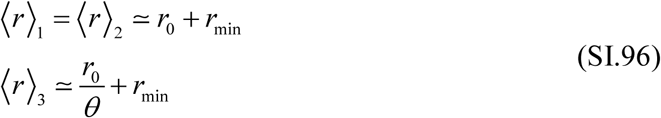

where *r_0_/θ* replaces *r_0_* in the exponential argument of Eq. (SI.95) for *p3(r)*. For *θ* < 1 (larger excitation rate, or detection efficiency), the bounded, mean peak count rate 〈*r*〉_3_ is larger than that computed for the other two cases, and the ratio of excitation intensity times detection efficiency can be obtained as:

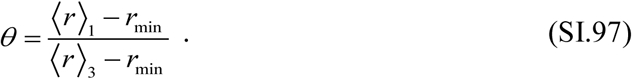

### 15.4 *γ*-Corrected Burst Size

Optimal burst selection criteria depend on the analysis purpose. If the goal of the analysis is to quantify the relative molecular concentrations of different species, using a uniform size selection criterion will in general provide a biased estimate of the relative concentrations. To understand this, we consider the experiments performed in this study, where identical molecules differing only in the number and location of dyes diffuse in an identical manner through the same excitation spot. The case of multiple spots with different characteristics is discussed later in the appendix.

In order to compare the expected burst sizes of different species, we need to go back to their theoretical expressions [29].

For *D-only molecules*, the signal (or burst size) depends on the absorption cross-section 
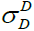
 of donor (species D) at the D-excitation wavelength, the mean excitation intensity *I*_*D*_ over the molecule’s trajectory through the excitation volume, the fluorescence quantum yield *Φ*_*D*_ of the molecule and the detection efficiency 
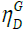
 of the setup for species D in the donor channel (Eq. (5) of Lee *et a*l. [29] with *E* = 0):

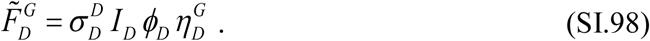

For *doubly-labeled species* (e.g. the FRET sub-population), 3 photon streams are available in µs-ALEX experiments:
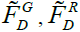
 and 
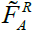

and 2 in single laser excitation experiments: 
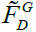
 and 
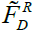

We will first discuss the latter case and come back to the µs-ALEX situation later on.

It is easy to verify from the theoretical expressions of 
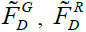
 [29]:

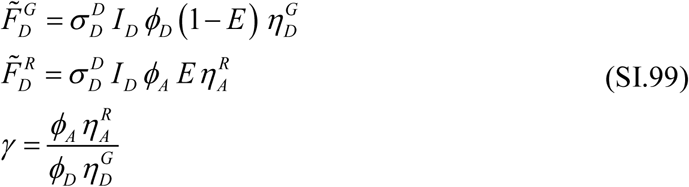

that the quantity 
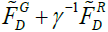
 is identical to that obtained with a D-only molecule (Eq. (SI.98)) following the same trajectory. We call the quantity:

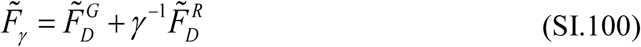

the *γ*-corrected burst size.

Note that this definition arbitrarily uses D-only as a reference, and a perfectly valid alternative definition of a corrected burst size could be chosen such that it equals that of an A-only molecule following the same trajectory:

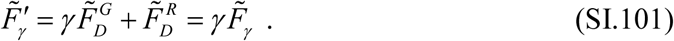

In µs-ALEX experiments, a 3^rd^ photon stream is available (
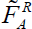
) and the uncorrected total burst size, 
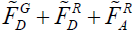
, will in general depend on which species follows a given trajectory. From the definition of 
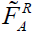
:

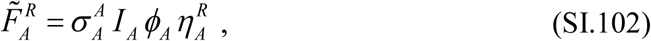

we obtain:

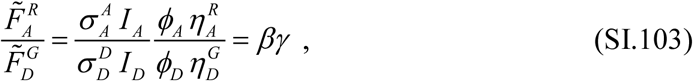

where *β* characterizing the *cross-section and excitation intensity ratio* between the two species (excitation properties, Eq. (14) in [29]):

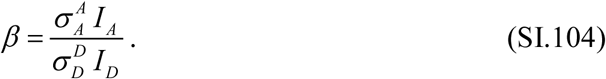

The relation between *β* and the *γ*-corrected stoichiometry ratio (Eq. (13) derived in Lee *S*γ *al.*)[29]:

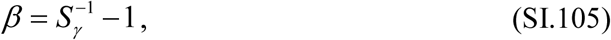

shows that, when excitation intensities are chosen such that 
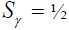
, for a doubly-labeled species, *β* = 1 and Eq. (SI.103) reduces to:

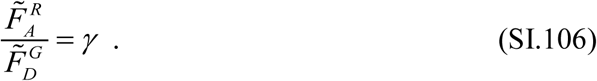

Equation (SI.103) simply states that, for the same hypothetical trajectory, the signal detected from an A-only (AO) molecule will be equal to *βγ* times that of a D-only (DO) molecule.

If *βγ* is different from 1, setting an identical signal threshold in an APBS for both species will reject more bursts of one species than the other. If the purpose of the analysis is to quantify the respective amount of D-only and A-only species, it is therefore necessary to perform a selection in which the size criterion for A-only species is *βγ* times that for the D-only species. To select all bursts with a single threshold (DO, AO and doubly-labeled species - DA), several definitions of a “corrected burst size” are possible. All involve first determining whether a burst is due to a DO, AO or DA molecule. Two examples are indicated in Table SI-16.

In Method 1, a different formula is used for different species, while in Method 2, the same formula is used to compute a “stoichiometry- and *γ*-corrected” burst size *F*_*SBC*_:

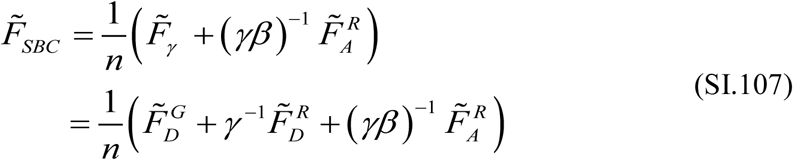

where *n* is the number of fluorophores in the molecule. Method 1 corresponds to the definition introduced for non-ALEX measurements (Eq. (SI.100)), for which *β* is undefined, and does not involve the AR stream for doubly-labeled species. Method 2 involves the AR stream, and is appropriate for studies in which it is critical to compare the relative amount of all 3 populations.

A simple way to determine which value of *n* to use for a particular burst consists in computing its stoichiometry ratio, *SR*. If *SRmin* ≤ *SR* ≤ *SR_max_*, where *SR_min_* = 0.2 and *SR_max_* = 0.85, then *n* = 2, otherwise, *n* = 1.

**Table SI-16:**
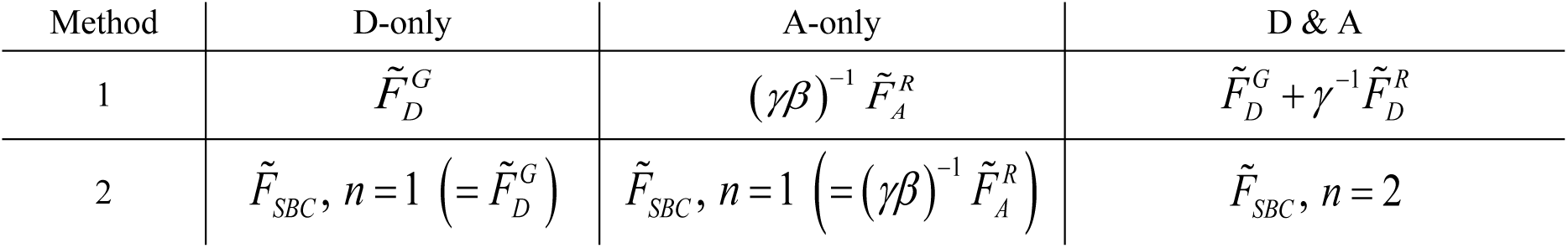
Two possible ways to compute a stoichiometry-and*γ*-corrected burst size. In Method 1, a different formula is used for each species, while in Method 2, the same formula is used, with an additional parameter *n* specifying the number of fluorophores in the molecule. For Method 2, the equivalent theoretical expression for *F*_*_SBC_*_ is indicated.

### 15.5 SBR & SNR

- *Signal-to-background ratio (SBR)* during excitation period X, defined as:

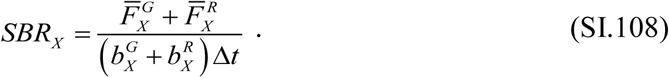

- *Signal-to-noise ratio (SNR)* during excitation period X, defined as:

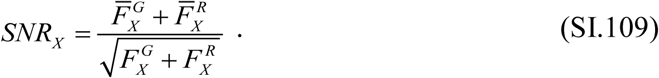

One of the main difference between the single-spot and multispot measurements is the observed background levels, due in part to the larger observation volumes of the multispot experiment, but most importantly, to the larger intrinsic (dark count) noise of some the SPADs in the arrays used in this study. While background correction takes this effect into account when computing burst statistics (such as the proximity ratio or the FRET efficiency), it is important to remember that background affects burst search results in many ways.

Here, we explore this question (also discussed in Appendix 13.2), by looking at the burst signal-to-background ratio (SBR, Eq. (SI.108)), and compare it to a related quantity, the burst signal-to-noise ratio (SNR, Eq. (SI.109)). We will focus on the donor-excitation version of these definition, since it is the only quantity common to both types of measurements.

SBR plays an important and obvious role in a minimum SBR search, where the burst count rate threshold is defined as a multiple (*F*) of the local background count rate. As discussed in Appendix 7.2.1, *F* - 1 is the minimum burst SBR, independent of the local background rate (Eq. (SI.26)). However, as illustrated in Fig. SI-24A, for a given measurement, a 15 kHz change in the burst count rate threshold can more than double (or halve) the final number of bursts after size selection. Assuming that the single-molecule signal itself is unchanged in both situations, a significant reduction of the duration of the bursts detected with higher threshold is expected, with a concomitant reduction in total burst size. This is the reason why a constant burst count rate threshold was used instead, when comparing burst duration among spots in Section 4.2.3 of the main text, since a search using a fixed burst count rate threshold, by definition, eliminates differences in threshold between experiments characterized by very different background rates.

**Fig SI-24.**
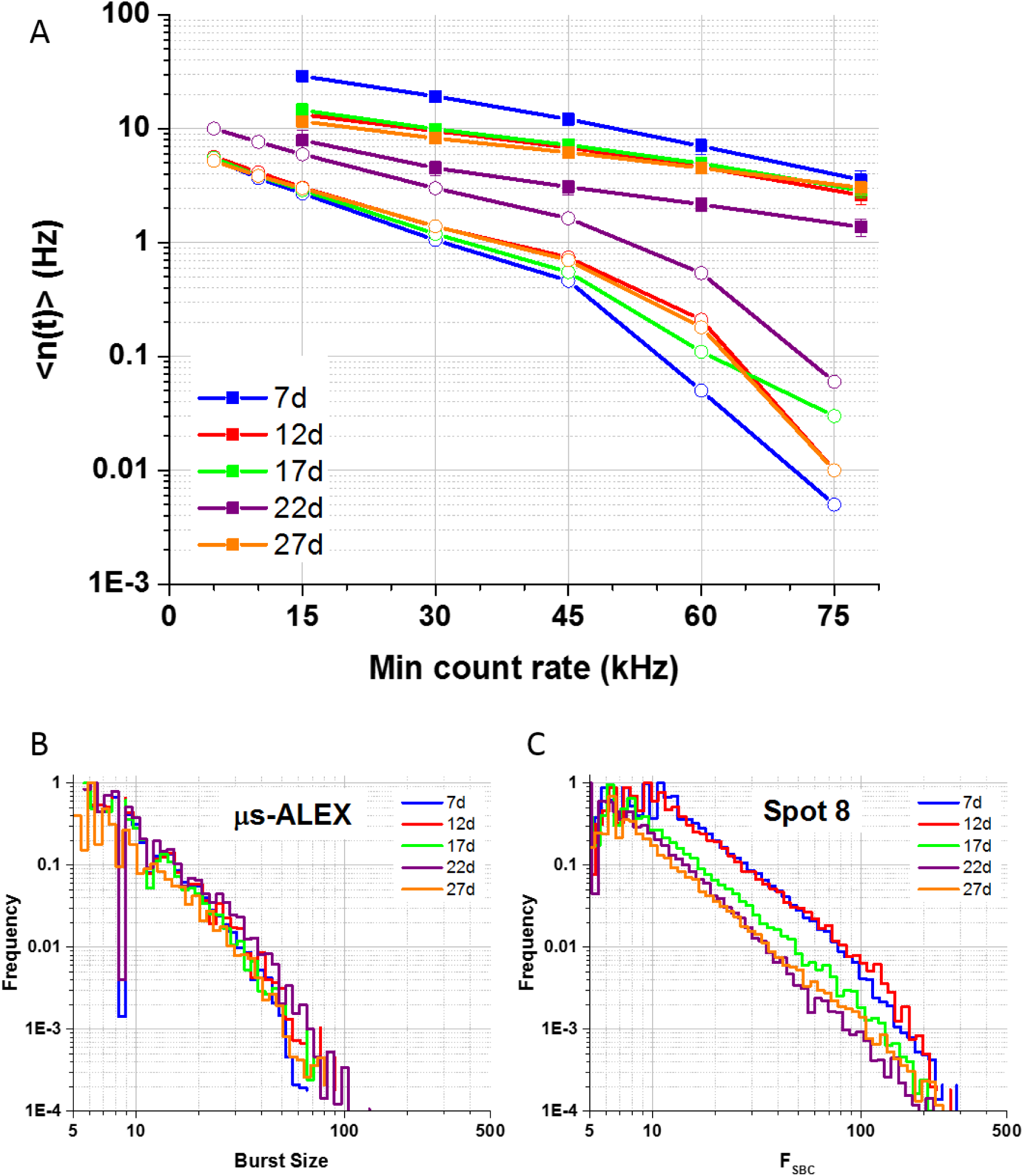
(A) Average burst rate measured for the 5 dsDNA samples, as a function of burst search count rate threshold*r_mi_*_n_. A constant *F_SBC_*≥ 30 selection criterion was used throughout (using *γ* = 1 for the single-spot µs-ALEX measurements and *γ* = 0.4 for the multispot measurements). The multispot points represent the average rate of all spots, while the error bar represents the standard deviation over all spots. Multispot analysis was limited to *r_mi_*_n_ ≥ 15 kHz due to a maximum observed background rate of 11.5 kHz. Single-spot µs-ALEX analysis was limited to *r_mi_*_n_ ≥ 5 kHz due to a maximum observed background rate of 4.9 kHz (Table SI-13). (B, C) Normalized *FSBC* (*γ*-corrected burst size) distributions observed with a donor-excitation burst search (*m* = 5, *r_min_* = 30 kHz) in the single-spot µs-ALEX measurements (B) and one of the spots (spot 8) in the multispot measurements (C). Similar distributions were observed for the others spots.

**Fig SI-25.**
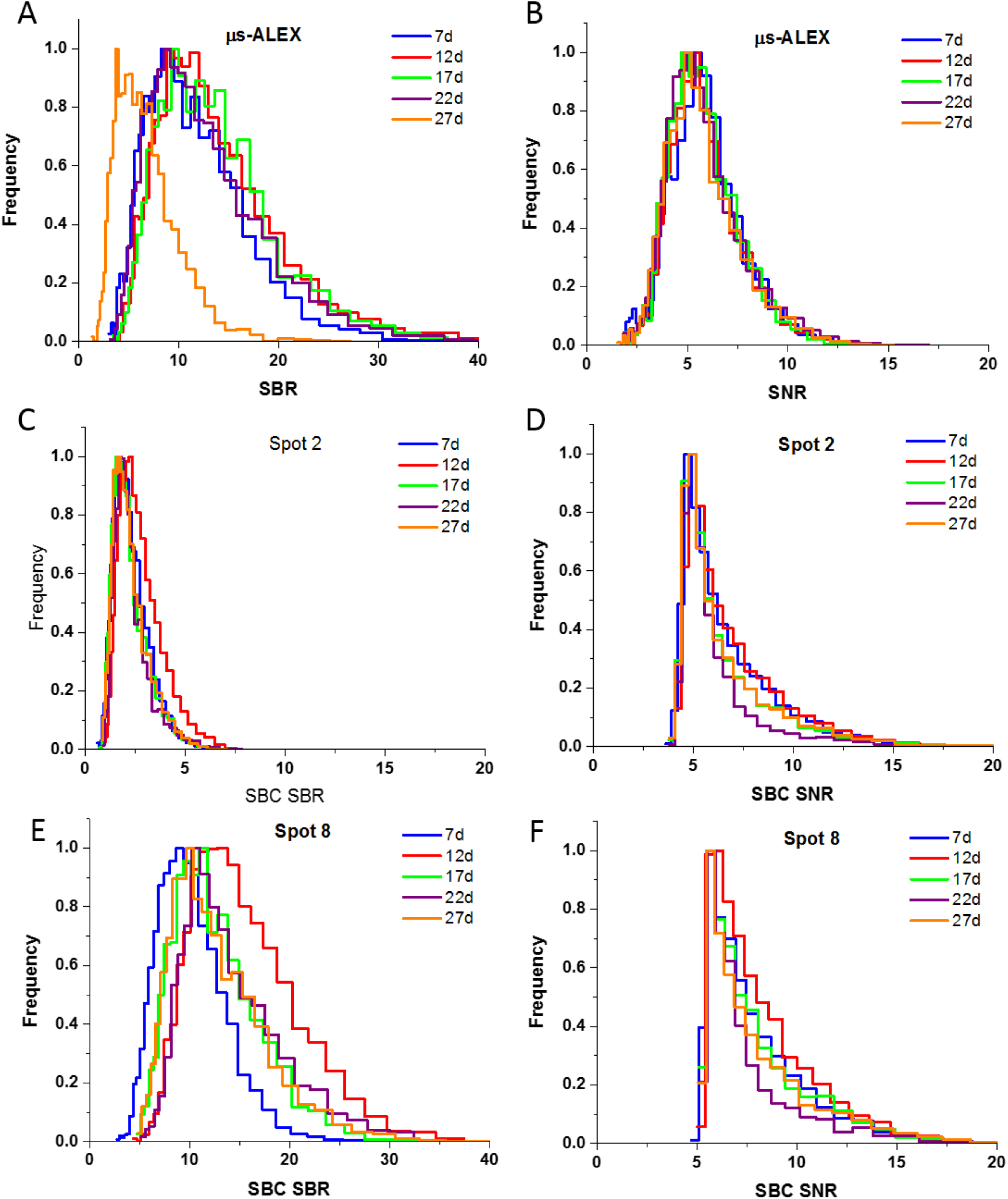
Signal-to-background-ratio (SBR) and signal-to-noise-ratio (SNR) distributions for single-spot µs-ALEX(A, B) and multispot (C-F) measurements. The single-spot µs-ALEX analysis was performed using a fixed burst search count rate threshold of 5 kHz, while the multispot analysis was performed using a fixed burst search count rate threshold of 15 kHz. 5 kHz is close to the background rate observed in sample 27d (Table SI-13), which results in a lower SBR than the for other samples (A), but similar SNR (B). Similarly, 15 kHz is close to the background observed in Spot 2, which results in a low SBR for all samples (C), while the SNR is relatively independent on the sample (D). By comparison, spot 8, which has a background rate close to 7 kHz lower than Spot 2 (Table 1, main text), exhibits much larger SBR (E), but comparable SNR (F). The single-spot µs-ALEX signal used in this study is the donor-excitation signal. For the multispot calculation, the *γ*-corrected signal *F*_*SBC*_ and *γ*-corrected background was used.

However, using a search with fixed burst count rate threshold does not guarantee a minimum value for the burst SBR anymore. This is illustrated in Fig. SI-25A (and SI-25B), where the SBR (and SNR) distributions are shown for the different samples studied with the single-spot µs-ALEX setup, using a common count rate threshold of 5 kHz (corresponding to the largest background rate among all samples, or leftmost point in Fig. SI-24A). The sample with the largest background rate (27d) is now characterized by the smallest SBR, while the signal itself (and hence the SNR) is almost identical for all samples.

A comparable conclusion can be drawn from the comparison of the noisiest (spot 2) and quietest spot (spot 8) of the multispot experiments. Using a common count rate threshold of 15 kHz (corresponding to the largest background rate among all spots and measurements, or leftmost point in Fig. SI-24A), the observed SBR in spot 2 (Fig. SI-25C) is systematically and markedly lower than that observed in spot 8 (Fig. SI-25E), while both exhibit similar SNR (Fig. SI-25D & SI-25F).

In summary, a constant SBR search, while convenient to compare samples characterized by similar background levels, can result in biased statistics in situations where background levels are quite different. The use of a constant rate threshold appears preferable in most situations, the optimal value depending on the peak count rate of the sample, the desired number of bursts and minimal SNR.

### 15.6 Burst rate *n(t)*

The number of single-molecule bursts detected per unit time during a measurement is a simple observable allowing to compare measurements or setups, and specifically to demonstrate the increased throughput of multispot data acquisition. However, its definition depends on parameters used for burst search and burst selection (*e.g.* Appendix 13.3), as well as sample and setup characteristics, such as concentration, excitation power, background rate or detection efficiency. For instance, comparing the number of bursts larger than a certain size in two different measurements characterized by different observation volumes (but otherwise similar characteristics), will obviously result in a larger number of bursts for the experiment characterized by the largest observation volume. As discussed previously, the single-spot µs-ALEX measurements and the multispot measurements were characterized by similar peak excitation power, but larger observation volumes and lower acceptor detection efficiency in the multispot experiments. Since the absolute concentration of the different samples was not determined, the only comparison which can be performed is between burst rates computed with similar burst selection criteria (*γ*-corrected burst size *F*_*SBC*_ ≥ 30), and for the multispot experiments, using searches with identical burst thresholds across spots. The results obtained for various burst search count rate thresholds *r_min_* are plotted in Fig. SI-24A, together with the corresponding rates measured in the single-spot µs-ALEX experiments (open symbols). The measured burst rates are significantly lower in the single-spot µs-ALEX experiment (up to one order of magnitude in some cases), in part because of the overall larger bursts observed in the multispot experiments (Fig. SI-24B & C), but also possibly due to concentration differences.

Fig. SI-24A also demonstrates the exponential dependence of the detected number of bursts on the burst search count rate threshold *r_min_*. While this suggests a simple way to increase the number of detected bursts (by reducing the count rate threshold), decreasing the count rate threshold has negative consequences on other metrics, as was discussed in the previous section.

## Appendix 16 Simple model of the DNA double-helix with two labels

### 16.1 Model description

To compare the FRET efficiencies measured for each sample in the µs-ALEX experiments to those measured in the multispot experiments, no model is necessary. However, since the dsDNA samples used in this work are simple, and a similar comparison was performed in previous studies, the corrected FRET efficiencies were compared to a simple model of double stranded helix, with fluorophores attached to base pairs located at different positions along its sequence.

The model used in this work is slightly different from the popular “Clegg model”[45] and offers some additional degrees of freedom, as illustrated in Fig. SI-26 & SI-27. The double helix is modeled by the two helices passing through the phosphorus atoms (P) of each nucleic acid base (helix 1: green, helix 2: red), supported by a 1 nm radius cylinder. The angle *H*_1_Ω*H*_2_ = ∆*ϕ* between the two helices and the cylinder axis (Fig. SI-26A) in any plane perpendicular to the axis is set to 2.31 radians (132.4 º), based on a distance of 18.3 Å between phosphorus atoms of opposite strands and a 10 Å helix radius. We define a P-P rung as the segment connecting the phosphorus atoms of opposite stands. The rise between successive base pairs/rung is 3.34 Å along the direction of the cylinder axis, and its rotation, 360/10.5 º (10.5 base pairs per turn). So far, this model does not have any free parameters (although the previous values can be adjusted to alternative values if needed).

**Fig. SI-26:**
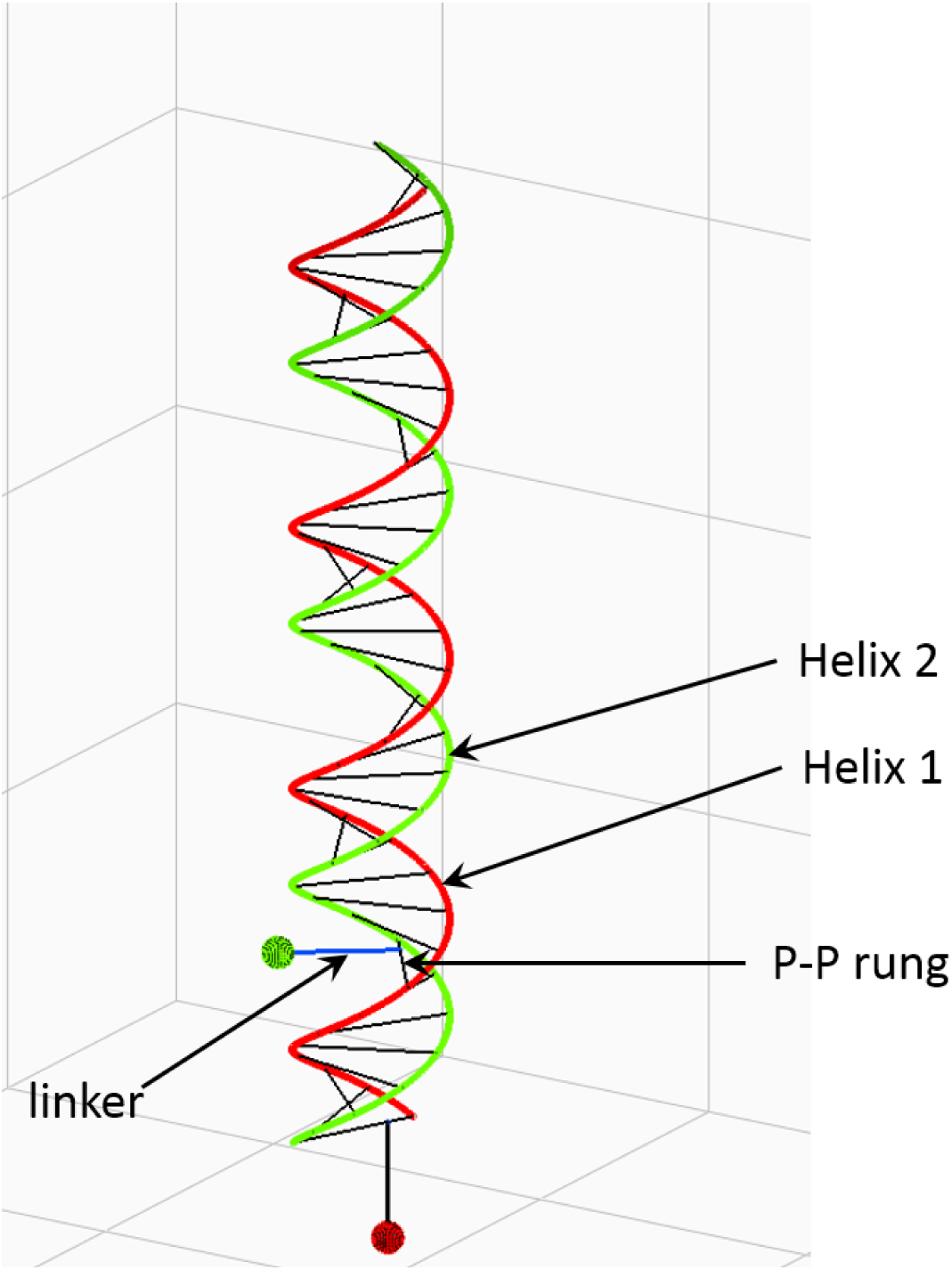
DNA double-helix model schematic. perspective view of a double stranded DNA molecule labeled with an acceptor dye (red, bottom) at position 1 and a donor dye (green) at position 7. The molecule is represented by the two phosphate backbone helices and rungs connecting opposite phosphorus atoms, and one linker per dye, each attached to its corresponding rung at a variable location.

To model a dye attached to a base, we introduce three parameters per dye (in practice we used the same set for both dyes): the distance *d* from helix 1 of the “attachment” point *S* (Fig. SI-27B), the inclination *ψ* of the vector connecting the dye D to S, and the distance *L* of the dye from this attachment point (Fig. SI-26B). Note that in reality, the dye’s linker is attached to a nucleic acid base, which is itself located somewhere in the base pair plane (in other words, the linker attachment point is different from S). However, the exact interpretation of the 3 parameters *d*, *L* and *ψ* does not really matter as long as the bases to which the dyes in different samples are attached are always the same, as is the case in our study (T = thymine). While we chose the “linker” to be located in the base pair plane (or more precisely, the phosphorus plane) for internal labels, an additional off-plane degree of freedom could be conceivably added to account for subtle conformation details, in the specific case of the terminal acceptor dye (red ball in Fig. SI-26), the linker was rotated 90º off-plane, as this is the most symmetrical (or neutral) conformation of all.

**Fig. SI-27:**
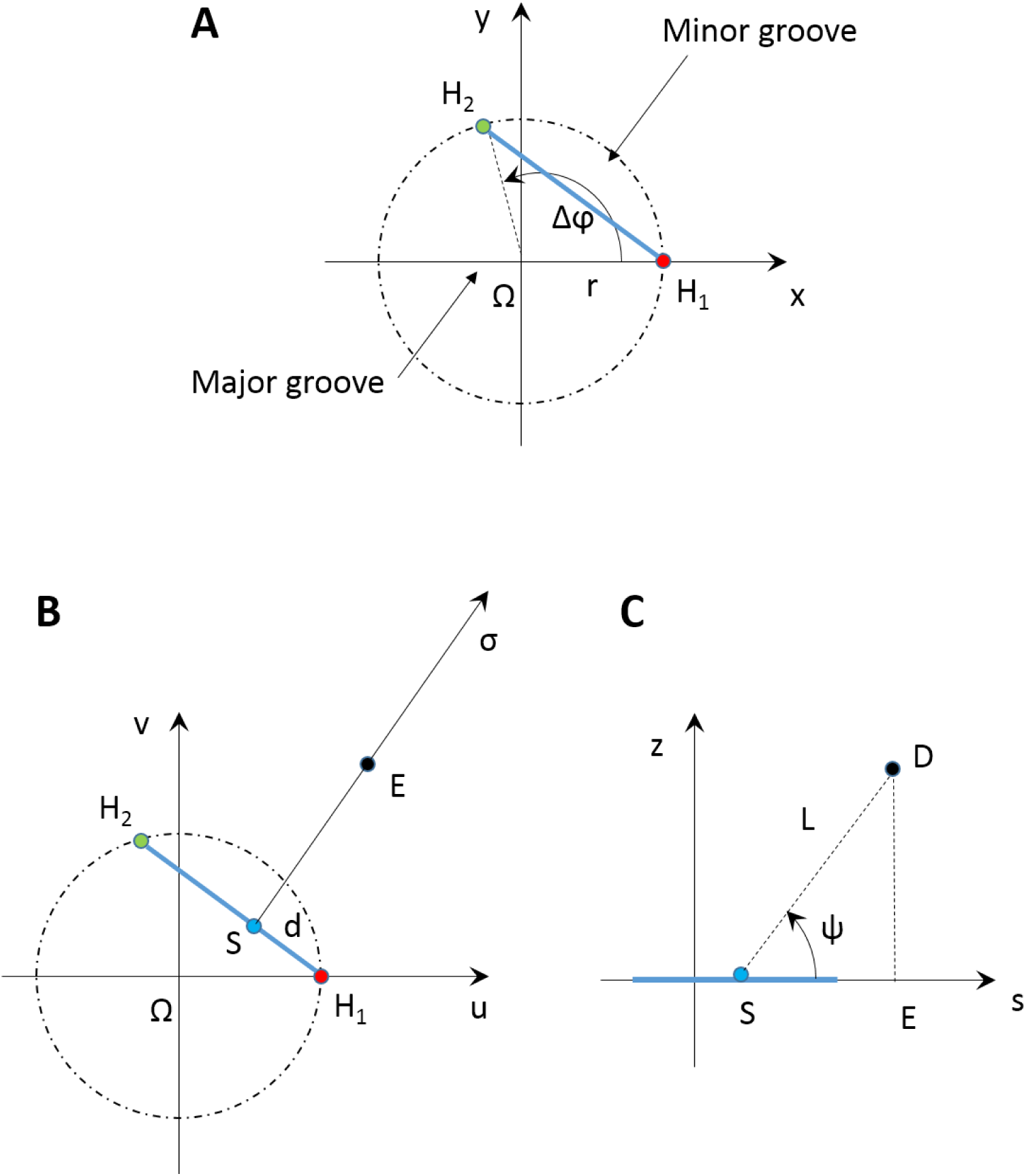
DNA double-helix model details. (**A**) Top view of a base pair plane, showing the two phosphorus atoms(H1 and H2), the DNA molecule axis (Ω). (**B**) Definition of parameter *d* (location of the linker attachment point on the rung) and projection E of the dye center in the rung plane. (**C**) Lateral view and definition of thinclination angle *ψ* and linker length *L*.

With these definitions in hand, it is simple to derive equations for the position of dyes attached along the DNA stands and their respective distances.

### 16.2 Model parameterization

Let’s assume (Figure SI-27A) that the first phosphorus of helix 1 is at position (*r*, 0, 0) and the first phosphorus of helix 2 is at position (*r* sin(*ΔΦ*), **r** cos(*ΔΦ*), 0). The phosphorus atoms of base pair *i* will have coordinates:

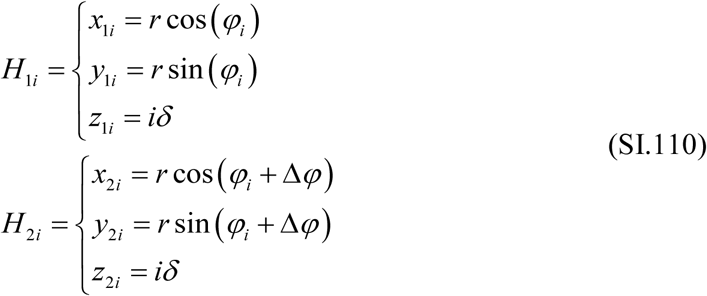

where *ϕ*_*i*_  ^2^_*n*_^*π*^*i* and *n* is the number of base pairs per turn (*n* = 10.5, *δ* = 0.334 nm, *r* = 1 nm in B-DNA).

The segment connecting the two opposite phosphorus atoms of a given base pair is a convenient reference to localize the average dye position *D* with respect to the double helix. We will project this point on the plane perpendicular to the double helix and defined by the two phosphorus atoms, calling this projection *E* (Figure SI-27B). The orthogonal projection *S* of that point on the *H1H2* segment will be our reference point for the dye localization and can be defined by:

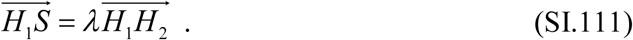

In general, 0 < *λ* < ½, but depending on the circumstances, a negative value of *λ* or *λ* > ½ is conceivable.

The dye’s average position will be specified by its distance *L* to that “anchor” point (this is not a linker length, but is loosely related to it) and its angle *ψ* with respect to the projection plane (in general, symmetry will impose *ψ* = 0, but some situations may require other values).

Noting *σ* the unit vector along segment SE defined as:

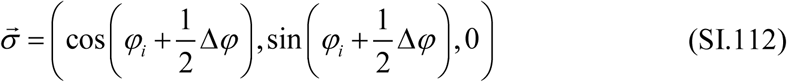

we have:

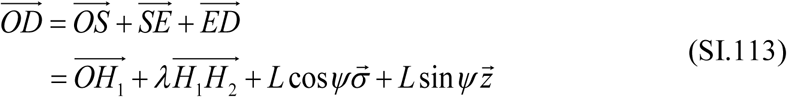

And finally:

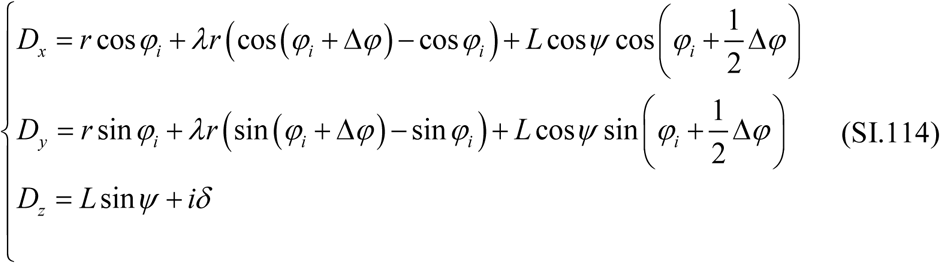

Note that in this expression *L* can be positive or negative, depending on whether the dye is located on the minor groove or the major groove side of the double helix.

So far, we have not clarified which strand the dye is attached to. This is because the above definition is dependent on which strand is considered as reference. However, if one dye (A) is attached to one strand and the other (B) to the opposite strand (and in a similar fashion), then for symmetry reasons, we need to impose:

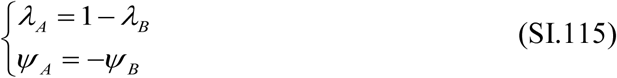

The distance between two dyes (and the corresponding FRET efficiency) is easily obtained from their individual coordinates (Eq. (SI.114)).

The following parameters (A: acceptor dye, D: donor dye) were used in Fig. 6 of the main text: *dA= dB= d* = 7.13 Å (or equivalently, *λA* = 0.39, *λD* = 0.61), *LA= LB= L* = 12.8 Å, *ψA* =-90º, *ψD* = 0º.

## Footnotes

1 In this document, we will indifferently refer to detector counts as “photons” or “counts”, since there is no way to distinguish between them at the individual count level.

2 Note that Eq. (SI.19) uses the fact that the “gap” regions discussed in Appendix 5 have a negligible influence.

3 Other possibilities are, for instance, to use only donor channel photons (“donor emission burst search”, or D_em_BS: all photons recorded by the donor channel, irrespective of the laser alternation period they were emitted in), or acceptor channel photons (“acceptor emission burst search”, or A_em_BS: same as above, but for acceptor photons), donor excitation period photons (“donor excitation burst search”, or D_ex_BS: all photons recorded by either channels but limited to the donor laser on period), or acceptor excitation period photons (“acceptor excitation burst search”, or A_ex_BS: same as above, but for the acceptor laser). Any of these burst searches can be combined using logic AND or OR operations. The AND combination of two searches simply keeps the burst parts overlapping in both searches (in other words, their intersection, ∩). The OR combination of two searches returns all bursts found in either search, fusing any two overlapping bursts into a single, larger burst (their union, ⋃). For instance, the “dual channel burst search” (DCBS) defined in ref. 21 corresponds to the intersection (OR operation) of a donor excitation and an acceptor excitation burst searches: DCBS = D_ex_BS AND A_ex_BS.

4 Starting with this appendix, notations in this Supporting Material depart from the simpler notations introduced the main text in order to better distinguish several related quantities.

5 In the main text, this quantity is called *n_Dem_* (or *n_D_*). Quantity *n_Aem_* (or *n_A_*) in the main text corresponds to 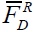 in this document. 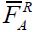 corresponds to *n_AA_*. *F*_*D*_^*R*^:

6 Eq. (SI.50) only applies to zero-FRET samples. For a *doubly-labeled sample characterized by a non-zero FRETefficiency E*, it needs to be modified into: 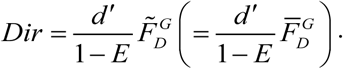 This expression reduces to Eq. (SI.50) for *E* = 0, but is indeterminate for a 100% FRET efficiency sample (*E* = 1), for which 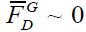 . In this case, Eq. (SI.55) is a preferable expression. Obviously, a donor-only molecule will not contribute any direct acceptor excitation signal (*i.e. Dir* = 0 for these molecules).

7 By construction, *N*_*D*_ + *N*_*A*_ = *F*_*D*_.

8 At this stage, it is possible to apply the donor leakage and acceptor direct excitation corrections to obtain an *E_PR_*-histogram rather than a PRH

9 A similar term accounting for a potential vertical shift can be included, but, as for the diffusion component of the ACF which was not necessary in this study, the ACF is much less sensitive to this effect than to a lateral shift.

## References

1. Deniz AA, Dahan M, Grunwell JR, Ha T, Faulhaber AE, Chemla DS, Weiss S, Schultz PG. Single-pair fluorescence resonance energy transfer on freely diffusing molecules: observation of Förster distance dependence and subpopulations. Proceedings of the National Academy of Sciences USA. 1999;96(7):3670–5. doi: 10.1073/pnas.96.7.3670.

2. DahanM, DenizAA, HaTJ, ChemlaDS, SchultzPG, WeissS. Ratiometric measurement and identification of single diffusing molecules. Chemical Physics. 1999;247(1):85–106. doi: 10.1016/S0301-0104(99)00132-9.

3. KapanidisAN, MargeatE, LaurenceTA, DooseS, HoSO, MukhopadhyayJ, KortkhonjiaE, MeklerV, EbrightRH, WeissS. Retention of transcription initiation factor sigma(70) in transcription elongation: Single-molecule analysis. Molecular Cell. 2005;20(3):347–56. doi: 10.1016/j.molcel.2005.10.012.

4. RoyR, HohngS, HaT., A practical guide to single-molecule FRET., Nature Methods. 2008;5(6):507–16. doi: 10.1038/nmeth.1208.

5. LeveneMJ, KorlachJ, TurnerSW, FoquetM, CraigheadHG, WebbWW. Zero-mode waveguides for single-molecule analysis at high concentrations. Science. 2003;299(5607):682–6. doi: 10.1126/science.1079700.

6. RondelezY, TressetG, TabataKV, ArataH, FujitaH, TakeuchiS, NojiH. Microfabricated arrays of femtoliter chambers allow single molecule enzymology. Nature Biotechnology. 2005;23(3):361–5. doi: 10.1038/nbt1072.

7. RhoadesE, GussakovskyE, HaranG. Watching proteins fold one molecule at a time. Proceedings of the National Academy of Sciences USA. 2003;100(6):3197–202. doi: 10.1073/pnas.2628068100.

8. KimJY, KimC, LeeNK., Real-time submillisecond single-molecule FRET dynamics of freely diffusing molecules with liposome tethering. Nature Communications. 2015;6:6992. doi: 10.1038/ncomms7992.

9. Colyer RA, Scalia G, Rech I, Gulinatti A, Ghioni M, Cova S, Weiss S, Michalet X., High-throughput FCS using an LCOS spatial light modulator and an 8x1 SPAD array. Biomedical Optics Express. 2010;1(5):1408–31. doi: 10.1364/BOE.1.001408.

10. ColyerRA, ScaliaG, VillaFA, GuerrieriF, TisaS, ZappaF, CovaS, WeissS, MichaletX. Ultrahigh-throughput single-molecule spectroscopy with a 1024 SPAD. Proceedings of SPIE. 2011;7905:790503. doi: 10.1364/BOE.1.001408.

11. BuchholzJ, KriegerJW, MocsárG, KreithB, CharbonE, VámosiG, KebschullU, LangowskiJ. FPGA implementation of a 32x32 autocorrelator array for analysis of fast image series. Optics Express. 2012;20(16):17767–82. doi: 10.1364/OE.20.017767.

12. GhioniM, GulinattiA, RechI, ZappaF, CovaS. Progress in silicon single-photon avalanche diodes. Ieee Journal of Selected Topics in Quantum Electronics. 2007;13(4):852–62. doi: 10.1109/jstqe.2007.902088.

13. IngargiolaA, ColyerRA, KimD, PanzeriF, LinR, GulinattiA, RechI, GhioniM, WeissS, MichaletX. Parallel multispot smFRET analysis using an 8-pixel SPAD array. Proceedings of SPIE. 2012;8228:82280B. doi: 10.1117/12.909470.

14. IngargiolaA, PanzeriF, SarkoshN, GulinattiA, RechI, GhioniM, WeissS, MichaletX. 8-spot smFRET analysis using two 8-pixel SPAD arrays. Proceeding of SPIE. 2013;8590:85900E. doi: 10.1117/12.2003704.

15. CuccatoA, AntonioliS, GulinattiA, LabancaI, RechI, GhioniM. Compact 32-channel time-resolved single-photon detection system. Proceedings of SPIE. 2013;8773:8777030. doi: 10.1117/12.2017355.

16. GulinattiA, RechI, MaccagnaniP, GhioniM. A 48-pixel array of single photon avalanche diodes for multispot single molecule analysis. Proceeding of SPIE. 2013;8631:86311D. doi: 10.1117/12.2003984.

17. IngargiolaA, LaurenceT, BoutelleR, WeissS, MichaletX. Photon-HDF5: An Open File Format for Timestamp-Based Single-Molecule Fluorescence Experiments. Biophysical Journal. 2016;110(1):26–33. doi: 10.1016/j.bpj.2015.11.013.

18. IngargiolaA, LernerE, ChungS, WeissS, MichaletX. FRETBursts: An Open Source Toolkit for Analysis of Freely-Diffusing Single-Molecule FRET. PLoS ONE. 2016;11(8). doi: 10.1371/journal.pone.0160716.

19. PanzeriF, IngargiolaA, LinRR, SarkhoshN, GulinattiA, RechI, GhioniM, CovaS, WeissS, MichaletX., Single-moleculeFRET experiments with a red-enhanced custom technology SPAD. Proceedings of SPIE. 2013;8590. doi: 10.1117/12.2003187.

20. RechI, ResnatiD, MarangoniS, GhioniM, CovaS. Compact-eight channel photon counting module with monolithic array detector. Proceedings of SPIE. 2007;6771:677113. doi: 10.1117/12.749483.

21. RechI, MarangoniS, ResnatiD, GhioniM, CovaS. Multipixel single-photon avalanche diode array for parallel photon counting applications. Journal of Modern Optics. 2009;56(2):326–33. doi: 10.1080/09500340802318309.

22. HillesheimLN, MullerJD. The photon counting histogram in fluorescence fluctuation spectroscopy with non-ideal photodetectors. Biophysical Journal. 2003;85(3):1948–58. doi: 10.1016/S0006-3495(03)74622-0.

23. RechI, IngargiolaA, SpinelliR, LabancaI, MarangoniS, GhioniM, CovaS. Optical crosstalk in single photon avalanche diode arrays: a new complete model. Optics Express. 2008;16(12):8381–94. doi: 10.1364/oe.16.008381.

24. EggelingC, BergerS, BrandL, FriesJR, SchafferJ, VolkmerA, SeidelCAM. Data registration and selective single-molecule analysis using multi-parameter fluorescence detection. Journal of Biotechnology. 2001;86(3):163–80. doi: 10.1016/S0168-1656(00)00412-0.

25. LeeNK, KapanidisAN, KohHR, KorlannY, HoSO, KimY, GassmanN, KimSK, WeissS. Three-color alternating-laser excitation of single molecules: Monitoring multiple interactions and distances. Biophysical Journal. 2007;92(1):303–12. doi: 10.1529/biophysj.106.093211.

26. NirE, MichaletX, HamadaniKM, LaurenceTA, NeuhauserD, KovchegovY, WeissS., Shot-noise limited single-molecule FRET histograms: Comparison between theory and experiments. Journal of Physical Chemistry B. 2006;110(44):22103–24. doi: 10.1021/jp063483n.

27. SisamakisE, ValeriA, KalininS, RothwellPJ, SeidelCAM., Accurate single-molecule FRET studies using multiparameter fluorescence detection. Methods in Enzymology. 2010;475:455–514. doi: 10.1016/s0076-6879(10)75018-7.

28. KudryavtsevV, SikorM, KalininS, MokranjacD, SeidelCAM, LambDC., CombiningMFD and PIE for Accurate Single-Pair Forster Resonance Energy Transfer Measurements. Chemical Physics Physical Chemistry. 2012;13(4):1060–78. doi: 10.1002/cphc.201100822.

29. KapanidisAN, LeeNK, LaurenceTA, DooseS, MargeatE, WeissS. Fluorescence-aided molecule sorting: Analysis of structure and interactions by alternating-laser excitation of single molecules. Proceedings of the National Academy of Sciences USA. 2004;101(24):8936–41. doi: 10.1073/pnas.0401690101.

30. LeeNK, KapanidisAN, WangY, MichaletX, MukhopadhyayJ, EbrightRH, WeissS., AccurateFRET, Measurements within Single Diffusing Biomolecules Using Alternating-Laser Excitation. Biophysical Journal. 2005;88(4):2939–53. doi: 10.1529/biophysj.104.054114.

31. GopichIV, SzaboA. Theory of the statistics of kinetic transitions with application to single-molecule enzyme catalysis. Journal of Chemical Physics. 2006;124(15). doi: 10.1063/1.2180770.

32. FriesJR, BrandL, EggelingC, KollnerM, SeidelCAM. Quantitative identification of different single molecules by selective time-resolved confocal fluorescence spectroscopy. Journal of Physical Chemistry A. 1998;102(33):6601–13. doi: 10.1021/jp980965t.

33. MichaletX, ColyerRA, ScaliaG, IngargiolaA, LinR, MillaudJE, WeissS, SiegmundOHW, TremsinAS, VallergaJV, ChengA, LeviM, AharoniD, ArisakaK, VillaF, GuerrieriF, PanzeriF, RechI, GulinattiA, ZappaF, GhioniM, CovaS. Development of new photon-counting detectors for single-molecule fluorescence microscopy. Philosophical Transactions of the Royal Society B. 2013;368(1611): 20120035. doi: 10.1098/rstb.2012.0035.

34. MichaletX, IngargiolaA, ColyerRA, ScaliaG, WeissS, MaccagnaniP, GulinattiA, RechI, GhioniM. Silicon photon counting avalanche diodes for single-molecule fluorescence spectroscopy. Journal of Selected Topics in Quantum Electronics. 2014;20(6):3804420. doi: 0.1109/JSTQE.2014.2341568.

35. MichaletX, ChengA, AntelmanJ, SuyamaM, ArisakaK, WeissS. Hybrid photodetector for single-molecule spectroscopy and microscopy. Proceedings of SPIE. 2008;6862:68620F. doi: 10.1117/12.763449.

36. WeidemannT, WachsmuthM, TewesM, RippeK, LangowskiJ. Analysis of ligand binding by two-colour fluorescence cross-correlation spectroscopy. Single Molecules. 2002;3(1):49–61. doi: 10.1002/1438-5171(200204)3:1<49::Aid-Simo49>3.3.Co;2–K.

37. IngargiolaA. Applying Corrections in Single-Molecule FRET. Biorxiv. 2016:preprint. doi: 10.1101/083287.

38. KongXX, NirE, HamadaniK, WeissS., Photobleaching pathways in single-molecule FRET experiments. Journal of the American Chemical Society. 2007;129(15):4643–54doi: 10.1021/ja068002s.

39. DertingerT, PachecoV, von der HochtI, HartmannR, GregorI, EnderleinJ. Two-focus fluorescence correlation spectroscopy: A new tool for accurate and absolute diffusion measurements. Chemical Physics Physical Chemistry. 2007;8(3):433–43. doi: 10.1002/cphc.200600638.

40. KimS, StreetsAM, LinRR, QuakeSR, WeissS, MajumdarDS. High-throughput single-molecule optofluidic analysis. Nature Methods. 2011;8:242–5. doi: 10.1038/NMETH.1569.

41. MeklerV, KortkhonjiaE, MukhopadhyayJ, KnightJ, RevyakinA, KapanidisAN, NiuW, EbrightYW, LevyR, EbrightRH., Structural organization of bacterial RNA polymerase holoenzyme and the RNA polymerase-promoter open complex. Cell. 2002;108(5):599–614. doi: 10.1016/S0092-8674(02)00667-0.

42. MukhopadhyayJ, KapanidisAN, MeklerV, KortkhonjiaE, EbrightYW, EbrightRH. Translocation of sigma(70) with RNA polymerase during transcription: fluorescence resonance energy transfer assay for movement relative to DNA. Cell. 2001;106(4):453–63. Epub 2001/08/30. doi: 10.1016/S0092-8674(01)00464-0.

43. LernerE, ChungS, AllenBL, WangS, LeeJ, LuWS, GrimaudLW, IngargiolaA, MichaletX, AlhadidY, BorukhovS, StrickTR, TaatjesDJ, WeissS. A, backtracked and paused transcription initiation intermediate of Escherichia Coli RNA polymerase. Proceedings of the National Academy of Sciences USA. 2016;113(43):E6562–E71. Epub 2016/10/11. doi: 10.1073/pnas.1605038113.

44. AbbondanzieriEA, ShaevitzJW, BlockSM., Picocalorimetry of transcription by RNA polymerase. Biophysical Journal. 2005;89(6):L61–L3. doi: 10.1529/biophysj.105.074195.

45. AbbondanzieriEA, GreenleafWJ, ShaevitzJW, LandickR, BlockSM., Direct observation of base-pair stepping by RNA polymerase. Nature. 2005;438(7067):460–5doi: 10.1038/nature04268.

46. AdelmanK, La Porta A, SantangeloTJ, LisJT, RobertsJW, WangMD., Single molecule analysis of RNA polymerase elongation reveals uniform kinetic behavior. Proceedings of the National Academy of Sciences USA. 2002;99(21):13538–43. doi: 10.1073/pnas.212358999.

47. MargeatE, KapanidisAN, TinnefeldP, WangY, MukhopadhyayJ, EbrightRH, WeissS. Direct observation of abortive initiation and promoter escape within single immobilized transcription complexes. Biophysical Journal. 2006;90(4):1419–31. doi: 10.1529/biophysj.105.069252.

48. KapanidisAN, LaurenceTA, LeeNK, MargeatE, KongXX, WeissS. Alternating-laser excitation of single molecules. Accounts of Chemical Research. 2005;38(7):523–33. doi: 10.1021/ar0401348.

49. MullerBK, ZaychikovE, BrauchleC, LambDC. Pulsed interleaved excitation. Biophysical Journal. 2005;89(5):3508–22. doi: 10.1529/biophysj.105.064766.

50. GulinattiA, RechI, MaccagnaniP, GhioniM, CovaS., Improving the performance of Silicon Single Photon Avalanche Diodes. Proceedings of SPIE. 2011;8033:803302. doi: 10.1117/12.883863.

51. GulinattiA, PanzeriF, RechI, MaccagnaniP, GhioniM, CovaS., Planar silicon SPADs with improved photon detection efficiency. Proceedings of SPIE. 2011;7945:79452P. doi: 10.1117/12.874685.

52. OikawaH, SuzukiY, SaitoM, KamagataK, AraiM, TakahashiS. Microsecond dynamics of an unfolded protein by a line confocal tracking of single molecule fluorescence. Scientific Reports. 2013;3. doi: 10.1038/srep02151.

53. WunderlichB, NettelsD, BenkeS, ClarkJ, WeidnerS, HofmannH, PfeilSH, SchulerB. Microfluidic mixer designed for performing single-molecule kinetics with confocal detection on timescales from milliseconds to minutes. Nature Protocols. 2013;8(8):1459–74. doi: 10.1038/nprot.2013.082.

## Appendix References

1. Ingargiola A, Laurence T, Boutelle R, Weiss S, Michalet X. Photon-HDF5: An Open File Format for Timestamp-Based Single-Molecule Fluorescence Experiments. Biophysical Journal. 2016;110(1):26–33. doi: 10.1016/j.bpj.2015.11.013.

2. Ingargiola A, Lerner E, Chung S, Panzeri F, Gulinatti A, Rech I, Ghioni M, Weiss S, Michalet X. smFRET μs-ALEX: 5 dsDNA samples. 2016. doi: 10.6084/m9.figshare.1098961.

3. Ingargiola A, Lerner E, Chung S, Panzeri F, Gulinatti A, Rech I, Ghioni M, Weiss S, Michalet X. 8-spot smFRET measurements on 6 dsDNA samples. 2016. doi: 10.6084/m9.figshare.1098962.

4. Ingargiola A, Lerner E, Chung S, Panzeri F, Gulinatti A, Rech I, Ghioni M, Weiss S, Michalet X. RNAP, Promoter Escape Kinetics Data Files. 2016. doi: 10.6084/m9.figshare.3810930.

5. Ingargiola A, Lerner E, Chung S, Panzeri F, Gulinatti A, Rech I, Ghioni M, Weiss S, Michalet X., Afterpulsing ACF, Correction Files. 2016. doi: 10.6084/m9.figshare.3817062.

6. Ingargiola A, Lerner E, Chung S, Panzeri F, Gulinatti A, Rech I, Ghioni M, Weiss S, Michalet X., Jupyter Notebooks for Multispot single-molecule FRET: high-throughput analysis of freely diffusing molecules. 2016. doi: not yet attributed.

7. Ingargiola A, Lerner E, Chung S, Panzeri F, Gulinatti A, Rech I, Ghioni M, Weiss S, Michalet X. ALiX Scripts. 2016. doi: 10.6084/m9.figshare.3839427.

8. Panzeri F, Ingargiola A, Lin RR, Sarkhosh N, Gulinatti A, Rech I, Ghioni M, Cova S, Weiss S, Michalet X., Single-molecule FRET experiments with a red-enhanced custom technology SPAD. Proceedings of SPIE. 2013;8590. doi: 10.1117/12.2003187.

9. Zhang ZC, You Z, Chu DP. Fundamentals of phase-only liquid crystal on silicon (LCOS) devices. Light-Science & Applications. 2014;3. doi: 10.1038/lsa.2014.94.

10. Colyer RA, Scalia G, Rech I, Gulinatti A, Ghioni M, Cova S, Weiss S, Michalet X., High-throughput FCS using an LCOS spatial light modulator and an 8x1 SPAD array. Biomedical Optics Express. 2010;1(5):1408–31. doi: 10.1364/BOE.1.001408.

11. Ingargiola A, Panzeri F, Sarkosh N, Gulinatti A, Rech I, Ghioni M, Weiss S, Michalet X. 8-spot smFRET analysis using two 8-pixel SPAD arrays. Proceeding of SPIE. 2013;8590:85900E. doi: 10.1117/12.2003704.

12. Rech I, Marangoni S, Resnati D, Ghioni M, Cova S. Multipixel single-photon avalanche diode array for parallel photon counting applications. Journal of Modern Optics. 2009; 56(2):326–33. doi: 10.1080/09500340802318309.

13. Guerrieri F, Tisa S, Tosi A, Zappa F., Two-Dimensional SPAD, Imaging Camera for Photon Counting. IEEE Photonics Journal. 2010;2(5):759–74. doi: 10.1109/jphot.2010.2066554.

14. Colyer RA, Scalia G, Villa FA, Guerrieri F, Tisa S, Zappa F, Cova S, Weiss S, Michalet X. Ultrahigh-throughput single-molecule spectroscopy with a 1024 SPAD. Proceedings of SPIE. 2011;7905:790503. doi: 10.1364/BOE.1.001408.

15. Michalet X, Ingargiola A, Colyer RA, Scalia G, Weiss S, Maccagnani P, Gulinatti A, Rech I, Ghioni M. Silicon photon counting avalanche diodes for single-molecule fluorescence spectroscopy. Journal of Selected Topics in Quantum Electronics. 2014;20(6):3804420. doi: 0.1109/JSTQE.2014.2341568.

16. Ingargiola A, Colyer RA, Kim D, Panzeri F, Lin R, Gulinatti A, Rech I, Ghioni M, Weiss S, Michalet X. Parallel multispot smFRET analysis using an 8-pixel SPAD array. Proceedings of SPIE. 2012;8228:82280B. doi: 10.1117/12.909470.

17. Rech I, Ingargiola A, Spinelli R, Labanca I, Marangoni S, Ghioni M, Cova S. Optical crosstalk in single photon avalanche diode arrays: a new complete model. Optics Express. 2008;16(12):8381–94. doi: 10.1364/oe.16.008381.

18. Restelli A, Rech I, Maccagnani P, Ghioni M, Cova S. Monolithic silicon matrix detector with 50 mu m photon counting pixels. Journal of Modern Optics. 2007;54(2–3):213–23. doi: 10.1080/09500340600790121.

19. Gopich IV, Szabo A. Theory of the statistics of kinetic transitions with application to single-molecule enzyme catalysis. Journal of Chemical Physics. 2006;124(15). doi: 10.1063/1.2180770.

20. Gopich IV, Szabo A., Theory of Photon Counting in Single-Molecule Spectroscopy. Theory and Evaluation of Single-Molecule Signals: World Scientific; 2008. p. 181–244.

21. Ingargiola A, Lerner E, Chung S, Weiss S, Michalet X. FRETBursts: An Open Source Toolkit for Analysis of Freely-Diffusing Single-Molecule FRET. PLoS ONE. 2016; 11(8). doi: 10.1371/journal.pone.0160716.

22. Press WH, Teukolsky SA, Vetterling WT, Flannery BP., Numerical Recipes in C. The Art of Scientific Computing. 2 ed. Cambridge: Cambridge University Press; 1992.

23. Nir E, Michalet X, Hamadani KM, Laurence TA, Neuhauser D, Kovchegov Y, Weiss S., Shot-noise limited single-molecule FRET histograms: Comparison between theory and experiments. Journal of Physical Chemistry B. 2006;110(44):22103–24. doi: 10.1021/jp063483n.

24. Fries JR, Brand L, Eggeling C, Kollner M, Seidel CAM. Quantitative identification of different single molecules by selective time-resolved confocal fluorescence spectroscopy. Journal of Physical Chemistry A. 1998;102(33):6601–13. doi: 10.1021/jp980965t.

25. Michalet X, Colyer RA, Scalia G, Ingargiola A, Lin R, Millaud JE, Weiss S, Siegmund OHW, Tremsin AS, Vallerga JV, Cheng A, Levi M, Aharoni D, Arisaka K, Villa F, Guerrieri F, Panzeri F, Rech I, Gulinatti A, Zappa F, Ghioni M, Cova S. Development of new photon-counting detectors for single-molecule fluorescence microscopy. Philosophical Transactions of the Royal Society B. 2013;368(1611): 20120035. doi: 10.1098/rstb.2012.0035.

26. Hoffmann A, Nettels D, Clark J, Borgia A, Radford SE, Clarke J, Schuler B., Quantifying heterogeneity and conformational dynamics from single molecule FRET of diffusing molecules: recurrence analysis of single particles (RASP). Physical Chemistry Chemical Physics. 2011;13(5):1857–71. doi: 10.1039/c0cp01911a.

27. Sigworth FJ, Sine SM., Data Transformations for Improved Display and Fitting of Single-Channel Dwell Time Histograms. Biophysical Journal. 1987;52(6):1047–54. doi: 10.1016/S0006-3495(87)83298-8.

28. Carr DB, Olsen AR, White D., Hexagon Mosaic Maps for Display of Univariate and Bivariate Geographical Data. Cartography and Geographic Information Systems. 1992;19(4):228. doi: 10.1559/152304092783721231.

29. Lee NK, Kapanidis AN, Wang Y, Michalet X, Mukhopadhyay J, Ebright RH, Weiss S., Accurate FRET, Measurements within Single Diffusing Biomolecules Using Alternating-Laser Excitation. Biophysical Journal. 2005;88(4):2939–53. doi: 10.1529/biophysj.104.054114.

30. Ingargiola A. Applying Corrections in Single-Molecule FRET. Biorxiv. 2016:preprint. doi: 10.1101/083287.

31. Jones MC, Marron JS, Sheather SJ. A brief survey of bandwidth selection for density estimation. Journal of the American Statistical Association. 1996;91(433):401–7. doi: 10.2307/2291420.

32. Antonik M, Felekyan S, Gaiduk A, Seidel CAM. Separating structural heterogeneities from stochastic variations in fluorescence resonance energy transfer distributions via photon distribution analysis. Journal of Physical Chemistry B. 2006;110(13):6970–8. doi: 10.1021/jp057257.

33. Churchman LS, Flyvbjerg H, Spudich JA. A non-Gaussian distribution quantifies distances measured with fluorescence localization techniques. Biophysical Journal. 2006;90(2):668–71. doi: 10.1529/biophysj.105.065599.

34. Krichevsky O, Bonnet G. Fluorescence correlation spectroscopy: the technique and its applications. Reports on Progress in Physics. 2002;65(2):251–97. doi: 10.1088/0034-4885/65/2/203.

35. Hess ST, Webb WW. Focal volume optics and experimental artifacts in confocal fluorescence correlation spectroscopy. Biophysical Journal. 2002;83(4):2300–17. doi: 10.1016/S0006-3495(02)73990-8.

36. Enderlein J, I G, Patra D, Fitter J. Art and artefacts of fluorescence correlation spectroscopy. Current Pharmaceutical Biotechnology. 2004;5(2):155–61. doi: 10.2174/1389201043377020.

37. Laurence TA, Fore S, Huser T. Fast, flexible algorithm for calculating photon correlations. Optics Letters. 2006;31(6):829–31. doi: 10.1364/OL.31.000829.

38. Zhao M, Jin L, Chen B, Ding Y, Ma H, Chen DY. Afterpulsing and its correction in fluorescence correlation spectroscopy experiments. Applied Optics. 2003;42(19):4031–6. doi: 10.1364/Ao.42.004031.

39. Schwille P, Meyer-Almes FJ, Rigler R. Dual-color fluorescence cross-correlation spectroscopy for multicomponent diffusional analysis in solution. Biophysical Journal. 1997;72(4):1878–86. doi: 10.1016/S0006-3495(97)78833-7.

40. Thompson NL. Fluorescence Correlation Spectroscopy. Topics in Fluorescence Spectroscopy, Volume 1: Techniques. 1. New York: Plenum Press; 1991. p. 337–78.

41. Weidemann T, Wachsmuth M, Tewes M, Rippe K, Langowski J. Analysis of ligand binding by two-colour fluorescence cross-correlation spectroscopy. Single Molecules. 2002;3(1):49–61. doi: 10.1002/1438-5171(200204)3:1<49::Aid-Simo49>3.3.Co;2–K.

42. Lerner E, Chung S, Allen BL, Wang S, Lee J, Lu WS, Grimaud LW, Ingargiola A, Michalet X, Alhadid Y, Borukhov S, Strick TR, Taatjes DJ, Weiss S. A, backtracked and paused transcription initiation intermediate of Escherichia Coli RNA polymerase. Proceedings of the National Academy of Sciences USA. 2016;113(43):E6562–E71. Epub 2016/10/11. doi: 10.1073/pnas.1605038113.

43. Kim S, Streets AM, Lin RR, Quake SR, Weiss S, Majumdar DS. High-throughput single-molecule optofluidic analysis. Nature Methods. 2011;8:242–5. doi: 10.1038/NMETH.1569.

44. Kapanidis AN, Margeat E, Ho SO, Kortkhonjia E, Weiss S, Ebright RH., Initial transcription by RNA polymerase proceeds through a DNA-scrunching mechanism. Science. 2006;314(5802):1144–7. Epub 2006/11/18. doi: 10.1126/science.1131399.

45. Clegg RM. Fluorescence resonance energy transfer and nucleic acids. Methods in Enzymology. 1992;211:353–88. doi: 10.1016/0076-6879(92)11020-J.

